# Acquired drug resistance enhances imidazoquinoline efflux by P-glycoprotein

**DOI:** 10.1101/2021.05.11.443528

**Authors:** Anunay J. Pulukuri, Anthony J. Burt, Larissa K. Opp, Colin M. McDowell, Amy E. Nielsen, Rock J. Mancini

## Abstract

Multidrug-Resistant (MDR) cancers mitigate the action of chemotherapeutics through drug efflux that occurs via ABC (ATP-Binding Cassette) transporters, including P-glycoprotein 1 (P-gp or ABCB1/*MDR1*). Because Toll-Like Receptor (TLR) agonist immunotherapies elicit abscopal anti-tumoral effects by modulating the activity of bystander tumor infiltrating immune cells, they not only circumvent the neutralizing effects of drug efflux, but could also work in synergy with this process. However, the effect of drug resistance on TLR agonist efflux is largely unknown. We begin to address this by investigating P-gp mediated efflux of model TLR agonists in cancer cell lines before and after acquired drug resistance. First, we used functionalized liposomes to determine that imidazoquinoline TLR agonists Imiquimod, Resiquimod, and Gardiquimod are substrates for P-gp. Next, we created Doxorubicin-resistant cancer cell lines from B16 melanoma, TRAMP prostate, and 4T1 breast cancer and observed that each cell line increased P-gp expression in response to Doxorubicin. Comparing imidazoquinoline efflux in Doxorubicin-resistant cell lines, relative to parent cancer cell lines, we used P-gp competitive substrates and inhibitors to demonstrate that imidazoquinoline efflux occurs through P-gp and is enhanced as a consequence of acquired drug resistance. We found that the most hydrophobic, yet least potent imidazoquinoline (Imiquimod), was the best substrate for efflux. This suggests a new parameter, susceptibility to drug efflux, could be an important consideration in the rationale design of the next generation of TLR agonist immunotherapies that are targeted to cancer cells, yet effect their mechanisms of action by modulating the activity of tumor infiltrating immune cells.

## Introduction

Multidrug resistance (MDR) directly correlates to over 90% of metastatic cancer deaths affecting a variety of blood cancers and solid tumors, including melanomas, breast, and prostate cancers alike (1). MDR cancers broadly enhance expression of proteins that promote drug efflux. A major mechanism of drug resistance, drug efflux enables active transport of chemotherapeutic drugs, from within a cancer cell to the extracellular space thereby lowering intracellular drug concentration and decreasing efficacy. (2).

The ABC (ATP-binding cassette) transporter superfamily, which consists of at least 48 members classified into different subfamilies based on sequence similarity, is the primary contributor to chemotherapeutic drug efflux (3,4). The first discovered (5,6) and most well studied ABC transporter that participates in drug efflux is P-glycoprotein (P-gp or ABCB1/*MDR1*) (7,8), and increased expression of P-gp is well established to promote cancer MDR (9). P-gp, along with other MDR-associated efflux proteins, exhibit a broad substrate scope capable of transporting entire classes of chemotherapeutics such as the taxanes or anthracyclines (10,11). P-gp is particularly promiscuous; it has been shown to transport diverse compounds with minimal correlation to chemical structure other than a weak association with hydrophobicity (12–16). Depending on the specific compound, the mechanism of P-gp-mediated drug efflux either involves release, as a consequence of decreased binding affinity caused by changes in specific residue contacts between the protein itself, or is facilitated by ATP binding and hydrolysis, which activate and reset the efflux action of P-gp, respectively (17).

Circumventing this process, several immunotherapeutics have emerged that appear to have efficacy fundamentally orthogonal to P-gp mediated drug resistance. Historically, agonists of highly conserved pattern recognition receptors, including Toll-Like Receptor (TLR) 4, contained in bacterial cell lysates, were found to generate anti-cancer immune responses (18,19). More recently, imidazoquinolines (TLR 7/8 agonists) have been reported to elicit anti-cancer properties by modulating tumor infiltrating immune cell populations (20). Imidazoquinolines promote activation of T, NK, and antigen-presenting cells, inhibit regulatory T cell function, and reduce myeloid derived suppressor cell populations, thereby promoting a myriad of immune-mediated anti-cancer effects (21–26). In particular, intratumoral administration of imidazoquinoline immunotherapeutics, which *presumably* undergo drug efflux, have demonstrated efficacy across a range of cancers both as mono (27) and combination (28) immunotherapies. This has led our group (29–31) and others (32–35) to develop drug delivery strategies that liberate imidazoquinolines within cancer cells or the tumor microenvironment. As such, knowing a pathway for imidazoquinoline efflux, and how MDR affects efflux (**Figure 1**), would be valuable in designing the next generation of TLR agonists targeted to MDR cancers.

**Figure 1:**
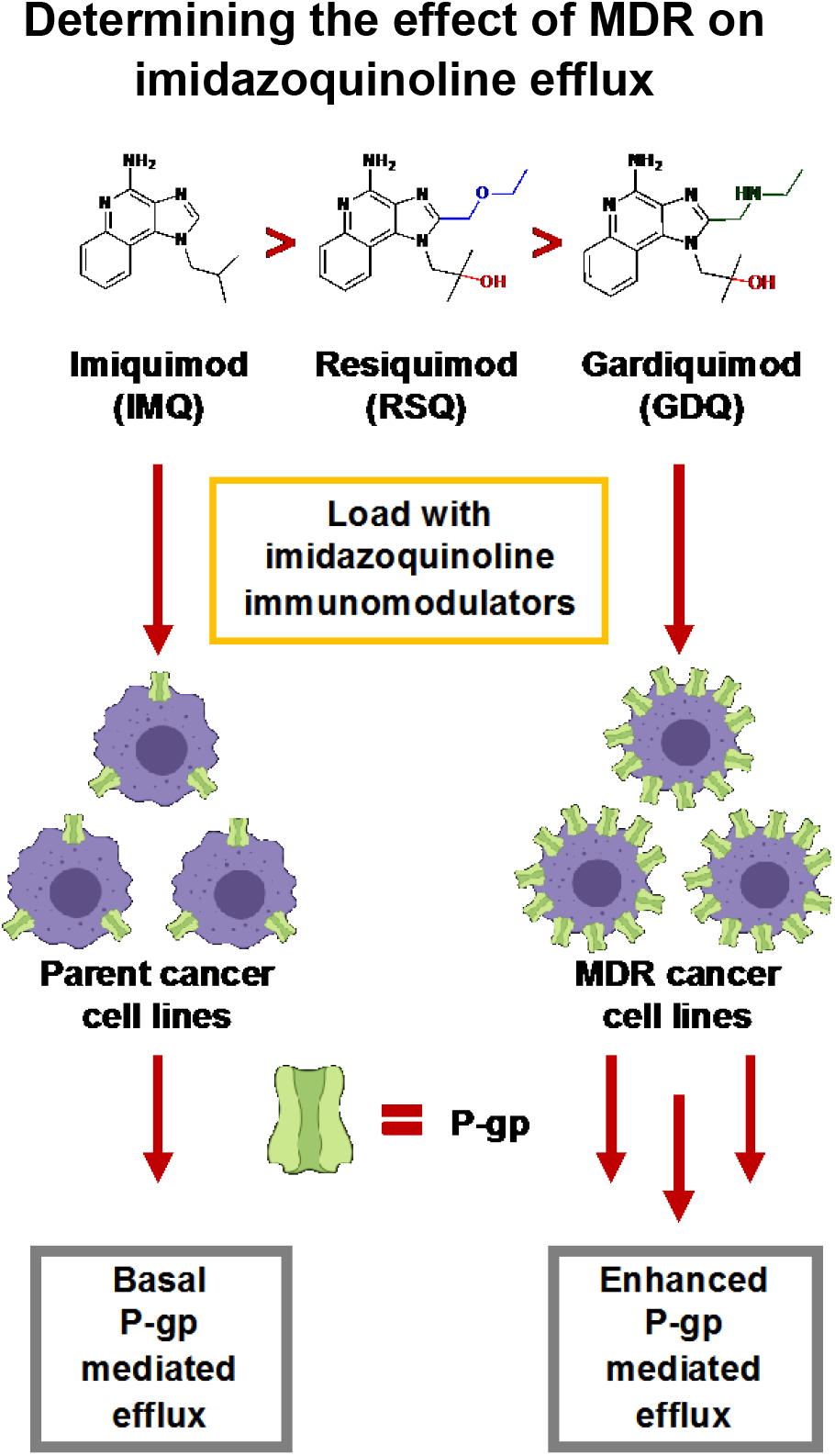
A schematic representation of our examination of P-gp-mediated efflux in parent cancer cell lines compared to Multidrug-resistant cell lines derived from parent cell lines that overexpress P-gp. We demonstrate that the imidazoquinolines Imiquimod (IMQ), Resiquimod, (RSQ) and Gardiquimod (GDQ) are substrates for P-gp. We also conclude that minimal efflux can also occur through other transport protein mediated efflux (active transport) or diffusion into the extracellular environment (passive transport).

However, pathways for imidazoquinoline efflux from MDR cancers is currently unproven. Herein, demonstrate that imidazoquinolines are substrates for P-gp and compare imidazoquinoline efflux in MDR cancer cells that overexpress P-gp alongside their non-MDR counterparts. Overall, we find efflux is significantly enhanced by the MDR phenotype, depending on both the type of cancer and the structure of the imidazoquinoline itself.

## Materials and Methods

*See Supporting Information for routine synthetic and cell culture procedures*.

### MDR Derived Cancer cells

The B16-F10 Melanoma (B16), TRAMP C-2 prostate (TC2), and 4T1-Luc2 breast (4T1) parent cancer cell lines were seeded at 3 × 10^5^ cells in T-175 culture flasks separate from the parent cancer cell lines. These cells were cultured in media which contained Doxorubicin (Dox). The original media was composed of: DMEM with 4.5 g L^-1^ glucose, 2 mM L-glutamine, 100 U mL^-1^ PenStrep, 10% HI-FBS and 1 nM Dox. The media was changed every 3-4 days until the cells reached 80% confluence. Passaging included changing media, counting, and seeding 3 × 10^5^ cells in 35 mL of new complete media in a new T-175 culture flask. Dox concentration was doubled with each passage until reaching a final concentration of 1 μM (**Figure 2)**.

**Figure 2:**
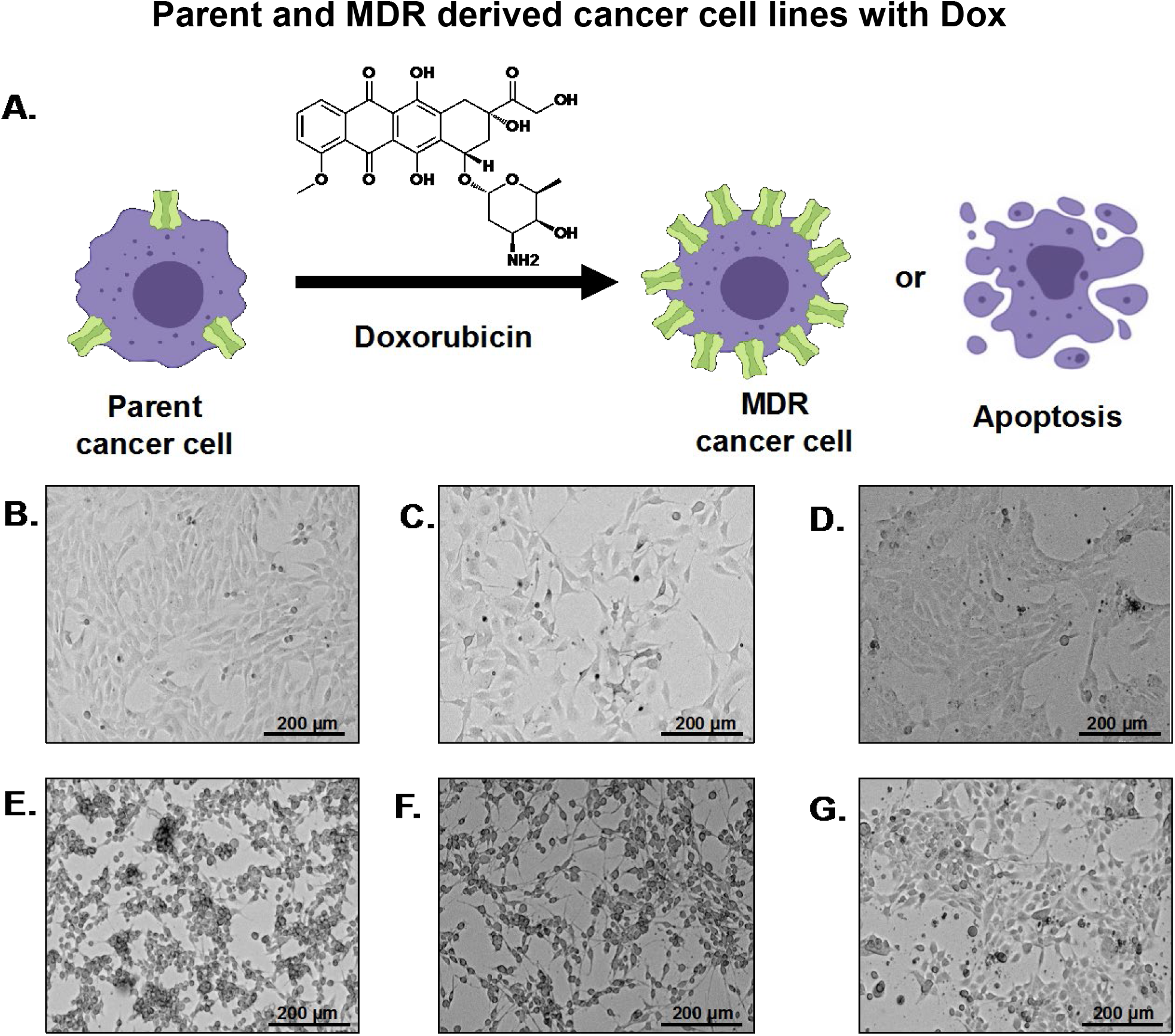
**A)** The MDR-derived cancer cell lines were derived from the parent cancer cell lines with increasing concentrations of Doxorubicin (Dox). The epigenetic pressure from Dox results in increased expression of transport proteins, thereby avoiding apoptosis. **B)** TC2 prostate parent cancer cell line. **C)** B16 melanoma parent cancer cell line. **D)** 4T1 breast parent cancer cell line. **E)** TC2–MDR derived cancer cell line. **F)** B16– MDR derived cancer cell line. **G)** 4T1–MDR derived cancer cell line. Scale Bar 200 μm

### Western Blot

Cell lysates were extracted using Triton X-100 lysing buffer and lysate was quantified using Pierce BCA Protein Assay Kit (ThermoFisher Scientific – 23225). 20 μg of protein for each cell lysate was run on a 4-15% SDS gel (Bio-Rad – 4561083DC) and electrotransferred onto a PVDF membrane. The membrane was washed with TBS and blocked overnight with 3% BSA in TBST. The membrane was incubated with primary Rabbit anti-P-gp antibody (Abcam – ab170904) for 2 h, washed 2x for 10 min with TBST, and incubated with a secondary Goat anti-Rabbit IgG H&L (Abcam – ab97051) antibody for 1 h. Mouse brain tissue lysate (Abcam – ab7190) and Rabbit anti-β-actin antibody (Abcam – ab8227) along with the MDR AT3B-1 cell line were used as controls (**Figure 3A**).

**Figure 3:**
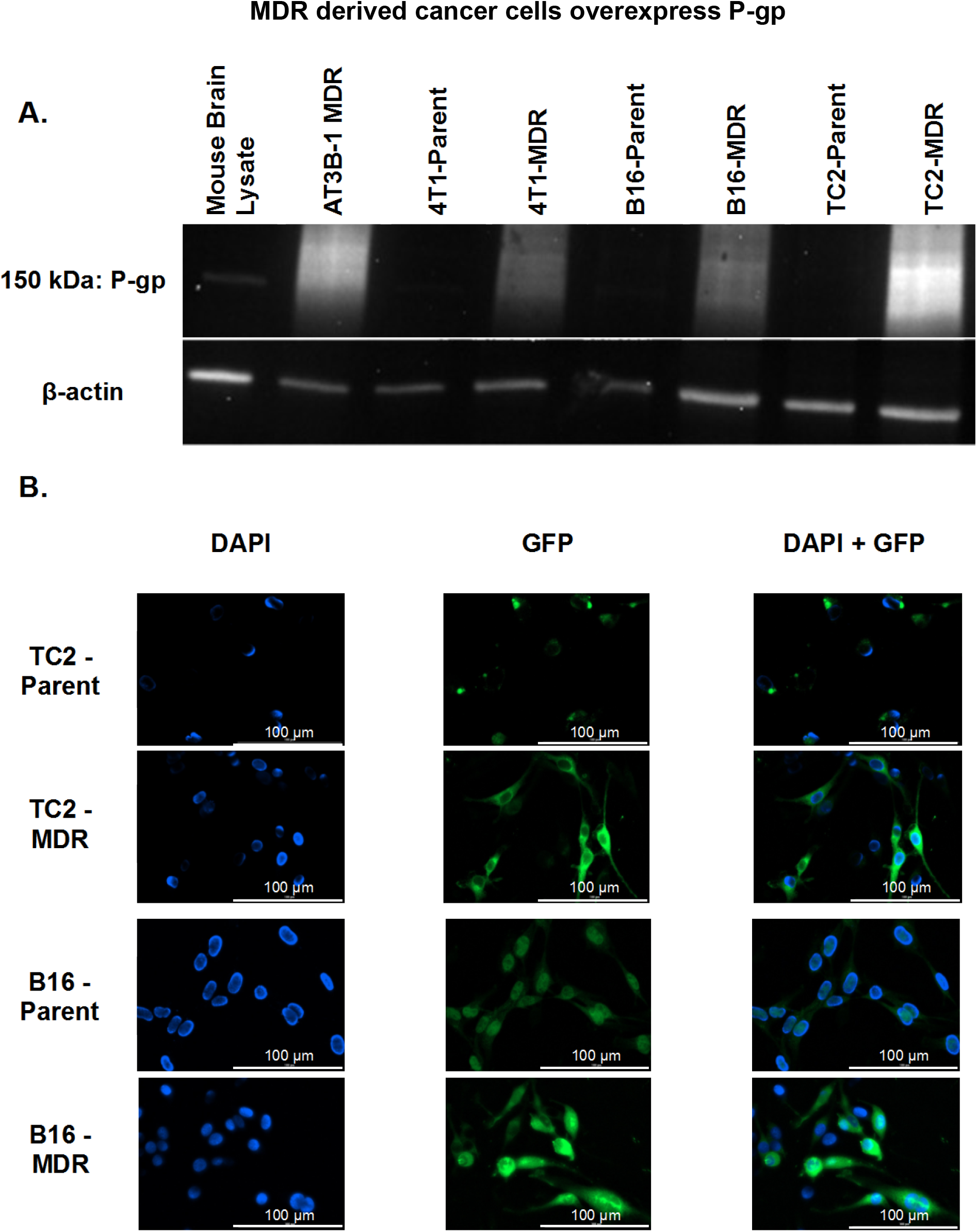
MDR-derived cancer cell lines express more P-gp when compared to the parent cancer cell lines. **A)** Western Blot shows the increased expression of P-gp in MDR-derived cancer cell lines when compared to the parent cancer cell lines. **B)** TC2-Parent, TC2-MDR, B16-Parent, and B16-MDR stained with Anti-MDR1/ABCB1 antibody conjugated to Alexa Fluor 488 not only shows the increased expression of P-gp in MDR-derived cancer cell lines, but also the localization of P-gp. Note: The 4T1 Parent and MDR-derived cell lines were not used in this study due to the compatibility between the fluorescent conjugated antibody and GFP expression linked to the Luc2 gene. Scale: 100 µm

### Immunofluorescence Assay

Cancer cells (Parent and MDR-derived) were plated on glass coverslips (neuVitro - GG-25-1.5-pdl) in a 6 well plate and allowed to reach 70% confluency. The cells were fixed using 4% formaldehyde for 15 min at room temperature. After three washes with 1 mL PBS, the cells were blocked (10% HI-FBS in PBS) for 1 h at 37 °C. Cells were incubated with Mouse anti-MDR1/ABCB1 antibody conjugated to Alexa Fluor 488 (Santa Cruz Biotechnology - sc55510 AF488) overnight in the dark at 4 °C before incubation with Mouse IgG Fc binding protein conjugated to CruzFluor 488 (Santa Cruz Biotechnology-sc533653) in the dark at room temperature for 1 h. Finally, coverslips were washed in PBS and mounted with Vectashield antifade mounting medium with DAPI (Vector Labs – H-2000-10) diluted in Vectashield antifade mounting media (Vector Labs – H-1900-10) to 0.1 μg mL^-1^. Fluorescent images (40x) were acquired on a Lionheart FX (BioTek Instruments) microscope (**Figure 3B**).

### ATPase Assay

Colorimetric measurement of imidazoquinoline interaction with P-gp was determined using a PREDEASY ATPase Assay Kit (SOLVO Biotechnology, Sigma-Aldrich) in 96 well plate format following the manufacturer’s protocol. Stock solutions of developer and blocker were diluted using Ultrapure DNase/RNase Free Distilled Water (Invitrogen – 10977015). Briefly, across individual wells, Imiquimod (IMQ) (eNovation Chemicals - SY017571), Resiquimod (RSQ) (Accel Pharmtech XP2356), Gardiquimod (GDQ) (synthesized in-house; see **SI** for detailed synthetic procedure and characterization) were added at 8 different concentrations (1-200 μM) to membrane vesicles expressing *hMDR1*. Each well contained 4 µg membrane protein and 1 µL of imidazoquinoline was added to arrive at the final concentration. The plate was pre-incubated (37 °C, 10 min) before 10 µL of MgATP solution was added to start the reaction. The plate was incubated (37°C, 10 min) before the ATPase reaction was quenched using 100 µL of Developer Solution at room temperature. After 2 min, 100 µL of Blocker solution was added to each well at room temperature before incubation (37°C, 10 min). Following incubation, the absorbance was measured using a microplate reader at 600 nm. Absorbance values were used to calculate liberated P_i_ (**Figure 4)**.

**Figure 4:**
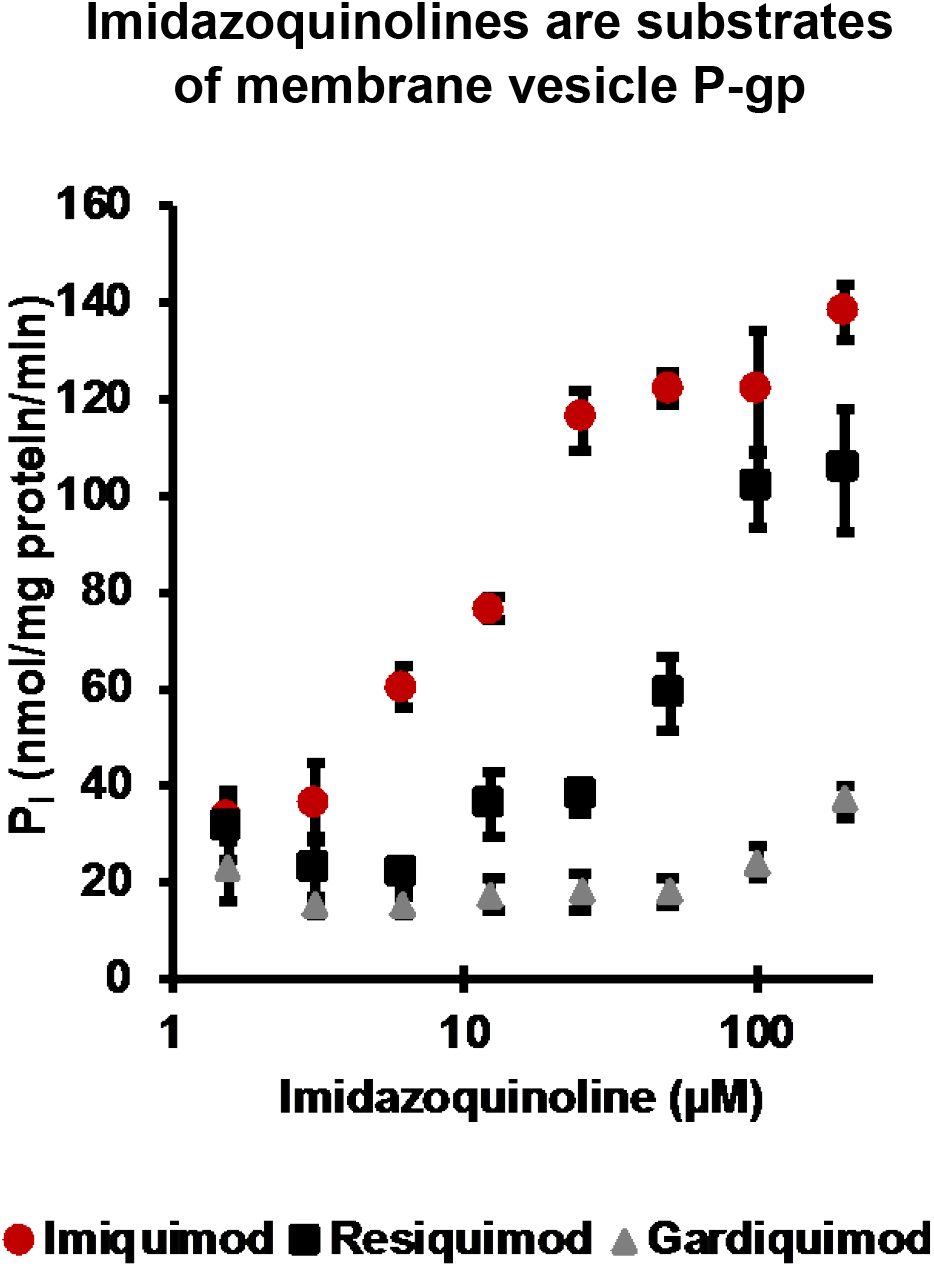
ATPase assay with IMQ, RSQ, and GDQ. In the activation test: IMQ, RSQ, and GDQ all stimulated vanadate sensitive ATPase activity above baseline confirming that the imidazoquinolines are substrates of P-gp. Error bars are standard deviation from the mean for experiments repeated in triplicate.

### Competitive Efflux studies with Rhodamine 123

Cancer cells (Parent and MDR-derived) were passaged, and 1 × 10^6^ cells were used for each compound tested with Rhodamine 123 (Rh123) (Cayman Chemical - 16672). Cells were suspended in DMEM, supplemented with 10% HI-FBS, and 1 μM Rh123; and incubated (37 °C, 30 min). Following incubation with Rh123, 1 or 10 μM of test compound (IMQ, RSQ, GDQ), and / or P-gp inhibitors Verapamil (VER) (Cayman Chemical - 14288) or Tariquidar (TQR) (Sigma Aldrich - SML 1790) were added to the cells and incubated for another 30 min at 37 °C. After incubation, samples were centrifuged (300 RCF, 5 min) and the supernatant discarded. Cells were fixed with 4% formaldehyde for 15 min at room temperature. The samples were centrifuged (300 RCF, 5 min) and suspended in 1 mL cold FACS buffer. Finally, the samples were loaded onto the Flow Cytometer (BD Accuri C6 Plus) to measure the Mean Fluorescence Intensity of Rh123. Results are representative of triplicate experiments and normalized to MFI for 1 μM Rh123 (**Figure 5**).

**Figure 5:**
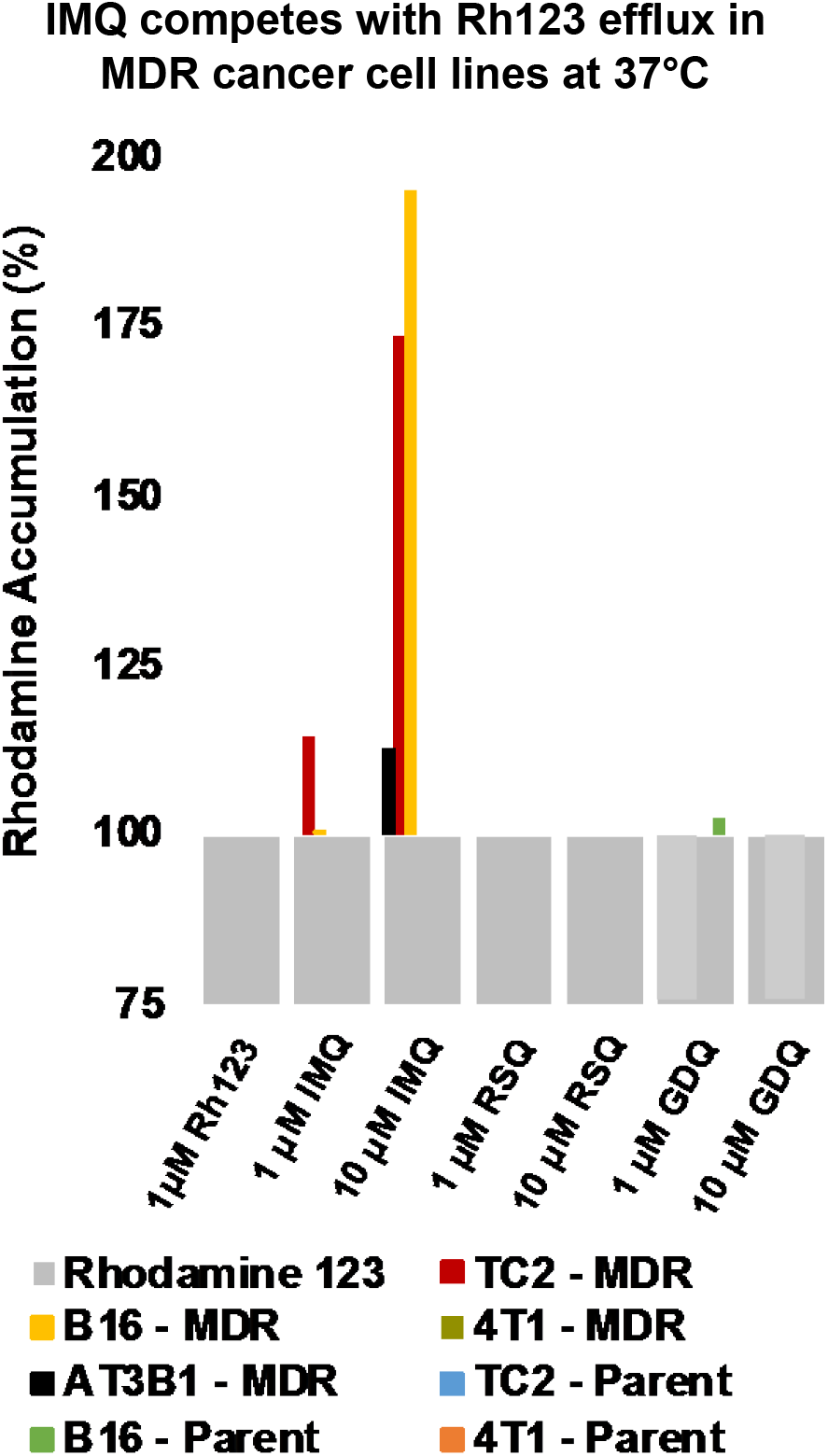
Competitive experiments performed at 37 °C with a known fluorescent substrate of P-gp, Rhodamine 123 (Rh123). IMQ leads to an increase in Rh123 accumulation in MDR-derived cancer cells under active transport conditions. Data representative of experiments repeated in triplicate.

### Uptake Studies with Rhodamine 123

Cancer cells (Parent and MDR-derived) were passaged, and 1 × 10^6^ cells were used for each compound being tested with Rh123. Cancer cells were suspended in DMEM, supplemented with 10% HI-FBS, and 1 μM Rh123. The cells were incubated (4 °C, 30 min) followed by adding IMQ, RSQ, GDQ, VER, or TQR (1 or 10 μM) and incubating further for 30 min at 4 °C. Next, samples were centrifuged (300 RCF, 5 min) and the supernatant discarded. Cells were fixed with 4% formaldehyde for 15 min at room temperature. The samples were centrifuged again (300 RCF, 5 min), suspended in 1 mL cold FACS buffer, and analyzed in triplicate for Rh123 MFI using the flow cytometer. Samples were normalized to uptake of 1 μM Rh123 as indicated in (**Figure 6**).

**Figure 6:**
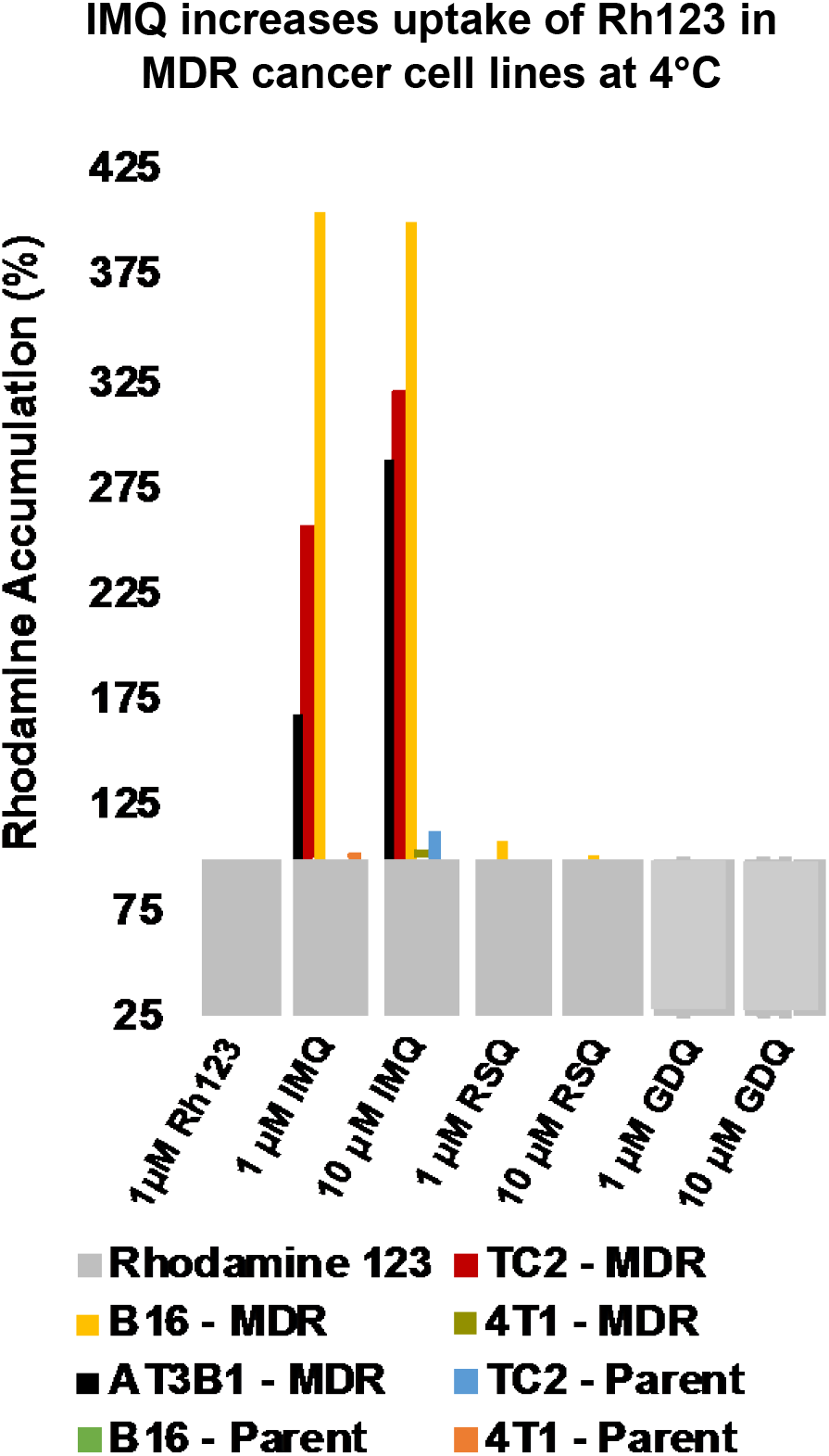
Uptake experiments performed at 4 °C with a known fluorescent substrate of P-gp, Rhodamine 123 (Rh123). IMQ leads to an increase in Rh123 accumulation in MDR-derived cancer cells under passive diffusion conditions. Data representative of experiments repeated in triplicate.

### Cellular Uptake Studies

Cancer cells (Parent and MDR-derived) were passaged, and 5 × 10^6^ cells were used for each experiment. Cells were suspended in 1 mL DMEM, supplemented with 10% HI-FBS, with 100 μM of IMQ, RSQ, or GDQ and incubated at 4 °C for 30 min. Samples were then centrifuged (300 RCF, 5 min) and washed 2 times with 1 mL cold PBS. The samples were re-suspended in lysing buffer: For IMQ and RSQ, lysing buffer consisted of: 40% HPLC Grade acetonitrile (ACN) in HPLC Grade water (H_2_O) with 0.1 % Trifluoroacetic acid (TFA) and 1% v/v Trition X-100. Due to solubility differences, GDQ lysing buffer consisted of HPLC Grade water with 0.1% TFA and 1% v/v Triton X-100. Cells were lysed on ice for 20 min. Following the lysing step, samples were centrifuged (12500 RCF, 10 min), supernatant collected, and filtered using a 0.2 μm PTFE filter. Filtered samples were analyzed by HPLC (Thermo Dionex UltiMate 3000 running Chromeleon software V6.80 SR14) with a C18 analytical column (Phenomenex XB-C18 100Å, 250 x 4.6 mm, 5µm) at a flow rate of 1 mL/min, A: H_2_O with 0.1% TFA, B: ACN with 0.1% TFA, with UV detection at 254 nm. For IMQ and RSQ an isocratic method (40% B) was used. For GDQ, a gradient method (10% B for 5 min, 10 to 95% B over 14 min, 95% B for 10 min) was used. 9-point calibration curves were derived for each of the imidazoquinolines using standards with concentrations ranging from 500 nM to 250 μM. The calibration curves were used to fit using linear regression for each imidazoquinoline: IMQ (y = 0.039x; R^2^ = 0.9976), RSQ (y = 0.0447x; R^2^ = 0.9994), and GDQ (y = 0.0271x; R^2^ = 0.9996). All samples were analyzed in triplicate. Cellular uptake ratio was obtained using the following equation (**Figure 7)**:

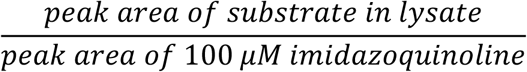

**Figure 7:**
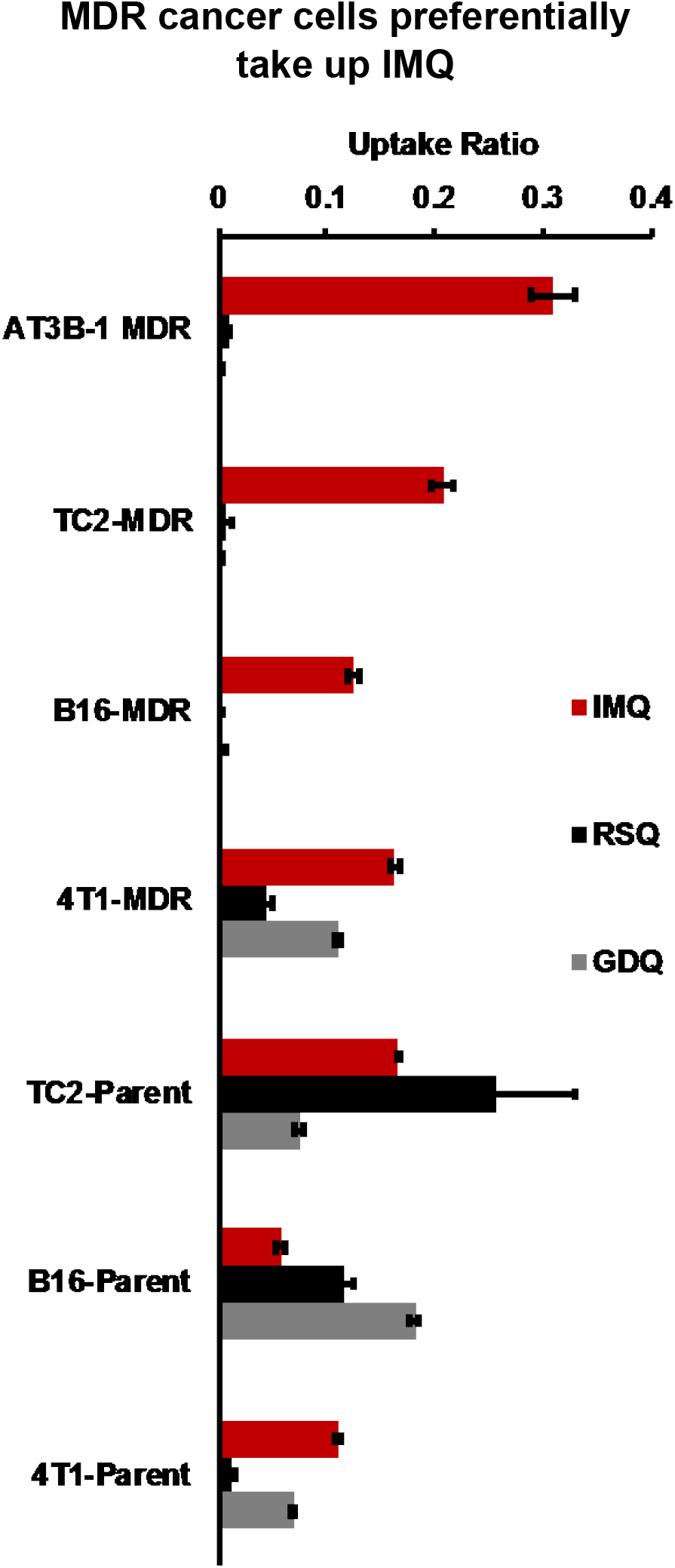
Uptake Experiments performed with an excess concentration (100 μM) of IMQ, RSQ, and GDQ under passive diffusion conditions in parent and MDR-derived cancer cell lines. Data representative of experiments performed in triplicate.

### Cytotoxicity Assay

Resazurin Cell Viability Assay kit (Abcam - ab129732) was used to determine if the concentrations of IMQ, RSQ, and GDQ used in experiments were cytotoxic to the cancer cell lines. To begin, cancer cells were plated in complete cell media (DMEM, 10% HI-FBS) in 96-well plates, with two rows per plate of each density/well tested: 2000 cells/well, 25,000 cells/well, 50,000 cells/well, and 100,000 cells/well. Cells were treated with 20 uL of 1 mM solutions of IMQ, RSQ, and GDQ, to give a final concentration per well of 100 uM for each compound. Plates were incubated at 37 °C for 1 h. After incubation, 10 uL of resazurin stain was added to wells in alternating rows on the plate, such that for each density and compound tested in triplicate there was a corresponding blank without stain added. Plates were incubated further with absorbance measured at 570 nm and 600 nm at 1, 2, 3, 4, and 24 h intervals. The absorbance measurements at 600 nm, which correlated to the absorbance of resazurin, were subtracted from those taken at 570 nm, correlating to the resorufin absorbance. After this, values were normalized to cell viability of the negative control. Experiments were performed in triplicate and the 3 h time-point for 5×10^5^ cells is shown (**Figure S4**).

## Results

### MDR cancer cell lines overexpress P-gp

MDR cancer cell lines were derived from parent cancer cell lines by introducing increasing concentrations of Dox until cell populations capable of stable proliferation in 1 µM Dox were obtained (**Figure 2**). To evaluate P-gp expression, a western blot and immunofluorescence assay were performed (**Figure 3**). These experiments showed that MDR-derived cell lines expressed more P-gp than their parent cancer cell line counterparts, with expression levels comparable to the MDR AT3B-1 positive control cell line (**Figure 3A**). Localization of P-gp expression with punctate accumulation on cellular membranes was subsequently confirmed by fluorescence microscopy (**Figure 3B**).

### Imiquimod, Resiquimod, and Gardiquimod are substrates of P-gp

To determine if imidazoquinolines are substrates of P-gp, an ATPase assay, using purified membrane vesicles expressing P-gp (*hMDR1*) was performed (**Figure 4**). All 3 imidazoquinolines were tested at the same concentrations from 1.56 to 200 µM. IMQ P-gp activity (quantified as liberated Pi) was detectable even at low concentrations (6.25 µM) with increased activity observed at increasing concentrations. RSQ also stimulated P-gp at moderately higher concentrations and GDQ was a poor substrate by comparison, only liberating P_i_ above baseline at the highest concentrations tested. Imidazoquinolines did not inhibit maximal vanadate sensitive ATPase activity when tested as inhibitors (**Figure S1**). Overall, all three imidazoquinolines liberated P_i_ in the activation test, confirming that they are substrates of P-gp to varying degrees; based upon this result, we concluded imidazoquinolines are P-gp substrates with order of affinity IMQ > RSQ > GDQ.

### Imiquimod competes with Rhodamine 123 efflux in MDR-derived cancer cell lines

To determine whether the imidazoquinolines compete with a known fluorescent P-gp substrate, Rhodamine 123 (Rh123), competitive efflux studies were performed at 37 °C in both parent and MDR-derived cancer cell lines. Significant increases in MFI were observed for Rh123 *and* IMQ for AT3B-1-MDR, TC2-MDR, and B16-MDR (**Figure 5**) as compared to the MFI of Rh123 alone. In all MDR samples, MFI also increased in the presence of the P-gp substrate & inhibitor VER (36,37), as well as TQR, a third generation inhibitor of P-gp (38) (**Figure S2**). Interestingly, there was no significant increase observed in the presence of RSQ and GDQ. Furthermore, there was no significant increase in MFI observed in the presence of any of the imidazoquinolines for the 4T1-MDR and all parent cancer cell lines (**Figure 5**). Additionally, there was no increase in fluorescence in the presence of either VER or TQR in all of the parent cancer cell lines. This result agreed with our experiment determining the expression of P-gp in the MDR cell lines. Overall, based on the results from competitive experiments in parent and MDR cancer cell lines, we concluded that IMQ competes with Rh123 only in MDR-derived cancer cell lines with the 4T1-MDR cell line as the lone exception. Although we did not observe significant direct competition with RSQ or GDQ, we hypothesized this could be due to poor loading of these imidazoquinolines.

### Imiquimod increases Rhodamine 123 uptake under passive diffusion conditions

The competitive efflux experiments performed at 37 °C confirmed that IMQ competes with a fluorescent P-gp substrate in most of the MDR-derived cancer cell lines, but not in parent cancer cell lines. However, this trend was not observed with RSQ or GDQ in either parent or MDR-derived cancer cell lines. In order to determine if changing the loading conditions would impact the results observed in the competitive efflux experiments, uptake studies with Rh123 were performed under passive diffusion conditions at 4 °C. Under these conditions, we anticipated that P-gp would be inactive. As shown in **Figure 6**, no significant increases in Rh123 accumulation were observed in the presence of RSQ and GDQ in all cell lines tested, relative to Rh123 alone. Interestingly, significant increases in MFI were observed in the presence of IMQ. VER exhibited the same trend as IMQ as did TQR, though to a lesser extent (**Figure S3**). This trend was only observed in the same MDR-derived cell lines similar to the competitive experiments performed at 37 °C. This suggested that IMQ facilitates the uptake of Rh123 under passive diffusion conditions, similar to VER.

### Resiquimod and Gardiquimod are not taken up by MDR derived cancer cells

To confirm our observations, from the uptake and competitive experiments with Rh123, that RSQ and GDQ are not being taken up by the cancer cell lines, an uptake study was performed without the presence of fluorophore. Cancer cells were loaded with imidazoquinolines (100 µM), lysed, and analyzed via HPLC. Even at the relatively high concentration of 100 µM, RSQ and GDQ were not taken up by TC2-MDR, B16-MDR, or AT3B-1 cells (**Figure 7**). Upon calculation of the concentration of the imidazoquinolines in the lysates, we found that when 100 µM IMQ was loaded into the cancer cells, TC2-MDR (20.75 µM) had the highest detectable concentration compared to the other MDR derived cancer cell lines. In comparison, the imidazoquinolines were taken up by all parent cancer cell lines. Here, the TC2-Parent (16.58 µM) had the highest concentration of IMQ, followed by the 4T1-Parent (11.06 µM) and B16-Parent (5.72 µM) cell lines. We found that loading 100 µM RSQ also did lead to detectable concentrations, but only in parent cancer cell lines: TC2-Parent (25.84 µM), B16-Parent (11.65 µM), and 4T1-Parent (1.34 µM). GDQ (when loaded at 100 µM) was only detected in the parent cancer cell lines as well, with B16-Parent having the highest detectable concentration (16.42 µM). Interestingly, detectable amounts of IMQ (16.31 µM), RSQ (4.47 µM), and GDQ (10.13 µM) are taken up by 4T1-MDR derived cancer cell line; however, no competition is observed in the presence of Rh123 for RSQ and GDQ. Finally, because imidazoquinolines are also known to directly induce apoptosis in a variety of cancers independently of immunogenic effects (39), we also confirmed that the tested concentrations / incubation times of imidazoquinoline were not cytotoxic to the cancer cells during by performing a cytotoxicity assay. Here we found no difference in resazurin oxidative potential between all cultures with imidazoquinolines and untreated controls **(Figure S4)**.

## Discussion

Multidrug resistance is a major challenge for traditional chemotherapy. This characteristic is attributed to many different mechanisms such as enhanced drug efflux, increased DNA damage repair, reduced apoptosis, and/or drug metabolism (40,41). Melanoma, Prostate, and Breast cancers are predominantly the most common types of cancers and are presently in the top 5 of estimated new cases in 2020 (42). Therefore, the cancer cell types chosen for this study were B16 melanoma, TRAMP prostate, and 4T1 breast cancer cell lines, as well as the AT3B-1 prostate cancer cell line which served as a positive control because it possessed the MDR phenotype which has been previously well-characterized (43). These cell lines were primarily chosen based on their ability to develop drug resistance (44–46), and we mimicked this in our parent cancer cell lines to achieve MDR cells which we expected (and confirmed) to overexpress P-gp.

Deriving the MDR cancer cell lines in this study began by inducing them with increasing concentrations of Dox from 1 nM to 1 µM. Each of the cancer cell lines reached the 1 µM Dox threshold at different times. The TC2-MDR version took 3 months compared to B16-MDR which took 7 months and the 4T1-MDR cells required well over 8 months to reach 1 µM Dox (see **SI** for specific timeline and additional microscope images). Interestingly, the morphology of the cancer cells was also observed to change, suggestive of the epithelial to mesenchymal transition that can occur upon chronic exposure to chemotherapeutics (47). Our MDR cancer cell lines showed a significant increase in the expression of P-gp when compared with their parent counterparts as measured by western blot (**Figure 3A**), and we confirmed the expected P-gp membrane localization via fluorescence microscopy (**Figure 3B**).

With increased P-gp expression in MDR cancer cell lines confirmed, we next investigated imidazoquinoline efflux through P-gp, both in membrane vesicles as well as in cell lines. The imidazoquinoline immunostimulants chosen for this study were: Imiquimod (IMQ), Resiquimod (RSQ), and Gardiquimod (GDQ). We chose these particular imidazoquinolines for their structural similarity and relevance in cancer immunotherapies. In particular the TLR7 agonist IMQ is FDA approved for a variety of conditions and known to confer anticancer immunogenicity (48). RSQ, a more potent TLR 7 and 8 dual agonist, features nanomolar potency, and is capable of activating TLR 8 in humans which is expressed by myeloid-derived dendritic cells, an advantage when compared to IMQ (49). GDQ, a TLR 7 agonist, is both more potent than IMQ, and also exerts enhanced antitumor effects (50).

Previously, our own work suggested that IMQ and RSQ undergo drug efflux from a range of MDR or parent cancer cell lines (30,31) and in the present study, we confirm that IMQ, RSQ, and GDQ are in fact substrates of P-gp using an ATPase membrane transport study (**Figure 4**). We also conclude that, although less potent, IMQ is a better substrate when compared to RSQ and GDQ in most of the MDR-derived cancer cell lines.

IMQ competed with Rh123 for efflux in most of the MDR-cancer cell lines, a trend not observed in the parent cancer cell lines (**Figures 5**). This result directly correlated with the observed overexpression of P-gp in the MDR-derived cancer cell lines and is consistent with our previous report that IMQ competes with Rh123 for efflux in MDR AT3B-1 cells (30). This also correlates with the enhanced efflux potential of our MDR cell lines, driven in-part by P-gp. Alternatively, we were able to conclude from the uptake experiments with (**Figure 6**) and without Rh123 (**Figure 7**), that we do not observe an increase in MFI with RSQ and GDQ because they are not taken up into the MDR cell lines.

Based upon these results, we can conclude that hydrophobicity / susceptibility to P-gp efflux should be considered, alongside potency, when choosing the optimal substrate for use in immunotherapeutics delivered to MDR cancers for the purposes of activating tumor infiltrating immune cells. Because cLogP can influence P-gp efflux and does not negatively impact passive permeability unless the values fall outside acceptable oral drug-like ranges (cLogP < 1 or cLogP ≥ 7),(11) the predicted cLogP value (ChemDraw 19.1 Software – PerkinElmer Informatics) for IMQ (cLogP = 1.428) falling within this range with values of RSQ (cLogP = 0.036) and GDQ (cLogP = - 0.254) falling outside could explain the lack of uptake by MDR cancers.

It is important to note that even though expression of P-gp was increased in the 4T1-MDR cell line relative to the non-MDR parent line, we did not observe imidazoquinoline competition with Rh123. Thus, while this study demonstrates imidazoquinolines, particularly IMQ, are substrates for P-gp mediated efflux, it is also likely that imidazoquinolines could also serve as substrates for some of the many other ABC transporters. Although some generalizable differences do exist (MRP1 mostly transports organic anions and shares only 23% sequence identity with P-gp), there is also significant overlap between the substrate scopes of efflux proteins associated with MDR that further complicate both development and analysis of efflux (51–53). It is also possible that Dox MDR provokes overexpression of other ABC transporters or other ABC independent mechanisms. To address these variables, future studies will characterize our MDR cancer cell lines at the genomic level to identify the genetic response to Dox, in addition to the P-gp expression reported here.

In conclusion, we used Dox to derive MDR-versions of B16, TC2, and 4T1 cells; and characterized them for enhanced P-gp expression. We also demonstrated that the imidazoquinoline immunostimulants IMQ, RSQ, and GDQ are substrates of P-gp and that, at a minimum, P-gp enhances IMQ efflux in the MDR cancer cell lines. Additionally, we were able to show that these imidazoquinolines are subject to an increased efflux potential correlated to the expression of P-gp based on our findings in the competitive experiments with Rh123. We envision these results will enable better understanding of immunostimulant trafficking, particularly that of the imidazoquinolines, ultimately contributing to the development of new cancer immunotherapies that could be enhanced by the mechanisms of drug efflux.

## Acknowledgements

Research reported in this publication was supported by the National Cancer Institute of the National Institutes of Health under Award Number 1R01CA234115. The content is solely the responsibility of the authors and does not necessarily represent the official views of the National Institutes of Health. The authors thank Professor Katrina Mealey for expert advice with efflux experiments and Professor Darrell Irvine for donating the 4T1 cells. The authors also thank Dr. Maryam Davaritouchaee for assisting with preliminary cell uptake experiments. NMR characterization of GDQ and synthetic intermediates was made possible through use of the Washington State University NMR Center with equipment supported by NIH RR0631401, RR12948, NSF CHE-9115282, and DBI-9604689, the Murdock Charitable Trust, and private donors Don and Marianna Matteson. Figures 1 and 2 created with BioRender.com.

## Author Contributions

AJP derived MDR cell lines, conducted uptake and competitive efflux experiments, and performed the immunofluorescence assays. AJB synthesized GDQ and assisted with uptake experiments. LKO performed the western blot and assisted with cell culture and immunofluorescence. CMM conducted the ATPase assay. AEN conducted the cytotoxicity assays. RJM conceived of the project and directed the work. All authors contributed to preparation of the manuscript.

## Supporting Information

Additional figures, flow cytometry plots, microscope images, and synthetic procedure and characterization of GDQ can be found in the supporting information.

## References

1. Longley DB, Johnston PG. Molecular mechanisms of drug resistance. The Journal of Pathology. 2005;205(2):275–92.

2. Szakács G, Paterson JK, Ludwig JA, Booth-Genthe C, Gottesman MM. Targeting multidrug resistance in cancer. Nat Rev Drug Discov. 2006 Mar;5(3):219–34.

3. Glavinas H, Krajcsi P, Cserepes J, Sarkadi B. The Role of ABC Transporters in Drug Resistance, Metabolism and Toxicity. CDD. 2004 Jan 1;1(1):27–42.

4. Borst P, Elferink RO. Mammalian ABC Transporters in Health and Disease. Annu Rev Biochem. 2002 Jun;71(1):537–92.

5. Juliano RL, Ling V. A surface glycoprotein modulating drug permeability in Chinese hamster ovary cell mutants. Biochimica et Biophysica Acta (BBA) - Biomembranes. 1976 Nov;455(1):152–62.

6. Riordan JR, Ling V. Purification of Pglycoprotein from plasma membrane vesicles of Chinese hamster ovary cell mutants with reduced colchicine permeability. J Biol Chem. 1979 Dec 25;254(24):12701–5.

7. Juranka PF, Zastawny RL, Ling V. Pglycoprotein: multidrug-resistance and a superfamily of membrane-associated transport proteins. FASEB J. 1989 Dec;3(14):2583–92.

8. Ferreira RJ, dos Santos DJ, Ferreira M-JU. P-glycoprotein and membrane roles in multidrug resistance. Future Medicinal Chemistry. 2015 Jun;7(7):929–46.

9. Gottesman MM, Ling V. The molecular basis of multidrug resistance in cancer: The early years of P-glycoprotein research. FEBS Letters. 2006 Feb 13;580(4):998– 1009.

10. Matsson P, Pedersen JM, Norinder U, Bergström CAS, Artursson P. Identification of Novel Specific and General Inhibitors of the Three Major Human ATP-Binding Cassette Transporters P-gp, BCRP and MRP2 Among Registered Drugs. Pharm Res. 2009 Aug;26(8):1816–31.

11. Hitchcock SA. Structural Modifications that Alter the P-Glycoprotein Efflux Properties of Compounds. J Med Chem. 2012 Jun 14;55(11):4877–95.

12. Ueda K, Cardarelli C, Gottesman MM, Pastan I. Expression of a full-length cDNA for the human “MDR1” gene confers resistance to colchicine, doxorubicin, and vinblastine. Proceedings of the National Academy of Sciences. 1987 May 1;84(9):3004–8.

13. Gottesman MM. Mechanisms of Cancer Drug Resistance. Annu Rev Med. 2002 Feb;53(1):615–27.

14. Gatlik-Landwojtowicz E, Äänismaa P, Seelig A. Quantification and Characterization of PGlycoprotein−Substrate Interactions. Biochemistry. 2006 Mar 1;45(9):3020–32.

15. Ramachandra M, Ambudkar SV, Chen D, Hrycyna CA, Dey S, Gottesman MM, et al. Human P-Glycoprotein Exhibits Reduced Affinity for Substrates during a Catalytic Transition State †. Biochemistry. 1998 Apr;37(14):5010–9.

16. Pawagi AB, Wang J, Silverman M, Reithmeier RAF, Deber CM. Transmembrane Aromatic Amino Acid Distribution in P-glycoprotein. Journal of Molecular Biology. 1994 Jan;235(2):554– 64.

17. Aller SG, Yu J, Ward A, Weng Y, Chittaboina S, Zhuo R, et al. Structure of PGlycoprotein Reveals a Molecular Basis for Poly-Specific Drug Binding. Science. 2009 Mar 27;323(5922):1718–22.

18. Coley WB. II. Contribution to the Knowledge of Sarcoma. Ann Surg. 1891 Sep;14(3):199–220.

19. Coley WB. The Treatment of Inoperable Sarcoma by Bacterial Toxins (the Mixed Toxins of the Streptococcus erysipelas and the Bacillus prodigiosus). Proc R Soc Med. 1910;3(Surg Sect):1–48.

20. Smits ELJM, Ponsaerts P, Berneman ZN, Van Tendeloo VFI. The Use of TLR7 and TLR8 Ligands for the Enhancement of Cancer Immunotherapy. The Oncol. 2008 Aug;13(8):859–75.

21. Prins RM, Craft N, Bruhn KW, Khan-Farooqi H, Koya RC, Stripecke R, et al. The TLR-7 Agonist, Imiquimod, Enhances Dendritic Cell Survival and Promotes Tumor Antigen-Specific T Cell Priming: Relation to Central Nervous System Antitumor Immunity. J Immunol. 2006 Jan 1;176(1):157–64.

22. Caron G, Duluc D, Frémaux I, Jeannin P, David C, Gascan H, et al. Direct Stimulation of Human T Cells via TLR5 and TLR7/8: Flagellin and R-848 Up-Regulate Proliferation and IFN-γ Production by Memory CD4 + T Cells. J Immunol. 2005 Aug 1;175(3):1551–7.

23. Huang SJ, Hijnen D, Murphy GF, Kupper TS, Calarese AW, Mollet IG, et al. Imiquimod Enhances IFN-γ Production and Effector Function of T Cells Infiltrating Human Squamous Cell Carcinomas of the Skin. Journal of Investigative Dermatology. 2009 Nov;129(11):2676–85.

24. Hart OM, Athie-Morales V, O’Connor GM, Gardiner CM. TLR7/8-Mediated Activation of Human NK Cells Results in Accessory Cell-Dependent IFN-γ Production. J Immunol. 2005 Aug 1;175(3):1636–42.

25. Peng G. Toll-Like Receptor 8-Mediated Reversal of CD4+ Regulatory T Cell Function. Science. 2005 Aug 26;309(5739):1380–4.

26. Yin T, He S, Wang Y. Toll-like receptor 7/8 agonist, R848, exhibits antitumoral effects in a breast cancer model. Molecular Medicine Reports. 2015 Sep;12(3):3515–20.

27. Han J-H, Lee J, Jeon S-J, Choi E-S, Cho SD, Kim B-Y, et al. In vitro and in vivo growth inhibition of prostate cancer by the small molecule imiquimod. International Journal of Oncology. 2013 Jun;42(6):2087– 93.

28. Mullins SR, Vasilakos JP, Deschler K, Grigsby I, Gillis P, John J, et al. Intratumoral immunotherapy with TLR7/8 agonist MEDI9197 modulates the tumor microenvironment leading to enhanced activity when combined with other immunotherapies. j immunotherapy cancer. 2019 Dec;7(1):244.

29. Hantho JD, Strayer TA, Nielsen AE, Mancini RJ. An Enzyme-Directed Imidazoquinoline for Cancer Immunotherapy. ChemMedChem. 2016 Nov 21;11(22):2496–500.

30. Burt AJ, Hantho JD, Nielsen AE, Mancini RJ. An Enzyme-Directed Imidazoquinoline Activated by Drug Resistance. Biochemistry. 2018 Apr 17;57(15):2184–8.

31. Ryan AT, Pulukuri AJ, Davaritouchaee M, Abbasi A, Hendricksen AT, Opp LK, et al. Comparing the immunogenicity of glycosidase-directed resiquimod prodrugs mediated by cancer cell metabolism. Acta Pharmacol Sin. 2020 Jul;41(7):995–1004.

32. Li H, Van Herck S, Liu Y, Hao Y, Ding X, Nuhn L, et al. Imidazoquinoline-Conjugated Degradable Coacervate Conjugate for Local Cancer Immunotherapy. ACS Biomater Sci Eng. 2020 Sep 14;6(9):4993–5000.

33. Mottas I, Bekdemir A, Cereghetti A, Spagnuolo L, Yang Y-SS, Müller M, et al. Amphiphilic nanoparticle delivery enhances the anticancer efficacy of a TLR7 ligand via local immune activation. Biomaterials. 2019 Jan;190–191:111–20.

34. Nuhn L, De Koker S, Van Lint S, Zhong Z, Catani JP, Combes F, et al. Nanoparticle-Conjugate TLR7/8 Agonist Localized Immunotherapy Provokes Safe Antitumoral Responses. Adv Mater. 2018 Nov;30(45):1803397.

35. Yoo E, Salyer ACD, Brush MJH, Li Y, Trautman KL, Shukla NM, et al. Hyaluronic Acid Conjugates of TLR7/8 Agonists for Targeted Delivery to Secondary Lymphoid Tissue. Bioconjugate Chem. 2018 Aug 15;29(8):2741–54.

36. Ledwitch KV, Gibbs ME, Barnes RW, Roberts AG. Cooperativity between verapamil and ATP bound to the efflux transporter P-glycoprotein. Biochemical Pharmacology. 2016 Oct;118:96–108.

37. Durie BGM, Dalton WS. Reversal of drugresistance in multiple myeloma with verapamil. Br J Haematol. 1988 Feb;68(2):203–6.

38. Weidner LD, Fung KL, Kannan P, Moen JK, Kumar JS, Mulder J, et al. Tariquidar Is an Inhibitor and Not a Substrate of Human and Mouse P-glycoprotein. Drug Metabolism and Disposition. 2016 Jan 16;44(2):275–82.

39. Schön MP, Schön M. Immune modulation and apoptosis induction: Two sides of the antitumoral activity of imiquimod. Apoptosis. 2004 May;9(3):291–8.

40. Kachalaki S, Ebrahimi M, Mohamed Khosroshahi L, Mohammadinejad S, Baradaran B. Cancer chemoresistance; biochemical and molecular aspects: a brief overview. European Journal of Pharmaceutical Sciences. 2016 Jun;89:20– 30.

41. Salehan MR, Morse HR. DNA damage repair and tolerance: a role in chemotherapeutic drug resistance. British Journal of Biomedical Science. 2013 Jan;70(1):31–40.

42. Siegel RL, Miller KD, Jemal A. Cancer statistics, 2020. CA A Cancer J Clin. 2020 Jan;70(1):7–30.

43. Replogle-Schwab TS, Schwab ED, Pienta KJ. Development of doxorubicin resistant rat prostate cancer cell lines. Anticancer Res. 1997 Dec;17(6D):4535–8.

44. Mariani M, Supino R. Morphological alterations induced by doxorubicin in B16 melanoma cells. Cancer Letters. 1990 Jun 15;51(3):209–12.

45. Foster BA, Gingrich JR, Kwon ED, Madias C, Greenberg NM. Characterization of Prostatic Epithelial Cell Lines Derived from Transgenic Adenocarcinoma of the Mouse Prostate (TRAMP) Model. Cancer Res. 1997 Aug 15;57(16):3325.

46. Bao L, Haque A, Jackson K, Hazari S, Moroz K, Jetly R, et al. Increased Expression of P-Glycoprotein Is Associated with Doxorubicin Chemoresistance in the Metastatic 4T1 Breast Cancer Model. The American Journal of Pathology. 2011 Feb;178(2):838–52.

47. Thiery JP, Acloque H, Huang RYJ, Nieto MA. Epithelial-Mesenchymal Transitions in Development and Disease. Cell. 2009 Nov;139(5):871–90.

48. Schon M, Schon M. The Antitumoral Mode of Action of Imiquimod and Other Imidazoquinolines. CMC. 2007 Mar 1;14(6):681–7.

49. Jurk M, Heil F, Vollmer J, Schetter C, Krieg AM, Wagner H, et al. Human TLR7 or TLR8 independently confer responsiveness to the antiviral compound R-848. Nat Immunol. 2002 Jun;3(6):499–499.

50. Ma F, Zhang J, Zhang J, Zhang C. The TLR7 agonists imiquimod and gardiquimod improve DC-based immunotherapy for melanoma in mice. Cell Mol Immunol. 2010 Sep;7(5):381–8.

51. Johnson ZL, Chen J. Structural Basis of Substrate Recognition by the Multidrug Resistance Protein MRP1. Cell. 2017 Mar;168(6):1075-1085.e9.

52. Orlando BJ, Liao M. ABCG2 transports anticancer drugs via a closed-to-open switch. Nat Commun. 2020 Dec;11(1):2264.

53. Rosenberg MF, Bikadi Z, Hazai E, Starborg T, Kelley L, Chayen NE, et al. Threedimensional structure of the human breast cancer resistance protein (BCRP/ABCG2) in an inward-facing conformation. Acta Crystallogr D Biol Crystallogr. 2015 Aug 1;71(8):1725–35.

