## Supplemental Information for "Acquired drug resistance enhances imidazoquinoline efflux by P-glycoprotein"

### Materials and Methods

#### Parent Cancer cell lines

B16: The B16-F10 melanoma cell line was purchased from the American Type Culture Collection (ATCC). As per manufacturer instructions, the B16-F10 cell line (ATCC CRL-6475, mouse melanoma) were grown in complete culture media composed of DMEM with 4.5 g $L^{-1}$ glucose, 2 mM L-glutamine, 100 U $mL^{-1}$ PenStrep, and supplemented with 10% HI-FBS. Media was changed every 3-4 days. Cells were passaged upon reaching 80% confluence. Passaging involved changing media, counting, and seeding 3 x $10^5$ cells in 35 mL of new complete media in a new T-175 culture flask.

TC2: The TRAMP-C2 prostate cell line was purchased from ATCC. As per manufacturer instructions, the TRAMP C2 cell line (ATCC CRL-2731, mouse transgenic adenocarcinoma) was grown in complete culture media composed of DMEM with 4.5 g $L^{-1}$ glucose, 2 mM L-glutamine, 100 U $mL^{-1}$ PenStrep, and supplemented with 1 µg $mL^{-1}$ insulin, 2 nM dehydroisoandrosterone, 5% HI-FBS, and 5% Nu-Serum IV. Media was changed every 3-4 days. Cells were passaged upon reaching 80% confluence. Passaging involved changing media, counting, and seeding 3 x $10^5$ cells in 35 mL of new complete media in a new T-175 culture flask.

4T1: 4T1-Luc2 breast cancer cell line was gifted from Darrell Irvine's lab (Massachusetts Institute of Technology, Cambridge, MA). The 4T1-Luc2 cell line (ATCC CRL-2539-LUC2, mouse mammary gland carcinoma) was grown in complete culture media composed of DMEM with 4.5 g $L^{-1}$ glucose, 2 mM L-glutamine, 100 U $mL^{-1}$ PenStrep, and supplemented with 10% HI-FBS. Media was changed every 3-4 days. Cells were passaged upon reaching 80% confluence. Passaging involved changing media, counting, and seeding 3 x $10^5$ cells in 35 mL of new complete media in a new T-175 culture flask.

AT3B-1 MDR: AT3B-1 MDR prostate cancer cells' were chosen for their known response to Doxorubicin, the upregulation of P-gp as a result of epigenetic pressure caused by exposure to the chemotherapeutic (1). The AT3B-1 cell line was purchased from ATCC. The AT3B-1 cell line (ATCC CRL-2375, mouse prostate) was grown in complete culture media composed DMEM with 4.5 g $L^{-1}$ glucose, 2 mM L-glutamine, 100 U $mL^{-1}$ PenStrep, supplemented with 10% HI-FBS and 1 µM Doxorubicin (TCI America - D4193100MG). Media was changed every 3-4 days. Cells were passaged upon reaching 80% confluence. Passaging involved changing media, counting, and seeding 3 x $10^5$ cells in 35 mL of new complete media in a new T-175 culture flask.

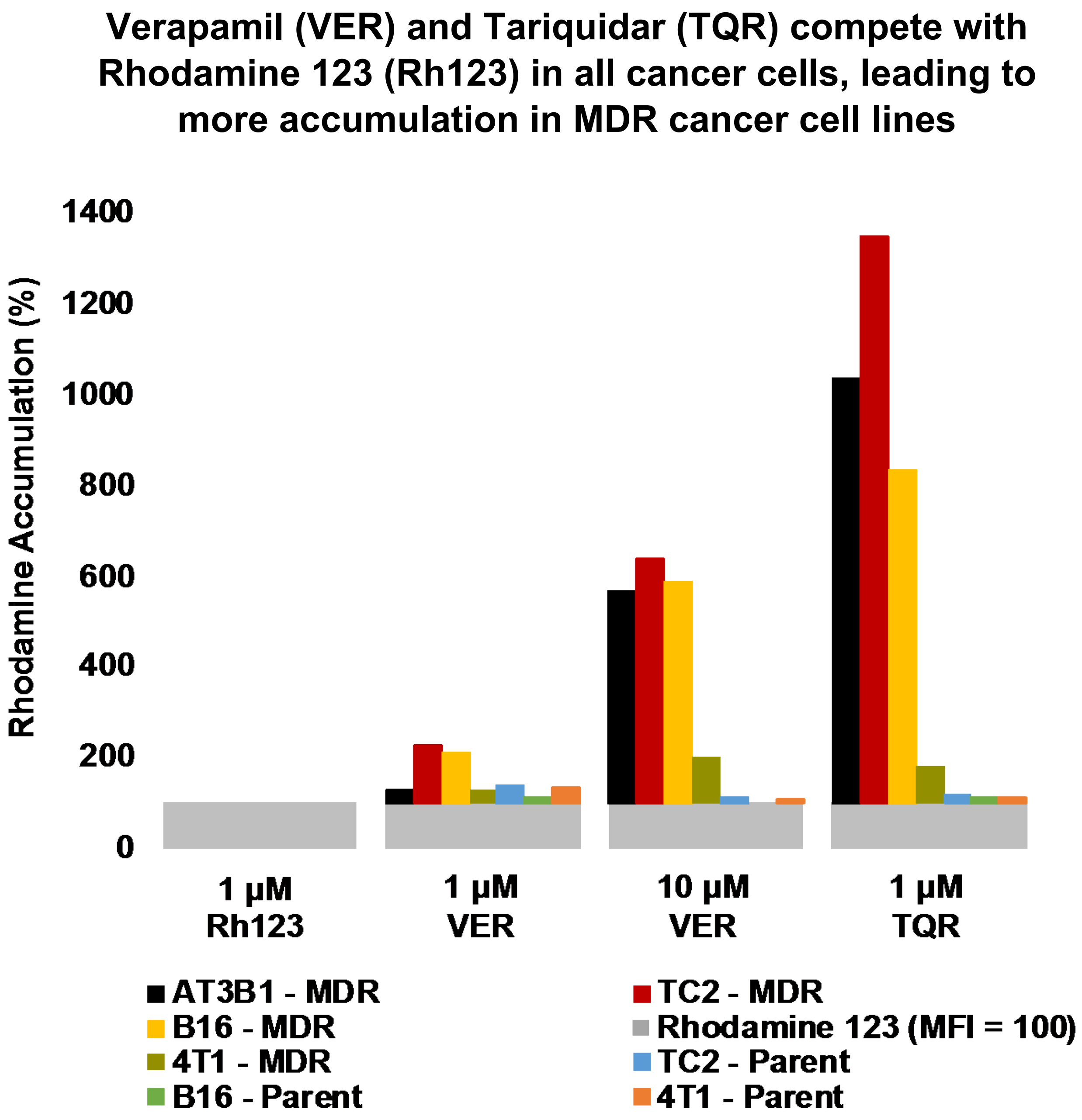

**Figure S1:** Competitive experiments with a known fluorescent substrate of P-gp, Rhodamine 123 (Rh123), in the presence of Verapamil (VER), a known competitive substrate and a first-generation inhibitor, and Tariquidar (TQR), a third-generation inhibitor. VER, and TQR compete with Rh123 in all of the cell lines tested. However, more Rh123 accumulation is observed in the MDR derived cancer cell lines with compared to the parent cancer cell lines. Data representative of experiments repeated in triplicate.

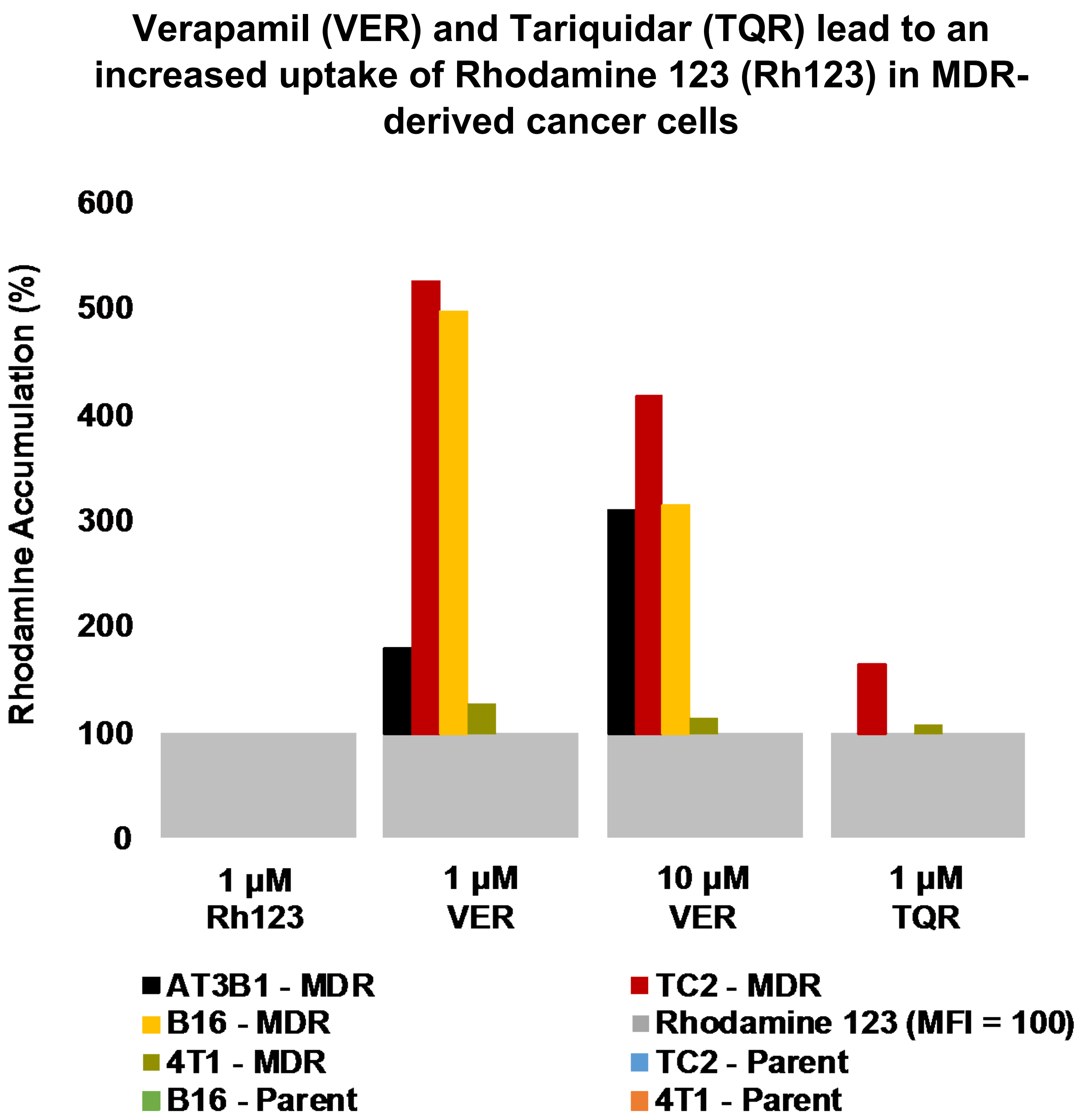

**Figure S2:** Uptake experiments with a known fluorescent substrate of P-gp, Rhodamine 123 (Rh123), in the presence of Verapamil (VER), a known competitive substrate and a first-generation inhibitor, and Tariquidar (TQR), a third-generation inhibitor. VER and TQR lead to an increased uptake of Rh123 under passive diffusion conditions in MDR-derived cancer cell lines. Data representative of experiments repeated in triplicate.

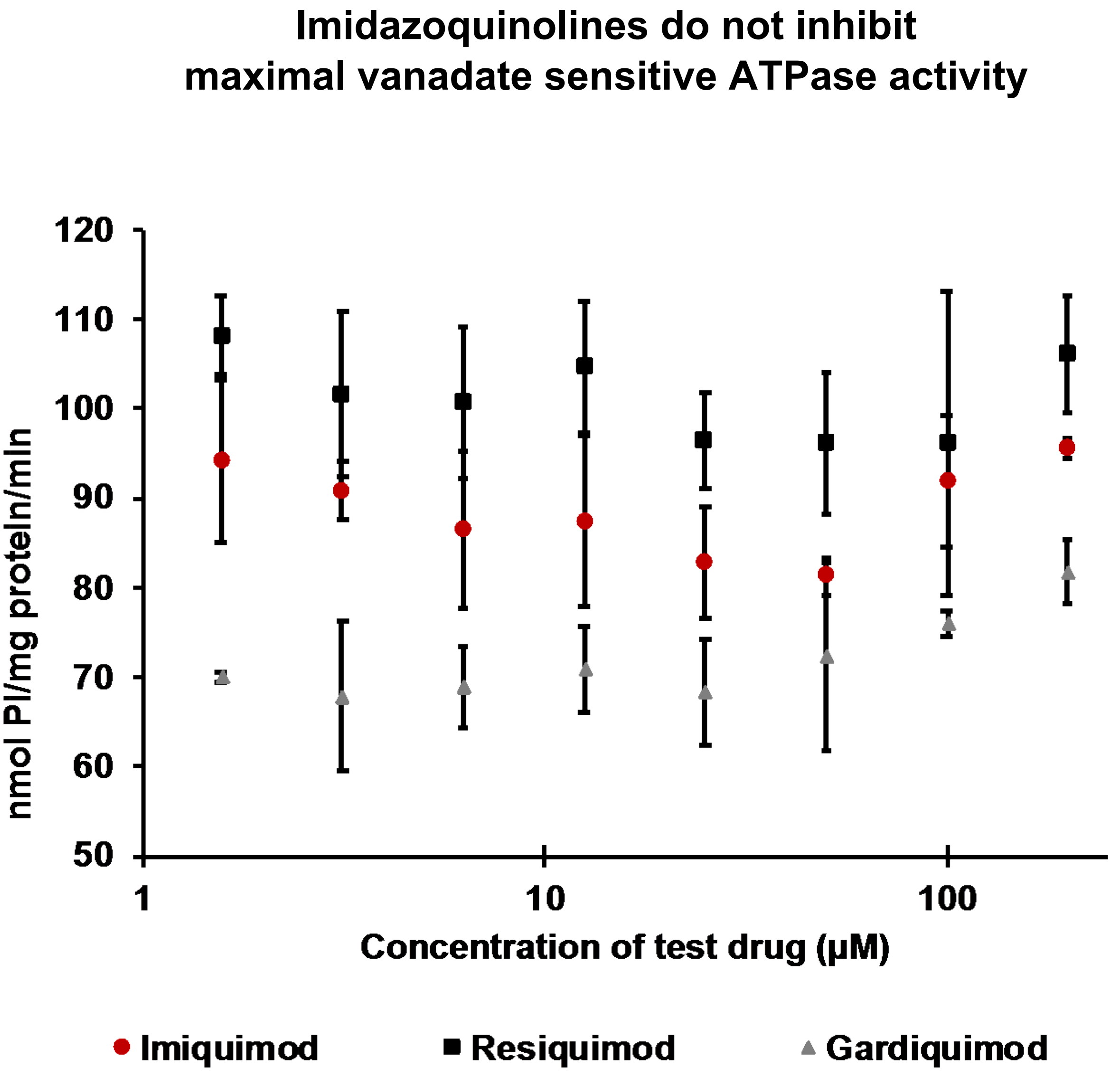

**Figure S3:** IMQ, RSQ, and GDQ do not inhibit maximal vanadate sensitive ATPase activity in the inhibition test, a complementary test from the ATPase assay. Because the inhibition test is carried out with a strong activator of the transporter, slowly transported compounds may inhibit maximal vanadate sensitive ATPase activity. Results indicate the interaction of the test compounds with the transporter, but do not give information on the nature (substrate or inhibitor) of the interaction in this inhibition assay.

#### Imidazoquinolines are not cytotoxic to cancer cells at loading concentrations

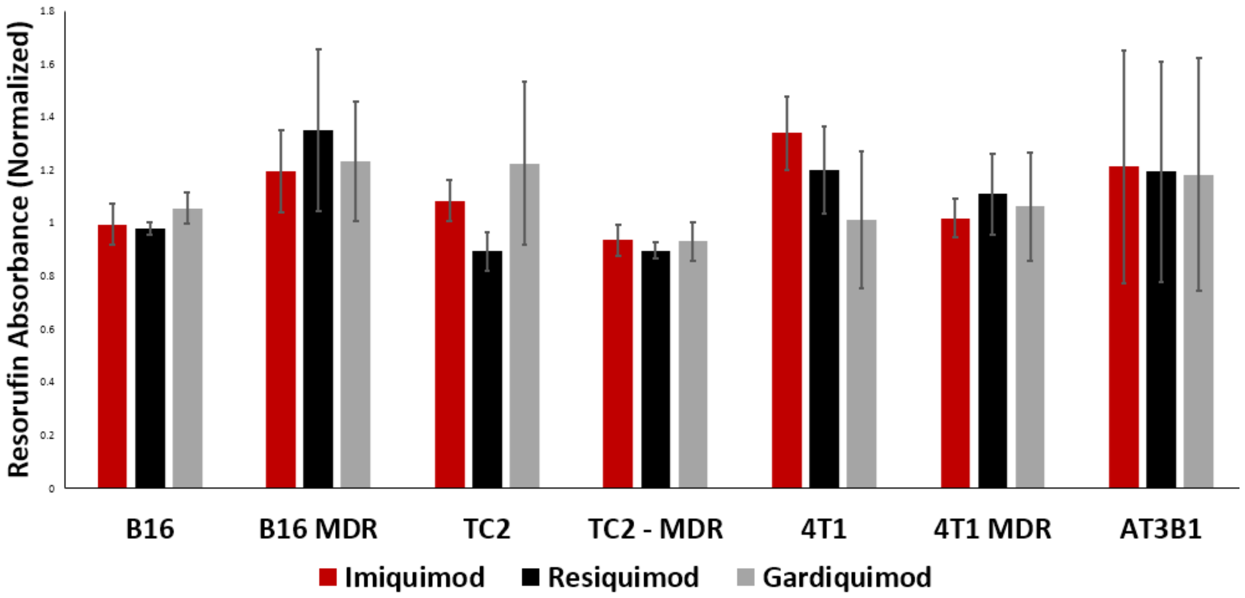

**Figure S4:** Resazurin Cell Viability Assay kit (Abcam - ab129732) was used to determine if the concentrations of imidazoquinolines used in experiments were cytotoxic to the cell lines tested: B16, B16 MDR, TC2, TC2 MDR, 4T1, 4T1 MDR, and AT3B1. Cells were treated with 20 μL of 1 mM solutions of imidazoquinoline in 5% DMSO in PBS to give a final concentration per well of 100 μM for each compound. Resazurin absorbance (600 nm) was subtracted from resorufin (570 nm). After this, the resulting values were normalized relative to blank (no cells) and cells incubated with DMSO / PBS alone. The data shown here represents a density of 50,000 cells/well with a 3h incubation time with stain.

#### Progression of TC2 MDR-derived cancer cell line from TC2 Parent cancer cell line

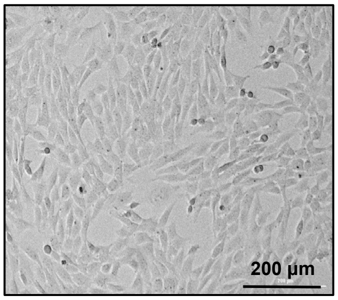

**Parent TRAMP C2**

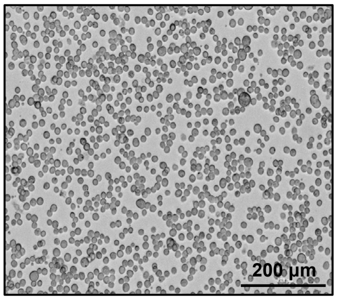

**4 nM Doxorubicin**

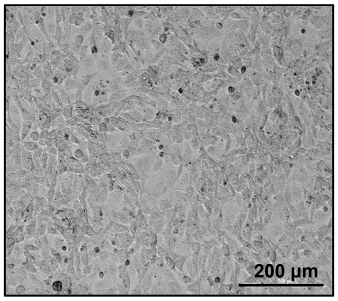

**8 nM Doxorubicin**

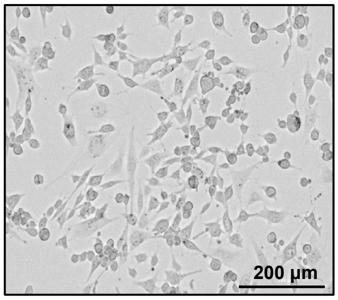

**16 nM Doxorubicin**

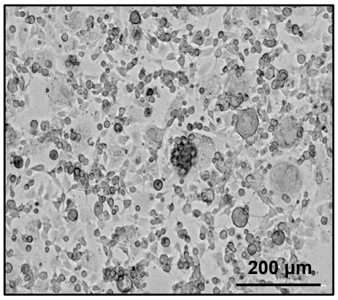

**32 nM Doxorubicin**

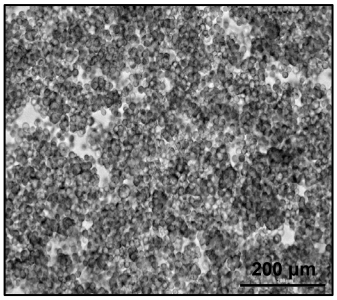

**64 nM Doxorubicin**

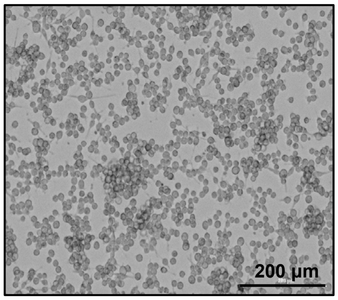

**128 nM Doxorubicin**

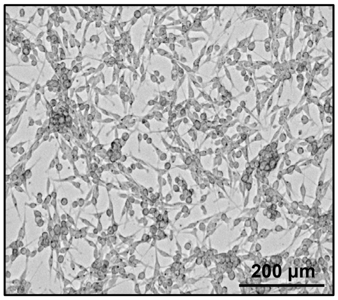

**256 nM Doxorubicin**

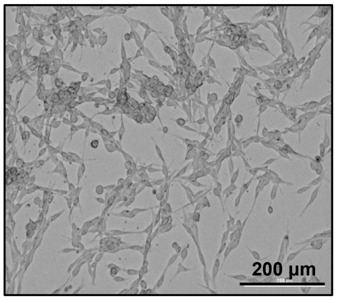

**500 nM Doxorubicin**

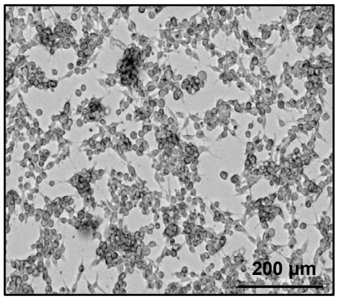

**1 µM Doxorubicin**

#### Progression of B16-MDR derived cancer cell line from B16 Parent cancer cell line

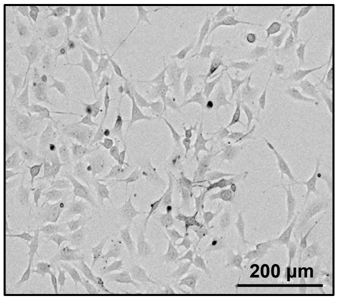

Parent B16

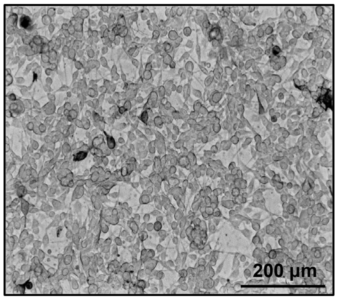

4 nM Doxorubicin

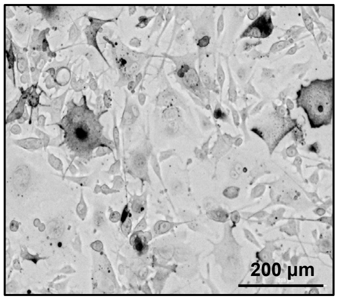

8 nM Doxorubicin

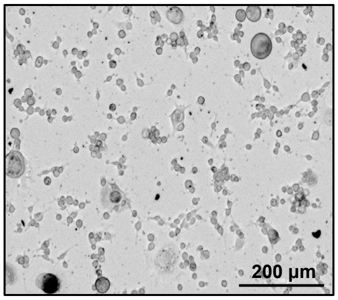

16 nM Doxorubicin

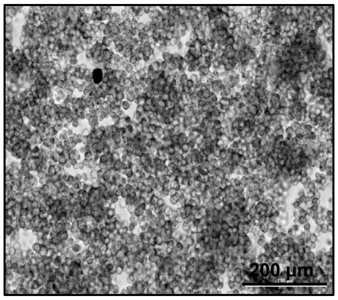

32 nM Doxorubicin

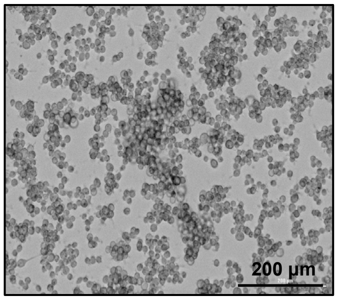

64 nM Doxorubicin

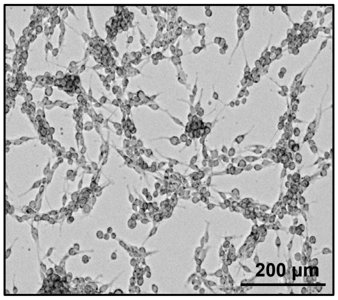

128 nM Doxorubicin

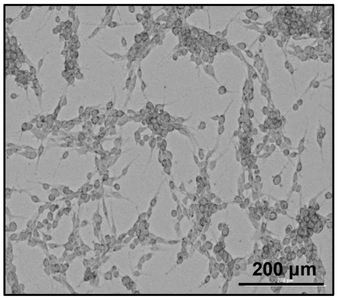

256 nM Doxorubicin

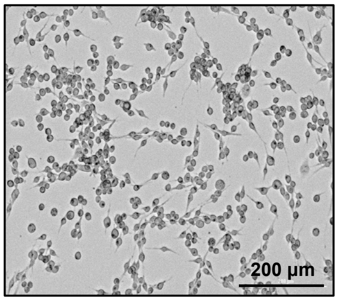

500 nM Doxorubicin

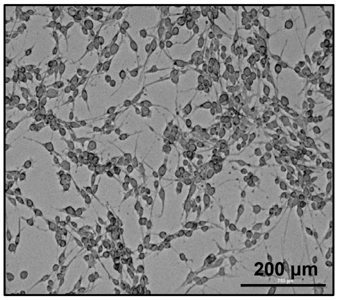

1 µM Doxorubicin

#### Progression of 4T1 MDR-derived cancer cell line from 4T1 Parent cancer cell line

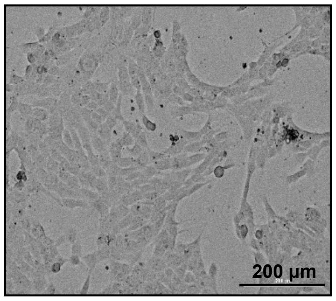

Parent 4T1

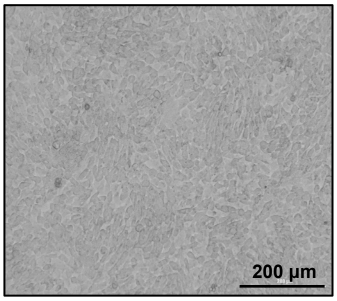

4 nM Doxorubicin

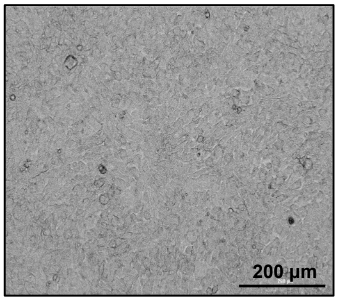

8 nM Doxorubicin

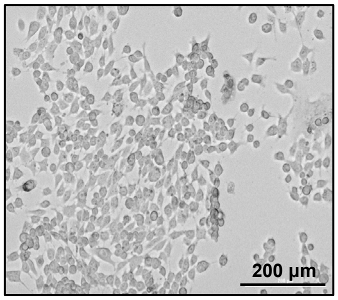

16 nM Doxorubicin

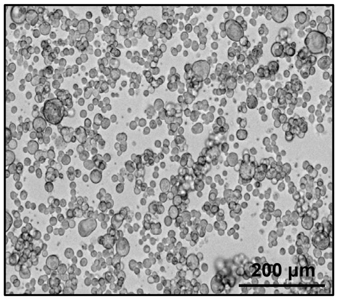

32 nM Doxorubicin

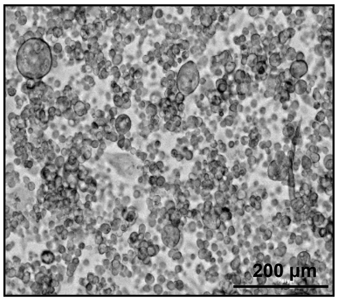

64 nM Doxorubicin

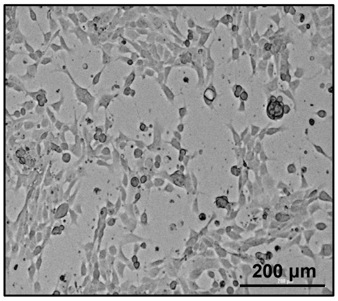

128 nM Doxorubicin

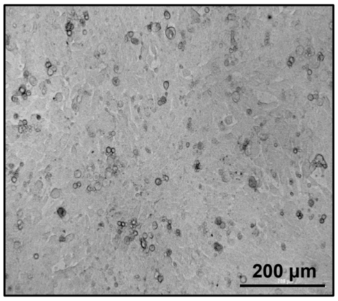

256 nM Doxorubicin

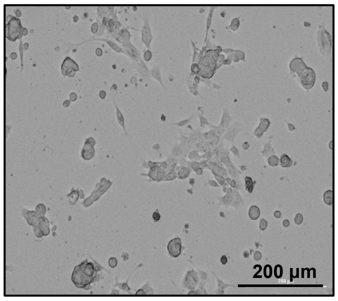

500 nM Doxorubicin

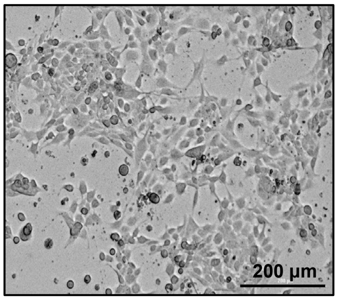

1 µM Doxorubicin

### Uptake Experiment with TC2 Parent Cancer Cell Line
### (Raw Flow Cytometry Plots)

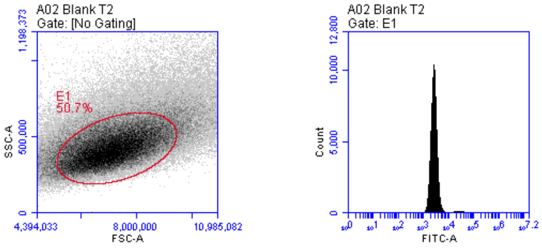

**Blank TRAMP C2 cells**

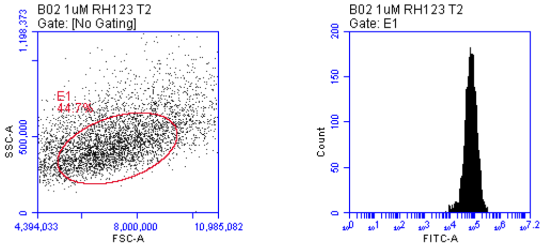

**1 μM Rhodamine 123**

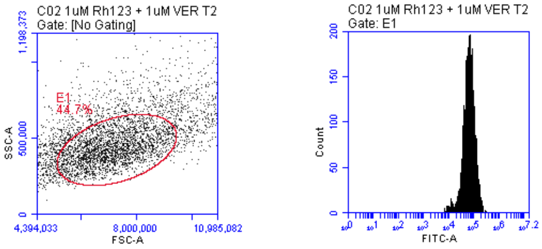

**1 μM Rhodamine 123 +
1 μM Verapamil**

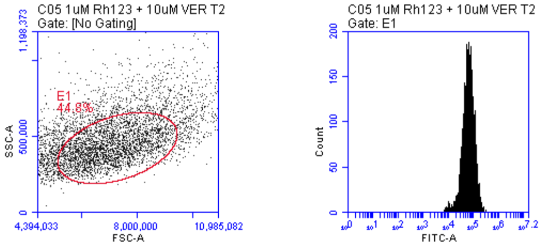

**1 μM Rhodamine 123 +
10 μM Verapamil**

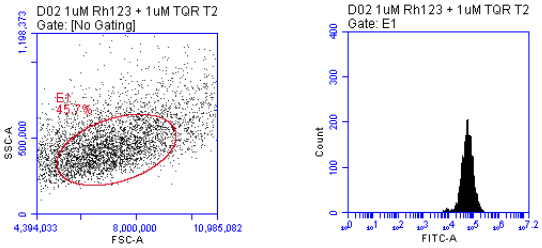

**1 μM Rhodamine 123 +
1 μM Tariquidar**

### Uptake Experiment with TC2 Parent Cancer Cell Line
### (Raw Flow Cytometry Plots)

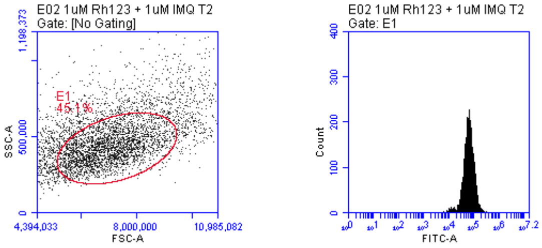

**1 µM Rhodamine 123 +**
**1 µM Imiquimod**

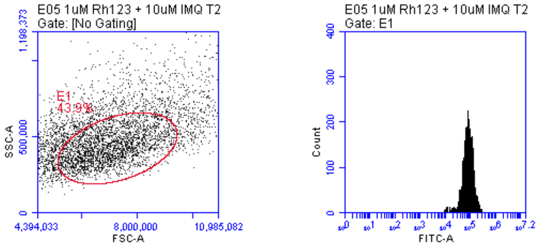

**1 µM Rhodamine 123 +**
**10 µM Imiquimod**

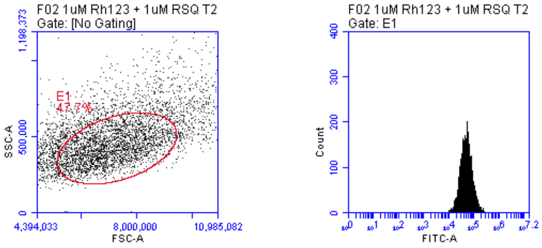

**1 µM Rhodamine 123 +**
**1 µM Resiquimod**

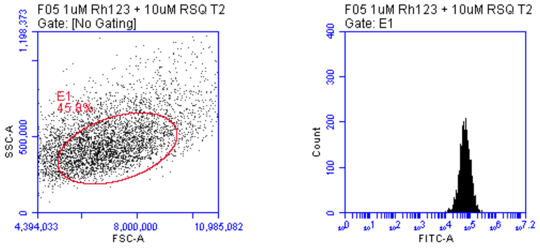

**1 µM Rhodamine 123 +**
**10 µM Resiquimod**

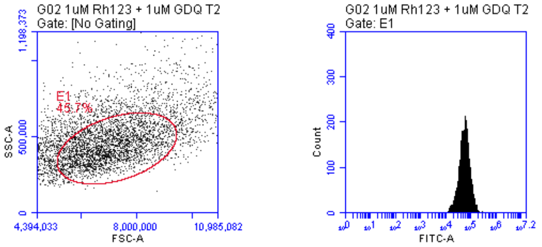

**1 µM Rhodamine 123 +**
**1 µM Gardiquimod**

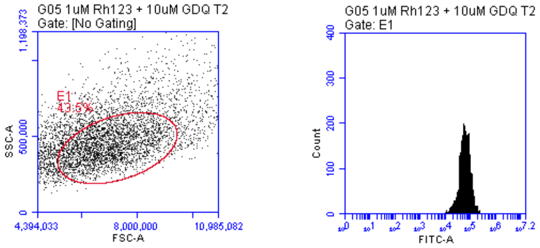

**1 µM Rhodamine 123 +**
**10 µM Gardiquimod**

### Uptake Experiment with B16 Parent Cancer Cell Line (Raw Flow Cytometry Plots)

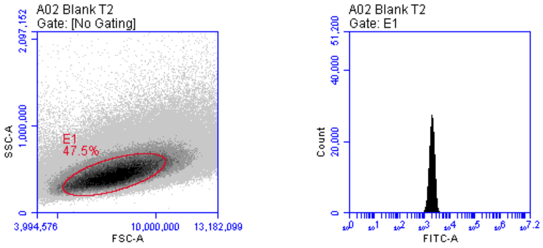

**Blank B16 cells**

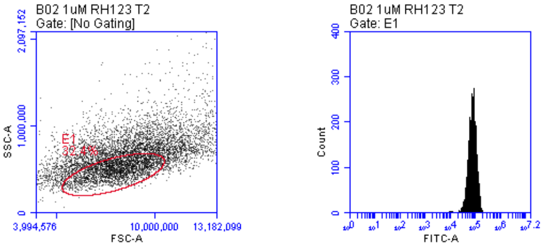

**1 μM Rhodamine**

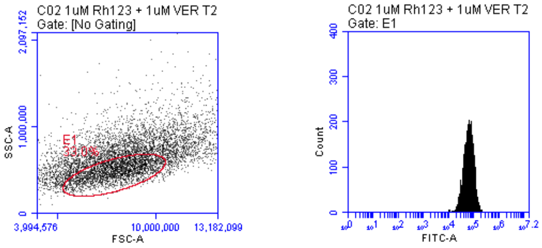

**1 μM Rhodamine 123 + 1 μM Verapamil**

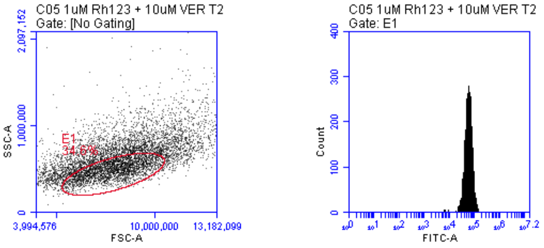

**1 μM Rhodamine 123 + 10 μM Verapamil**

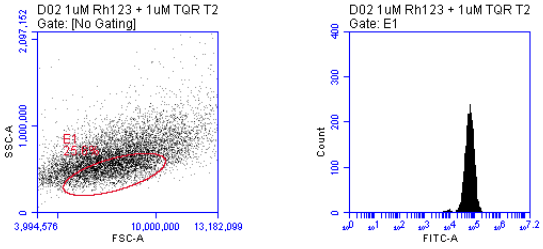

**1 μM Rhodamine 123 + 1 μM Tariquidar**

### Uptake Experiment with B16 Parent Cancer Cell Line (Raw Flow Cytometry Plots)

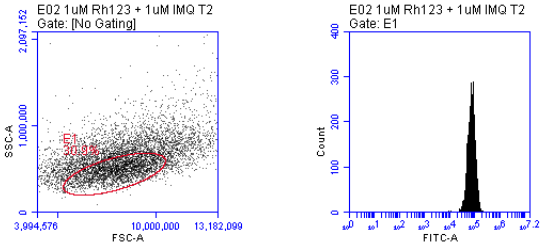

**1 µM Rhodamine 123 +
1µM Imiquimod**

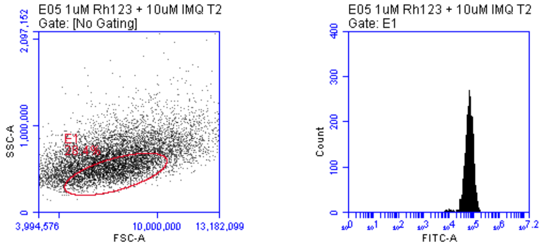

**1 µM Rhodamine 123 +
10 µM Imiquimod**

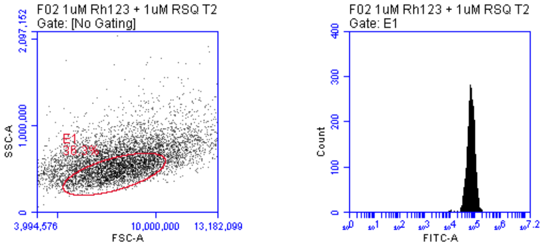

**1 µM Rhodamine 123 +
1 µM Resiquimod**

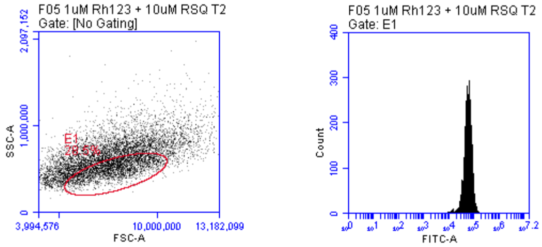

**1 µM Rhodamine 123 +
10 µM Resiquimod**

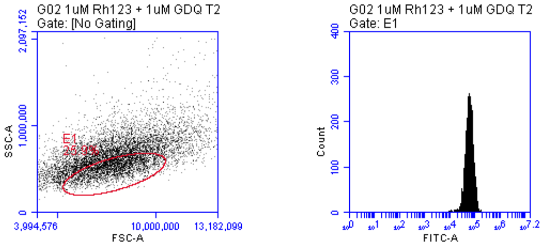

**1 µM Rhodamine 123 +
1 µM Gardiquimod**

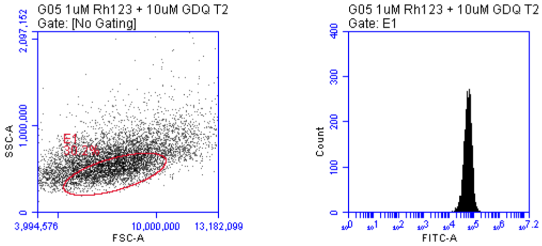

**1 µM Rhodamine 123 +
10 µM Gardiquimod**

### Uptake Experiment with 4T1 Parent Cancer Cell Line
### (Raw Flow Cytometry Plots)

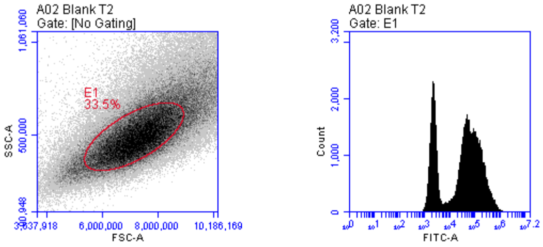

**Blank 4T1 cells**

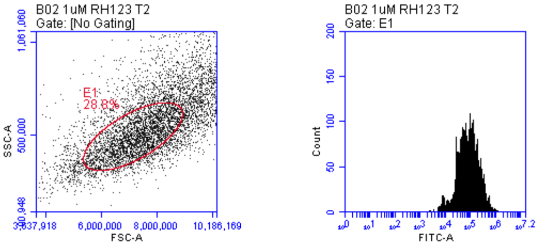

**1 μM Rhodamine 123**

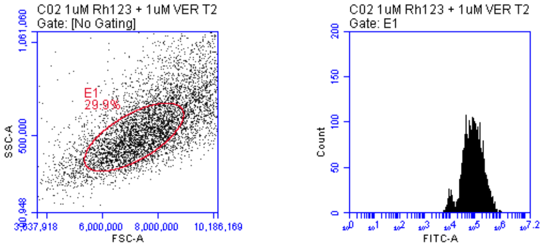

**1 μM Rhodamine 123 +**
**1 μM Verapamil**

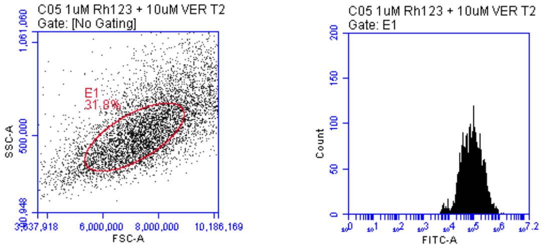

**1 μM Rhodamine 123 +**
**10 μM Verapamil**

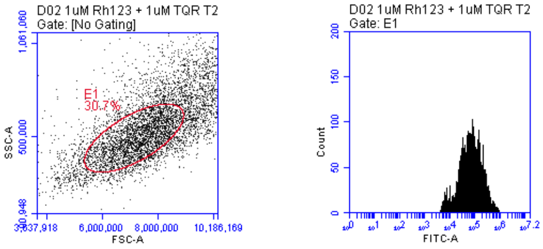

**1 μM Rhodamine 123 +**
**1 μM Tariquidar**

### Uptake Experiment with 4T1 Parent Cancer Cell Line
### (Raw Flow Cytometry Plots)

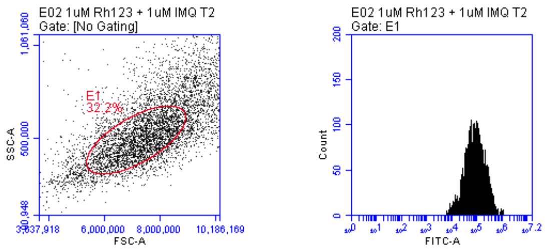

**1 µM Rhodamine 123 +
1 µM Imiquimod**

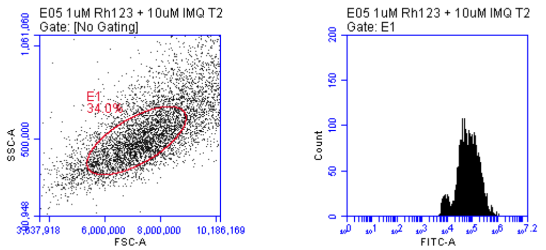

**1 µM Rhodamine 123 +
10 µM Imiquimod**

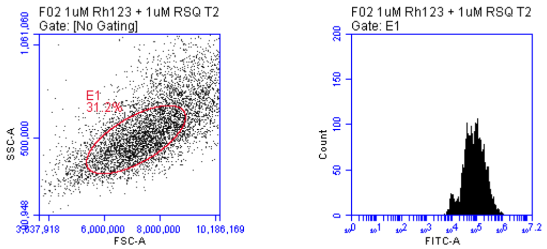

**1 µM Rhodamine 123 +
1 µM Resiquimod**

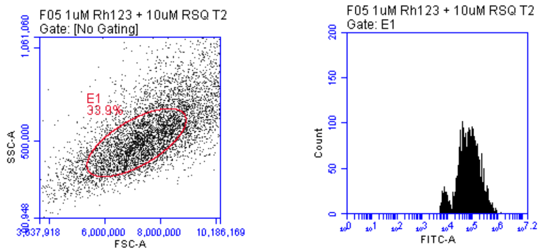

**1 µM Rhodamine 123 +
10 µM Resiquimod**

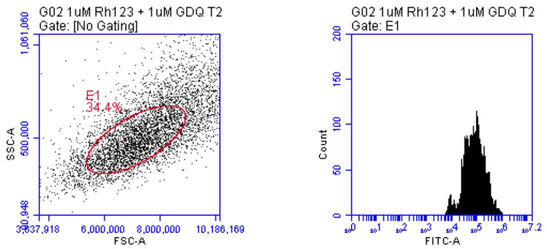

**1 µM Rhodamine 123 +
1 µM Gardiquimod**

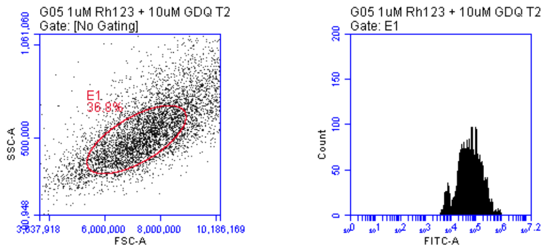

**1 µM Rhodamine 123 +
10 µM Gardiquimod**

### Uptake Experiment with TC2-MDR Cancer Cell Line
### (Raw Flow Cytometry Plots)

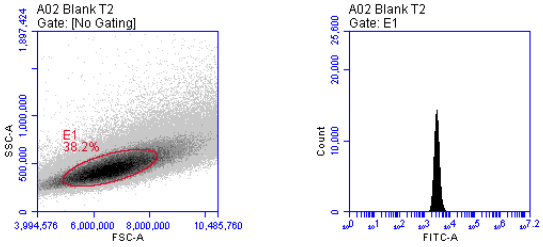

**Blank TC2 - MDR cells**

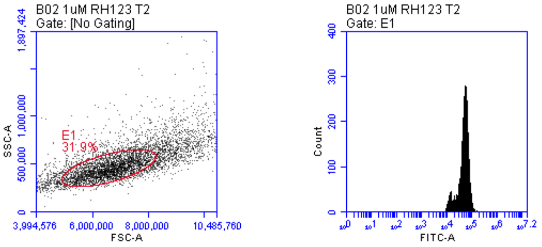

**1 μM Rhodamine 123**

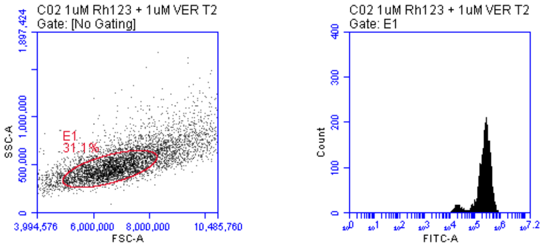

**1 μM Rhodamine 123 +**
**1 μM Verapamil**

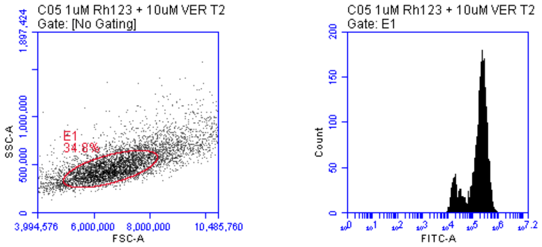

**1 μM Rhodamine 123 +**
**10 μM Verapamil**

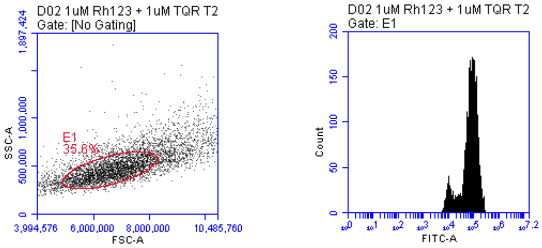

**1 μM Rhodamine 123 +**
**1 μM Tariquidar**

### Uptake Experiment with TC2-MDR Cancer Cell Line
### (Raw Flow Cytometry Plots)

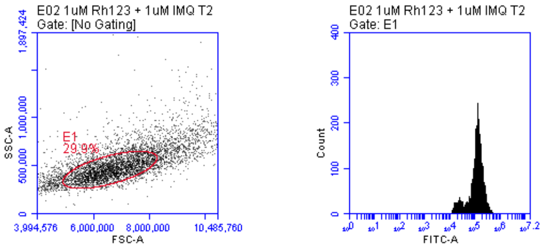

**1 µM Rhodamine 123 + 1µM Imiquimod**

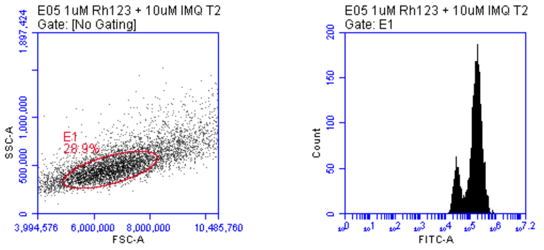

**1 µM Rhodamine 123 + 10µM Imiquimod**

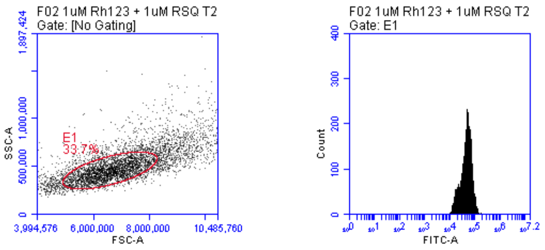

**1 µM Rhodamine 123 + 1 µM Resiquimod**

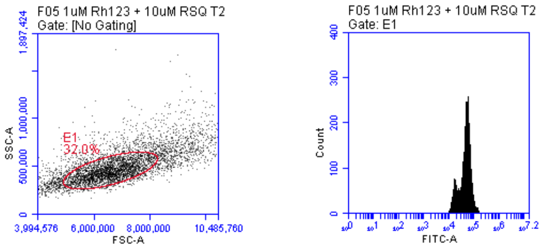

**1 µM Rhodamine 123 + 10 µM Resiquimod**

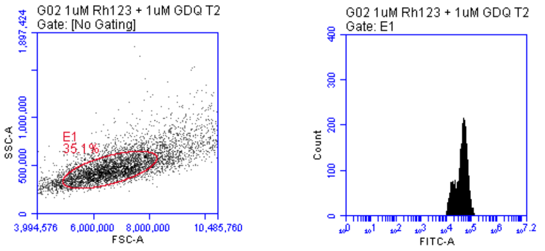

**1 µM Rhodamine 123 + 1 µM Gardiquimod**

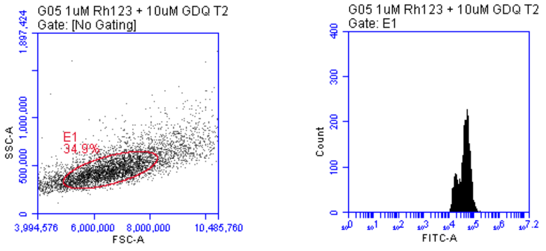

**1 µM Rhodamine 123 + 10 µM Gardiquimod**

### Uptake Experiment with B16-MDR Cancer Cell Line
### (Raw Flow Cytometry Plots)

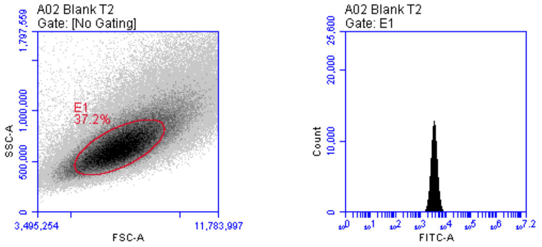

**Blank B16 - MDR cells**

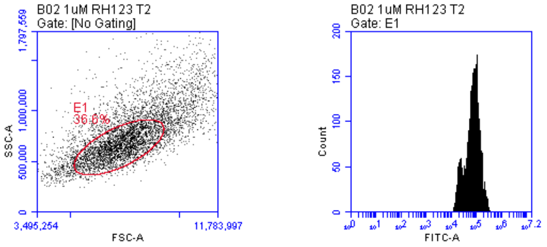

**1 µM Rhodamine 123**

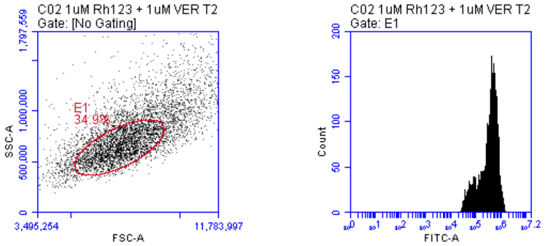

**1 µM Rhodamine 123 +**
**1 µM Verapamil**

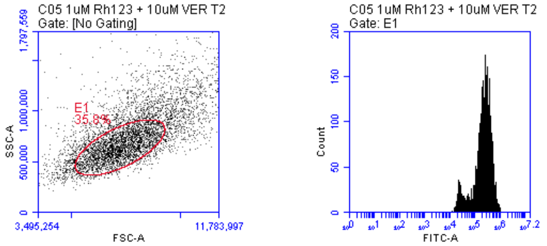

**1 µM Rhodamine 123 +**
**10 µM Verapamil**

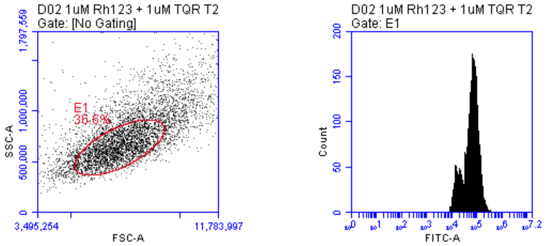

**1 µM Rhodamine 123 +**
**1 µM Tariquidar**

### Uptake Experiment with B16-MDR Cancer Cell Line
### (Raw Flow Cytometry Plots)

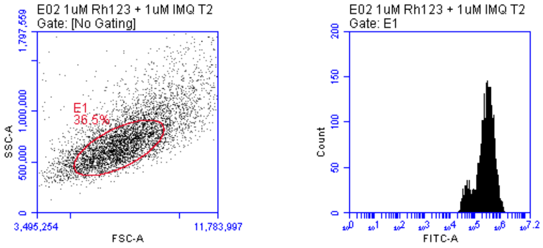

**1 μM Rhodamine 123 +**
**1 μM Imiquimod**

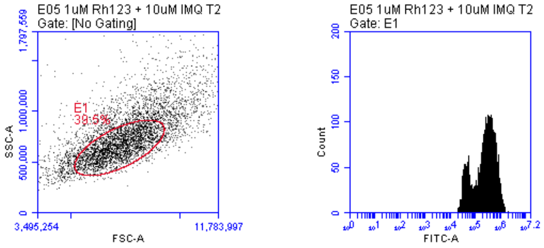

**1 μM Rhodamine 123 +**
**10 μM Imiquimod**

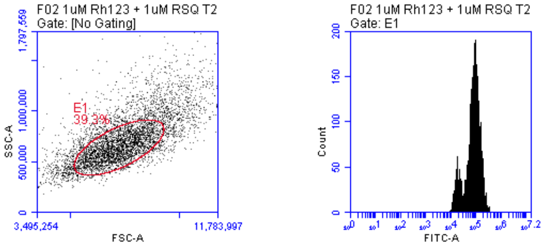

**1 μM Rhodamine 123 +**
**1 μM Resiquimod**

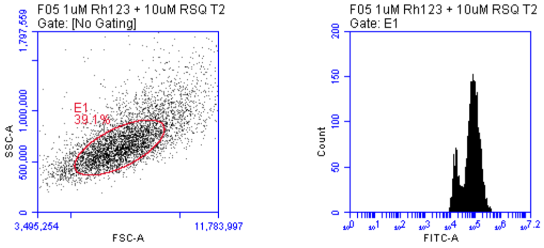

**1 μM Rhodamine 123 +**
**10 μM Resiquimod**

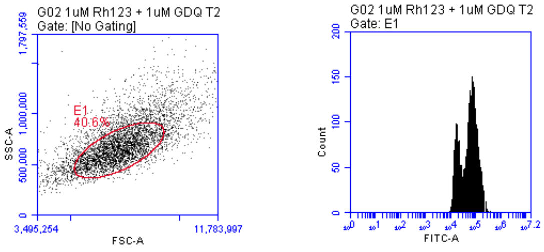

**1 μM Rhodamine 123 +**
**1 μM Gardiquimod**

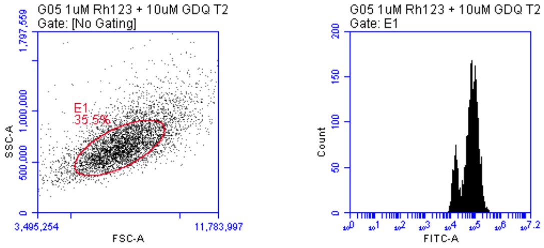

**1 μM Rhodamine 123 +**
**10 μM Gardiquimod**

### Uptake Experiment with 4T1-MDR Cancer Cell Line
### (Raw Flow Cytometry Plots)

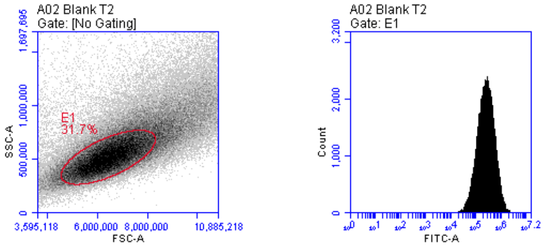

**Blank 4T1 - MDR cells**

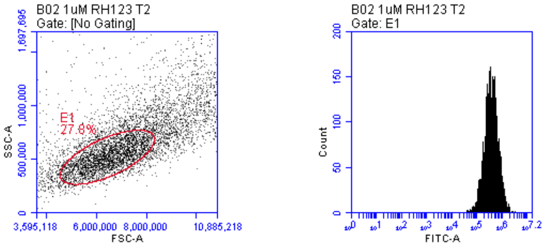

**1 µM Rhodamine 123**

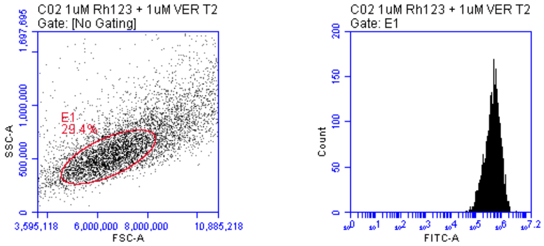

**1 µM Rhodamine 123 +**
**1 µM Verapamil**

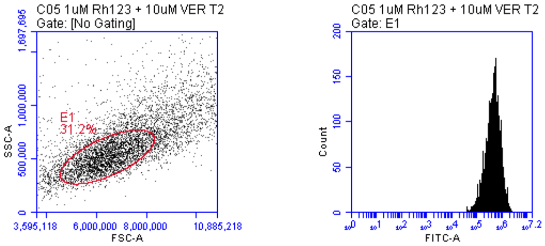

**1 µM Rhodamine 123 +**
**10 µM Verapamil**

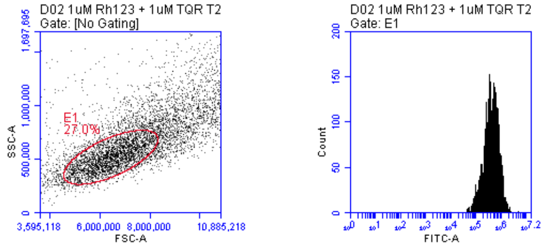

**1 µM Rhodamine 123 +**
**1 µM Tariquidar**

### Uptake Experiment with 4T1 MDR Cancer Cell Line
### (Raw Flow Cytometry Plots)

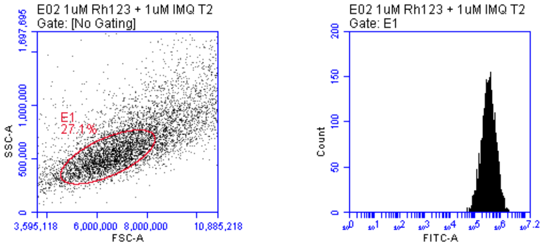

**1 µM Rhodamine 123 +**
**1 µM Imiquimod**

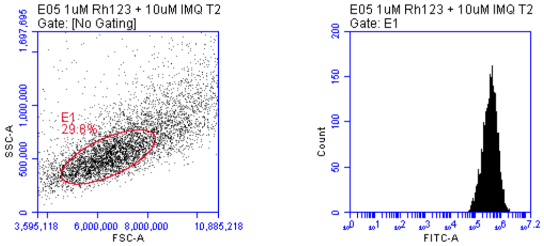

**1 µM Rhodamine 123 +**
**10 µM Imiquimod**

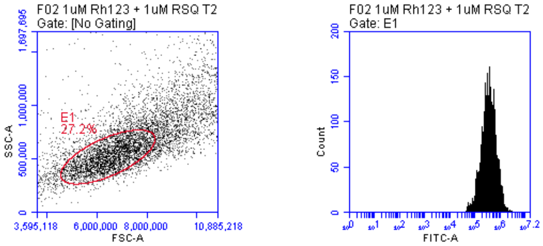

**1 µM Rhodamine 123 +**
**1 µM Resiquimod**

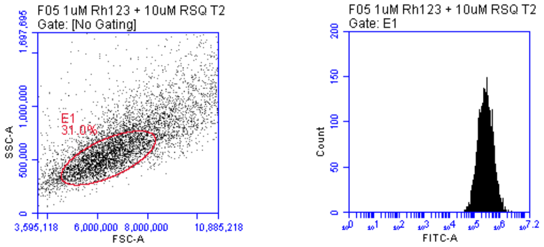

**1 µM Rhodamine 123 +**
**10 µM Resiquimod**

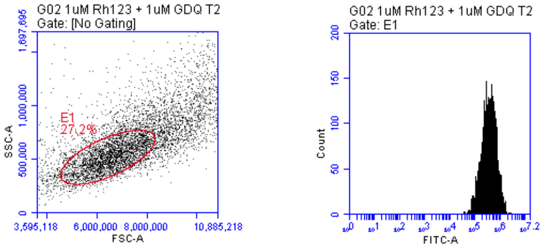

**1 µM Rhodamine 123 +**
**1 µM Gardiquimod**

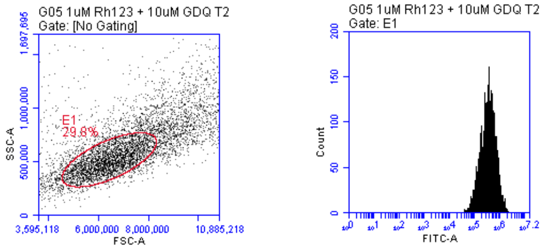

**1 µM Rhodamine 123 +**
**10 µM Gardiquimod**

### Uptake Experiment with AT3B-1 MDR Cancer Cell Line
### (Raw Flow Cytometry Plots)

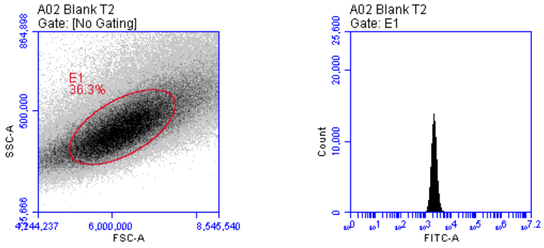

**Blank AT3B-1 MDR cells**

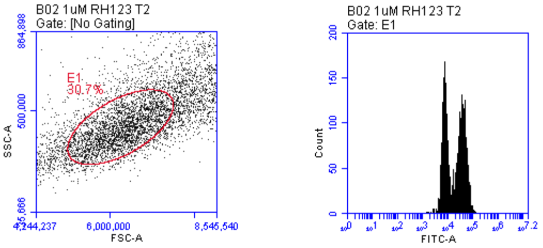

**1 µM Rhodamine 123**

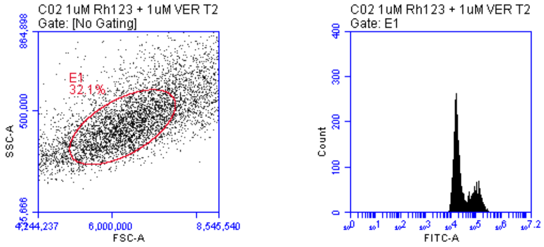

**1 µM Rhodamine 123 +**
**1 µM Verapamil**

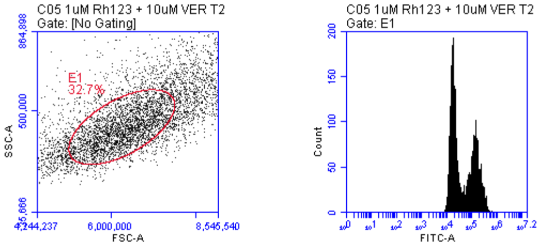

**1 µM Rhodamine 123 +**
**10 µM Verapamil**

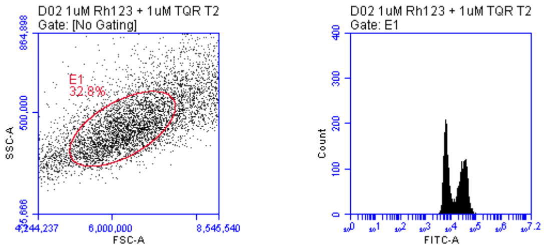

**1 µM Rhodamine 123 +**
**1 µM Tariquidar**

### Uptake Experiment with AT3B-1 MDR Cancer Cell Line (Raw Flow Cytometry Plots)

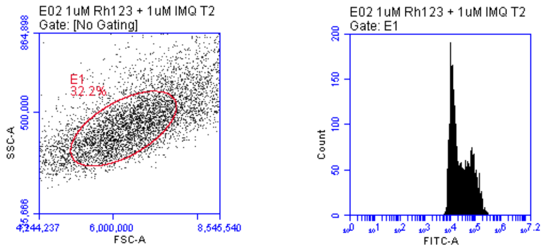

**1 µM Rhodamine 123 +
1 µM Imiquimod**

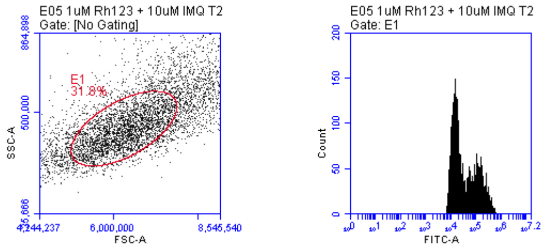

**1 µM Rhodamine 123 +
10 µM Imiquimod**

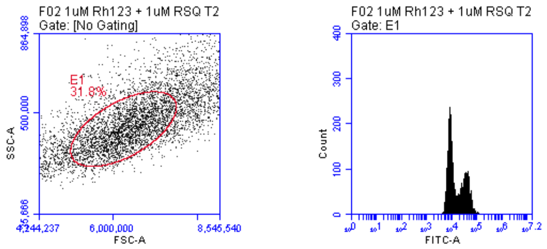

**1 µM Rhodamine 123 +
1 µM Resiquimod**

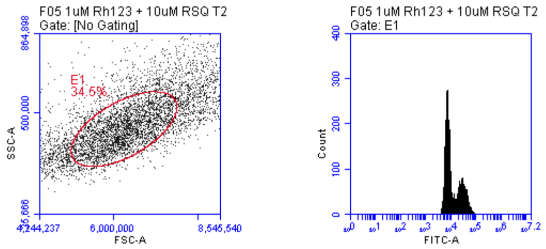

**1 µM Rhodamine 123 +
10 µM Resiquimod**

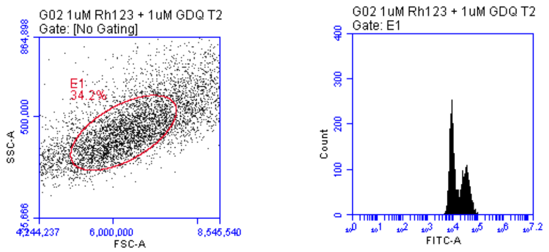

**1 µM Rhodamine 123 +
1 µM Gardiquimod**

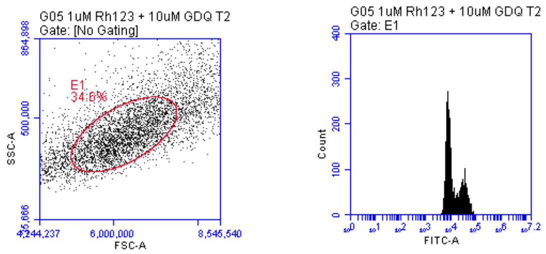

**1 µM Rhodamine 123 +
10 µM Gardiquimod**

### Competitive Experiment with TC2 Parent Cancer Cell Line (Raw Flow Cytometry Plots)

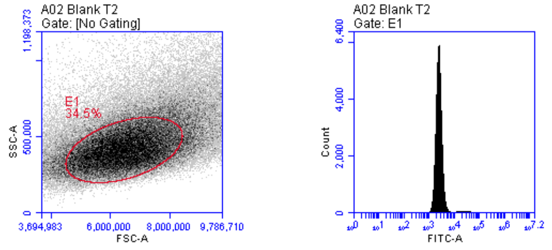

**Blank TRAMP C2 cells**

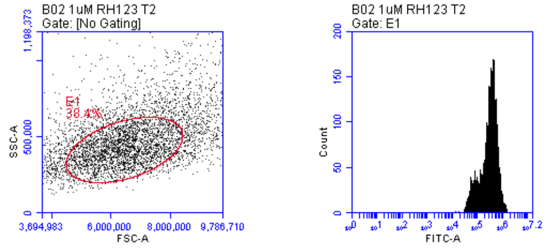

**1 µM Rhodamine 123**

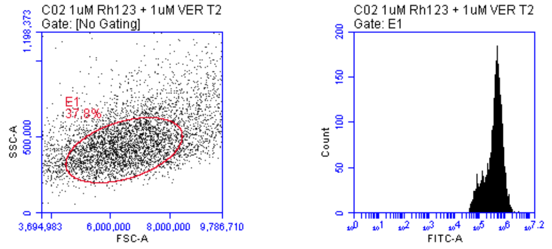

**1 µM Rhodamine 123 + 1 µM Verapamil**

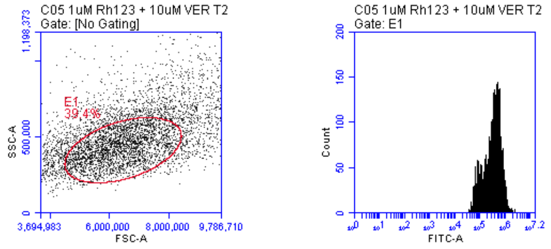

**1 µM Rhodamine 123 + 10 µM Verapamil**

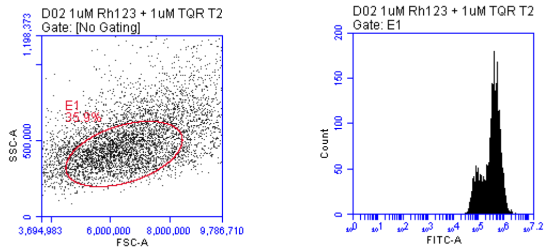

**1 µM Rhodamine 123 + 1 µM Tariquidar**

### Competitive Experiment with TC2 Parent Cancer Cell Line (Raw Flow Cytometry Plots)

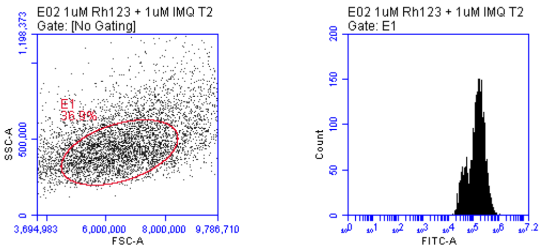

**1 µM Rhodamine 123 + 1 µM Imiquimod**

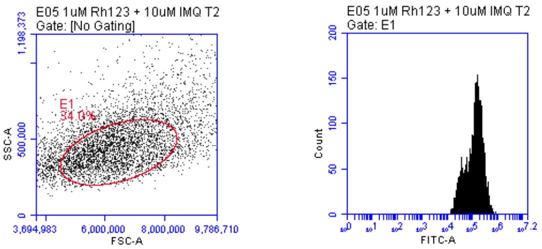

**1 µM Rhodamine 123 + 10 µM Imiquimod**

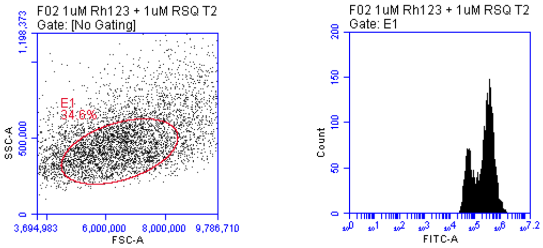

**1 µM Rhodamine 123 + 1 µM Resiquimod**

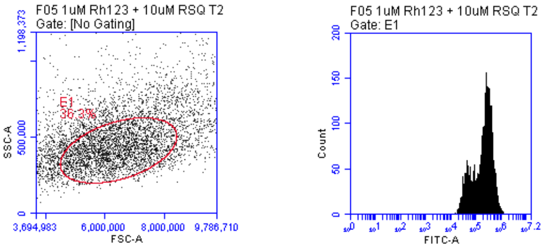

**1 µM Rhodamine 123 + 10 µM Resiquimod**

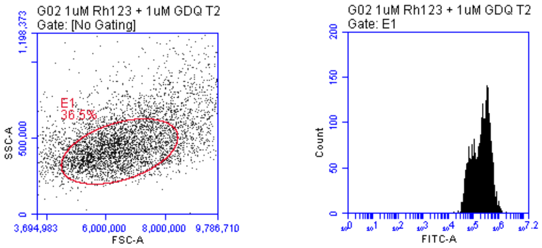

**1 µM Rhodamine 123 + 1 µM Gardiquimod**

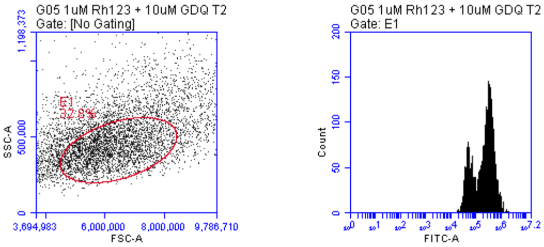

**1 µM Rhodamine 123 + 10 µM Gardiquimod**

### Competitive Experiment with B16 Parent Cancer Cell Line
### (Raw Flow Cytometry Plots)

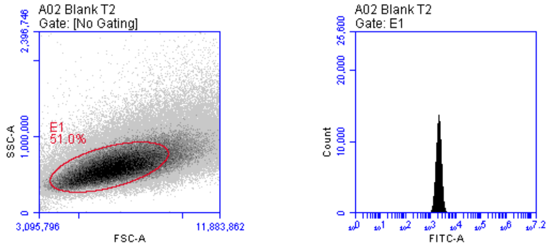

**Blank B16 cells**

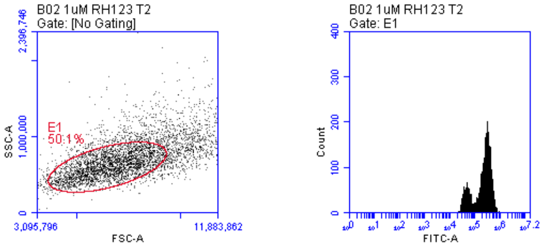

**1 μM Rhodamine 123**

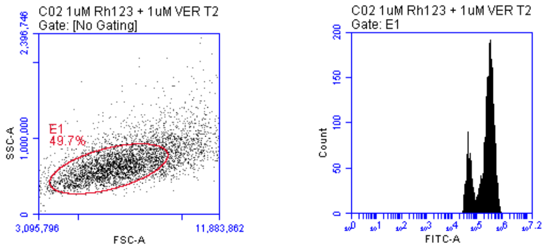

**1 μM Rhodamine 123 +
1 μM Verapamil**

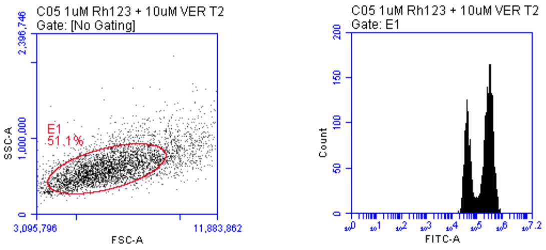

**1 μM Rhodamine 123 +
10 μM Verapamil**

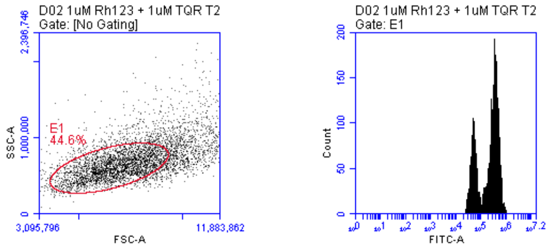

**1 μM Rhodamine 123 +
1 μM Tariquidar**

### Competitive Experiment with B16 Parent Cancer Cell Line
### (Raw Flow Cytometry Plots)

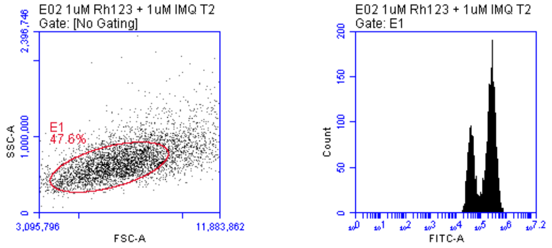

**1 μM Rhodamine 123 +**
**1 μM Imiquimod**

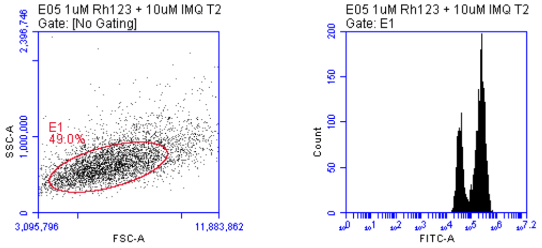

**1 μM Rhodamine 123 +**
**10 μM Imiquimod**

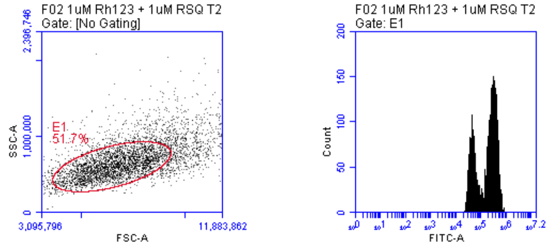

**1 μM Rhodamine 123 +**
**1 μM Resiquimod**

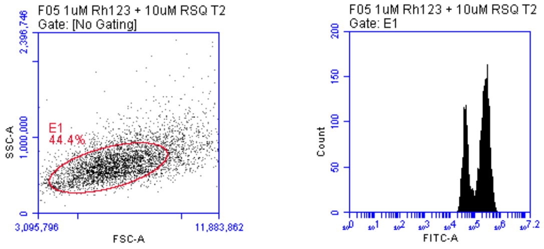

**1 μM Rhodamine 123 +**
**10 μM Resiquimod**

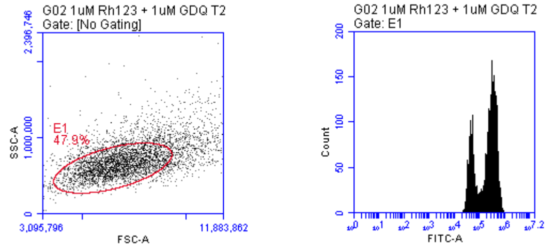

**1 μM Rhodamine 123 +**
**1 μM Gardiquimod**

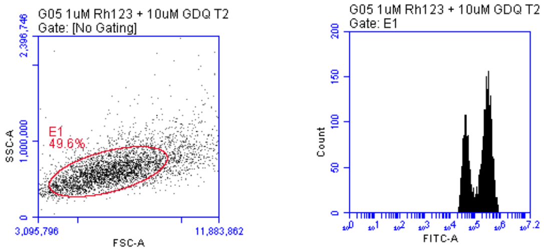

**1 μM Rhodamine 123 +**
**10 μM Gardiquimod**

### Competitive Experiment with 4T1 Parent Cancer Cell Line (Raw Flow Cytometry Plots)

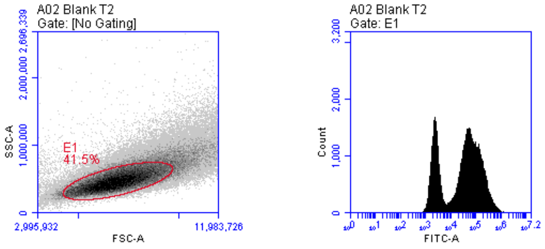

**Blank 4T1 cells**

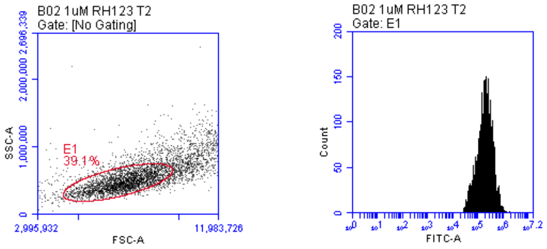

**1 μM Rhodamine 123**

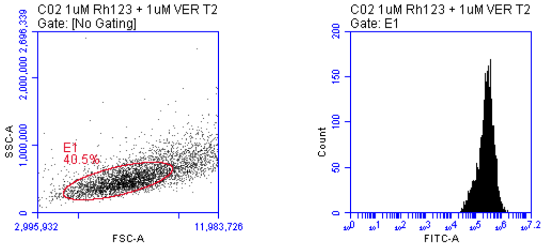

**1 μM Rhodamine 123 + 1 μM Verapamil**

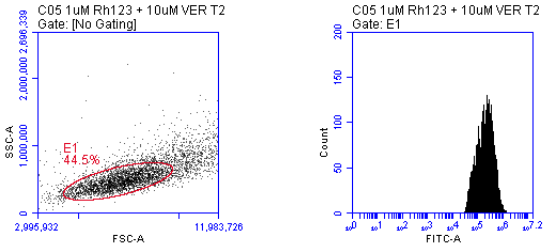

**1 μM Rhodamine 123 + 10 μM Verapamil**

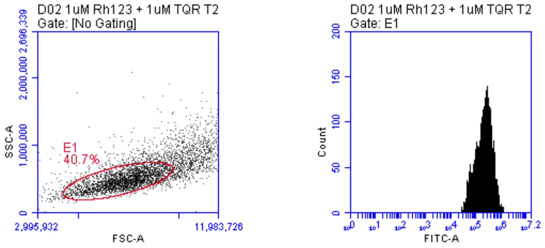

**1 μM Rhodamine 123 + 1 μM Tariquidar**

### Competitive Experiment with 4T1 Parent Cancer Cell Line (Raw Flow Cytometry Plots)

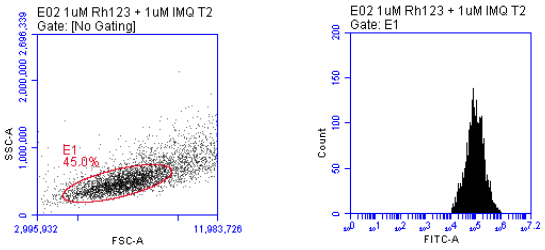

**1 µM Rhodamine 123 + 1 µM Imiquimod**

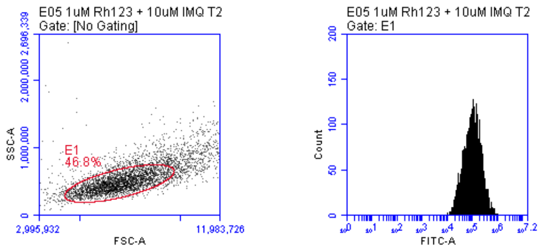

**1 µM Rhodamine 123 + 10 µM Imiquimod**

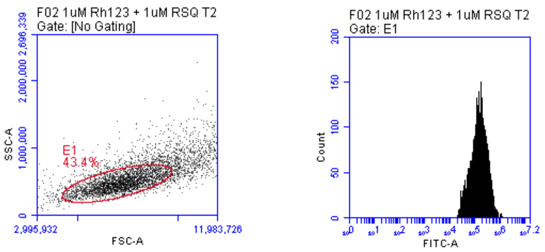

**1 µM Rhodamine 123 + 1 µM Resiquimod**

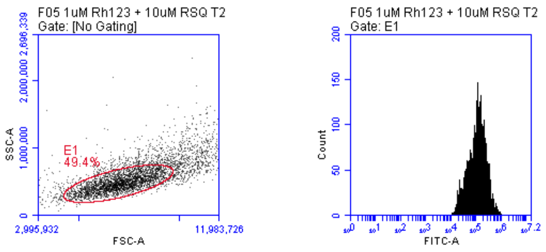

**1 µM Rhodamine 123 + 10 µM Resiquimod**

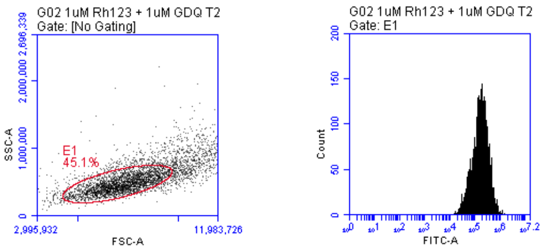

**1 µM Rhodamine 123 + 1 µM Gardiquimod**

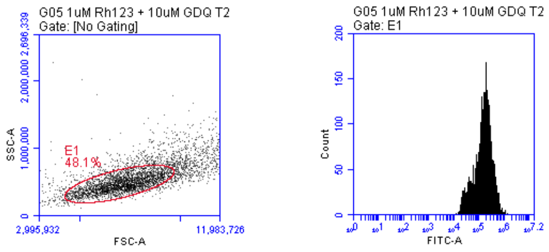

**1 µM Rhodamine 123 + 10 µM Gardiquimod**

### Competitive Experiment with TC2-MDR Cancer Cell Line (Raw Flow Cytometry Plots)

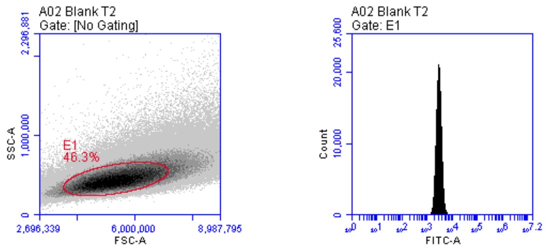

**Blank TRAMP C2 - MDR cells**

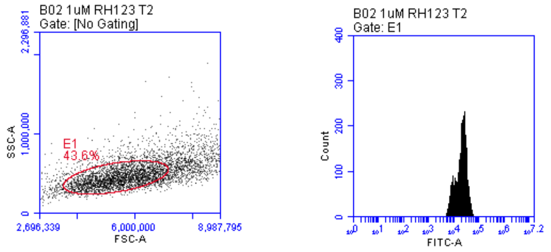

**1 µM Rhodamine 123**

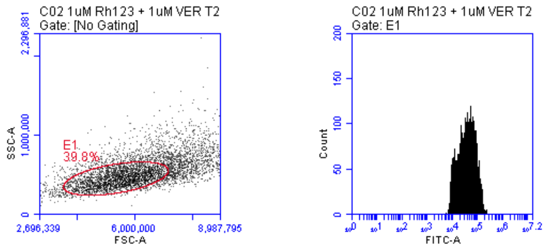

**1 µM Rhodamine 123 + 1 µM Verapamil**

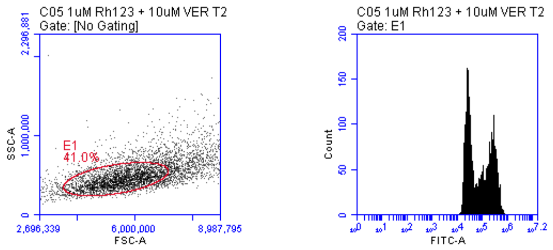

**1 µM Rhodamine 123 + 10 µM Verapamil**

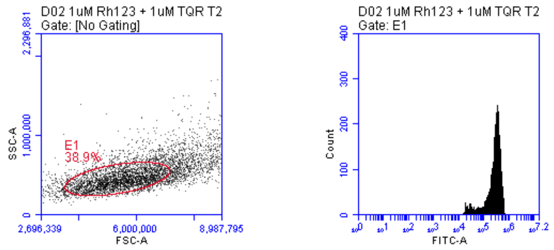

**1 µM Rhodamine 123 + 1 µM Tariquidar**

### Competitive Experiment with TC2-MDR Cancer Cell Line (Raw Flow Cytometry Plots)

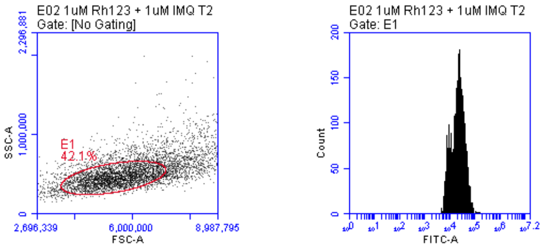

**1 µM Rhodamine 123 + 1 µM Imiquimod**

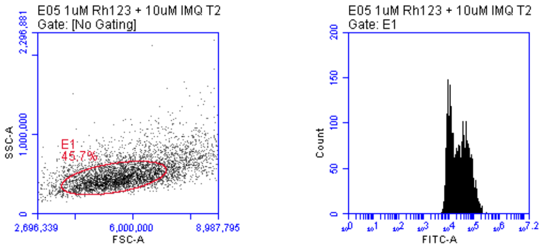

**1 µM Rhodamine 123 + 10 µM Imiquimod**

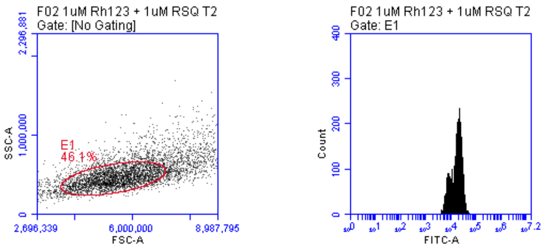

**1 µM Rhodamine 123 + 1 µM Resiquimod**

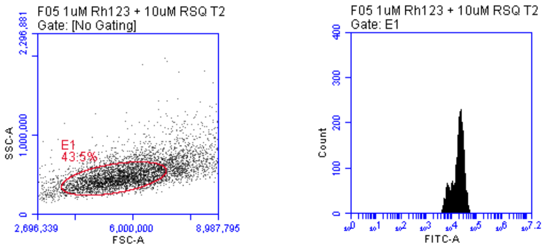

**1 µM Rhodamine 123 + 10 µM Resiquimod**

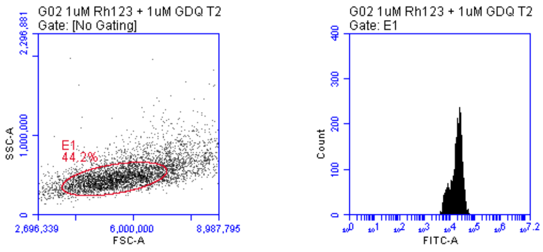

**1 µM Rhodamine 123 + 1 µM Gardiquimod**

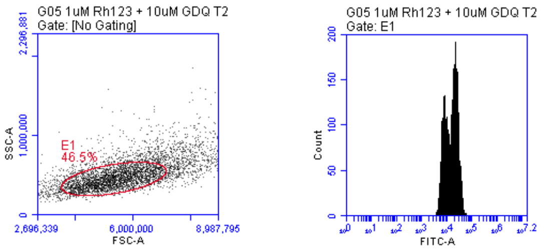

**1 µM Rhodamine 123 + 10 µM Gardiquimod**

### Competitive Experiment with B16-MDR Cancer Cell Line
### (Raw Flow Cytometry Plots)

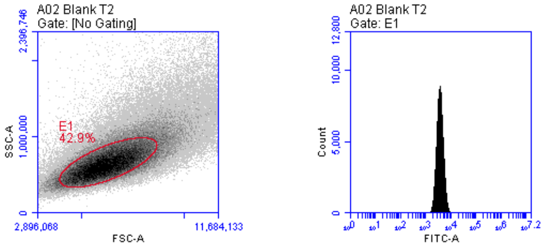

**Blank B16 - MDR cells**

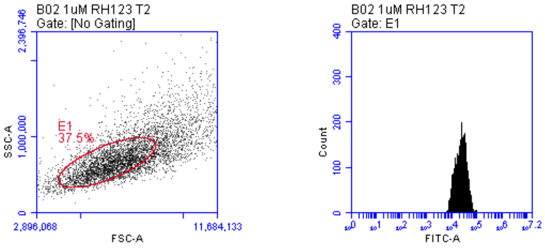

**1 μM Rhodamine 123**

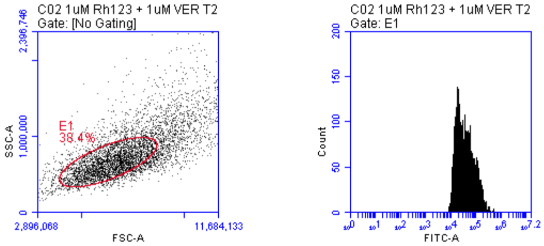

**1 μM Rhodamine 123 +**
**1 μM Verapamil**

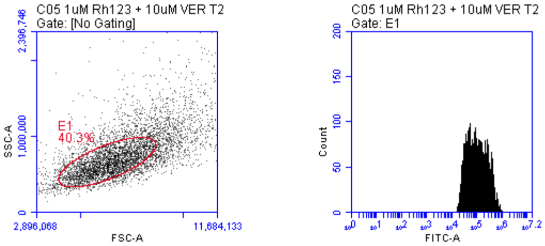

**1 μM Rhodamine 123 +**
**10 μM Verapamil**

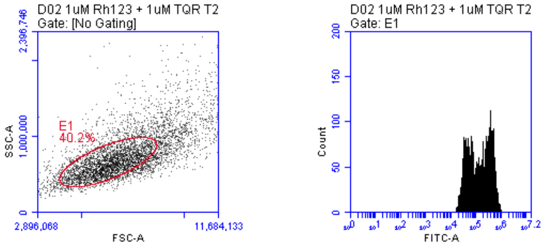

**1 μM Rhodamine 123 +**
**1 μM Tariquidar**

### Competitive Experiment with B16-MDR Cancer Cell Line
### (Raw Flow Cytometry Plots)

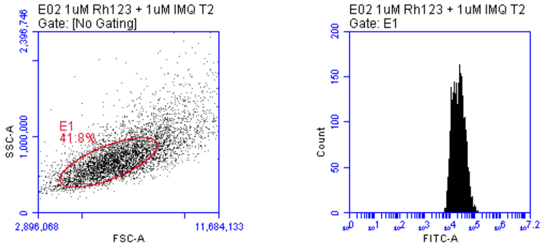

**1 µM Rhodamine 123 +
1 µM Imiquimod**

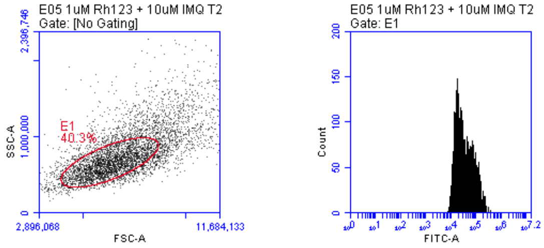

**1 µM Rhodamine 123 +
10µM Imiquimod**

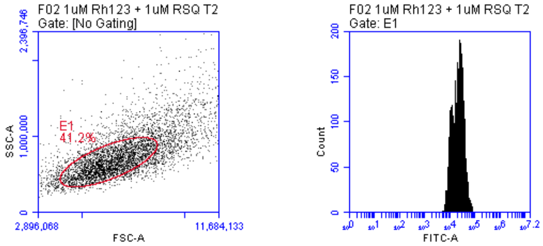

**1 µM Rhodamine 123 +
1 µM Resiquimod**

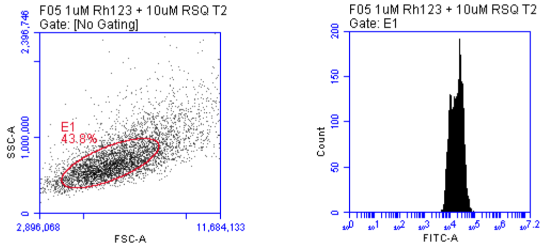

**1 µM Rhodamine 123 +
10 µM Resiquimod**

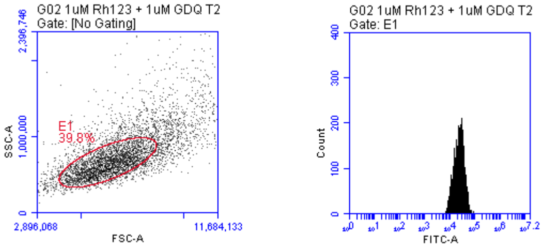

**1 µM Rhodamine 123 +
1 µM Gardiquimod**

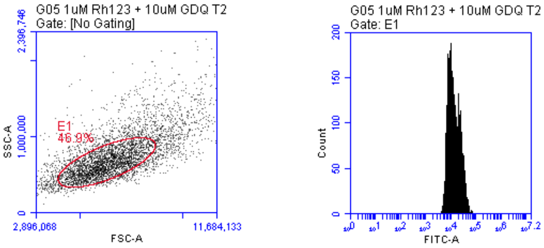

**1 µM Rhodamine 123 +
10 µM Gardiquimod**

### Competition Experiment with 4T1-MDR Cancer Cell Line
### (Raw Flow Cytometry Plots)

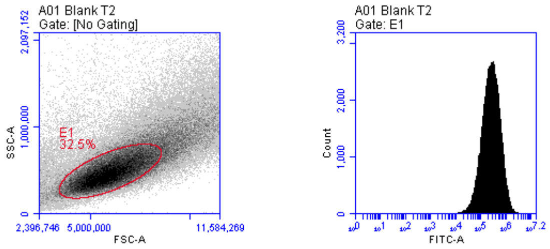

**Blank 4T1 MDR cells**

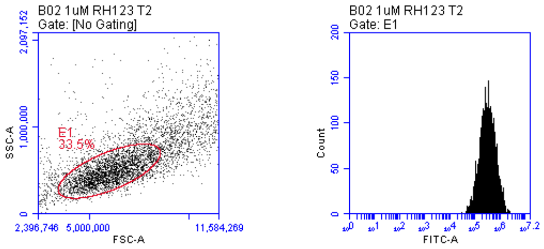

**1 µM Rhodamine 123**

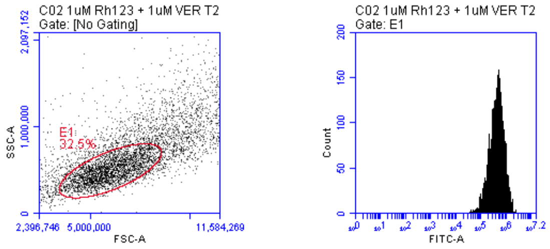

**1 µM Rhodamine 123 +**
**1 µM Verapamil**

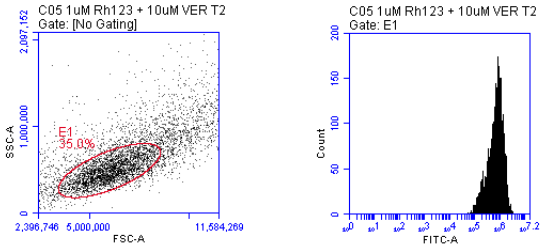

**1 µM Rhodamine 123 +**
**10 µM Verapamil**

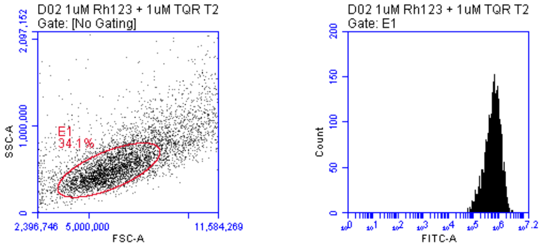

**1 µM Rhodamine 123 +**
**1 µM Tariquidar**

### Competition Experiment with 4T1 MDR Cancer Cell Line
### (Raw Flow Cytometry Plots)

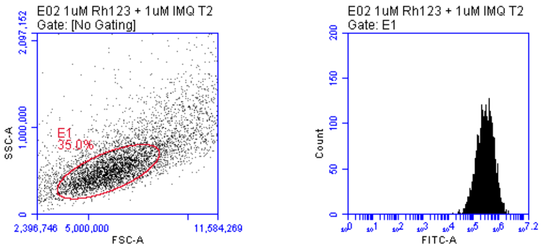

**1 µM Rhodamine 123 +**
**1 µM Imiquimod**

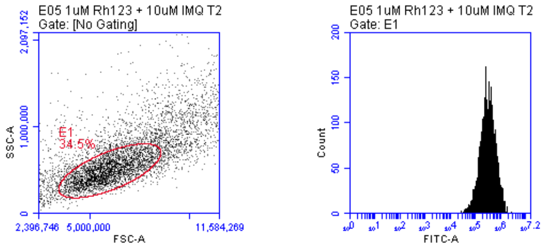

**1 µM Rhodamine 123 +**
**10 µM Imiquimod**

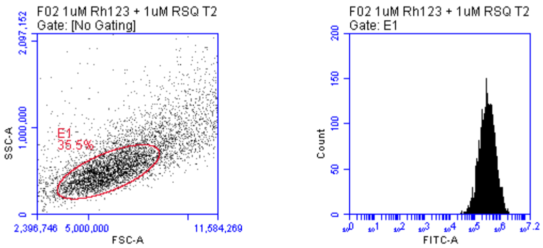

**1 µM Rhodamine 123 +**
**1 µM Resiquimod**

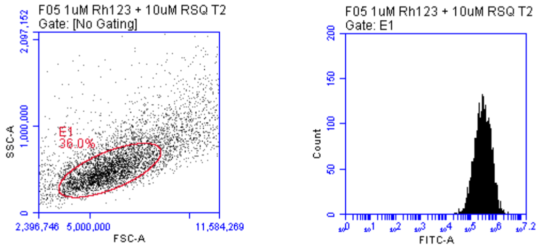

**1 µM Rhodamine 123 +**
**10 µM Resiquimod**

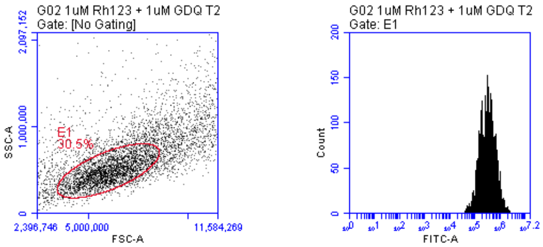

**1 µM Rhodamine 123 +**
**1 µM Gardiquimod**

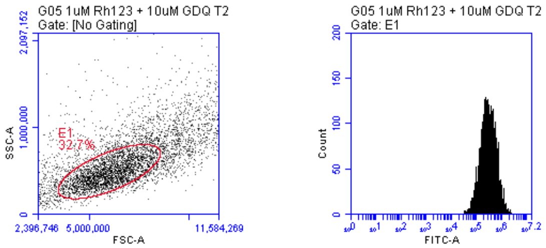

**1 µM Rhodamine 123 +**
**10 µM Gardiquimod**

### Competitive Experiment with AT3B-1 MDR Cancer Cell Line
### (Raw Flow Cytometry Plots)

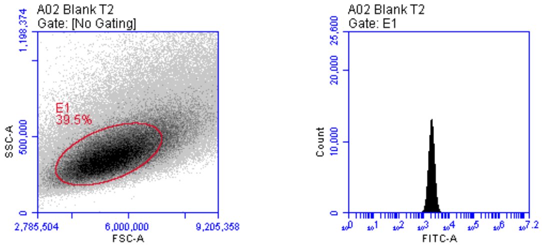

**Blank AT3B-1 MDR cells**

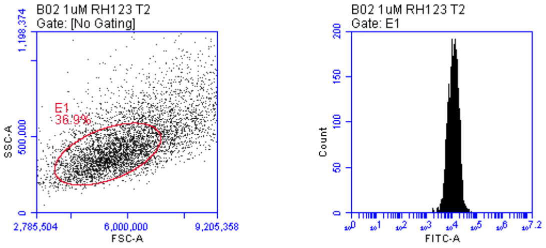

**1 µM Rhodamine 123**

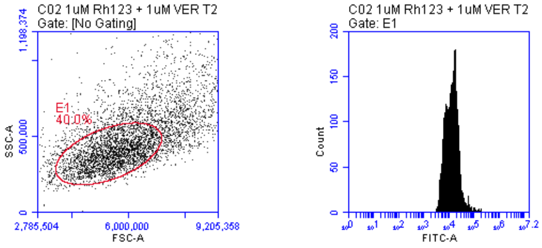

**1 µM Rhodamine 123 +**
**1 µM Verapamil**

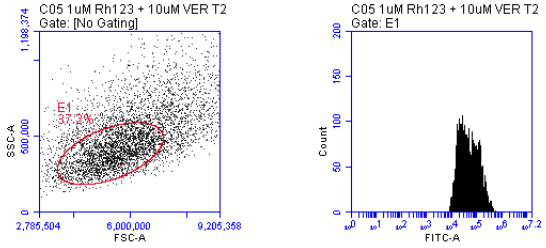

**1 µM Rhodamine 123 +**
**10 µM Verapamil**

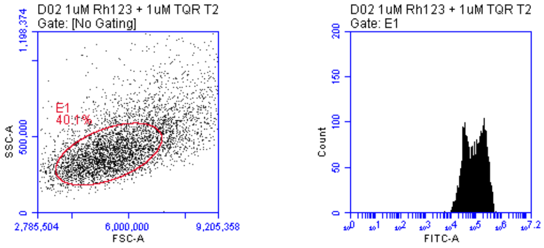

**1 µM Rhodamine 123 +**
**1 µM Tariquidar**

#### Competitive Experiment with AT3B-1 MDR Cancer Cell Line (Raw Flow Cytometry Plots)

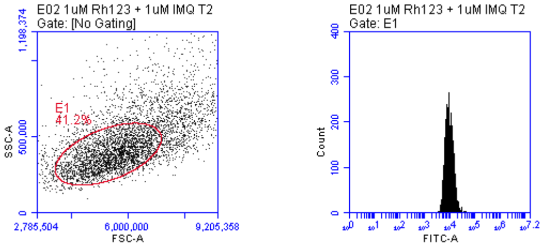

**1 µM Rhodamine 123 + 1 µM Imiquimod**

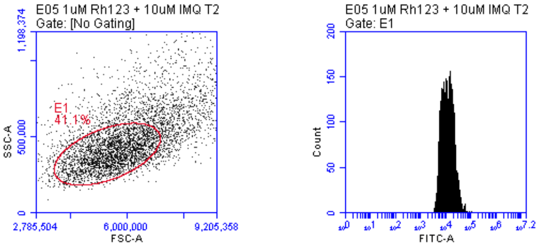

**1 µM Rhodamine 123 + 10 µM Imiquimod**

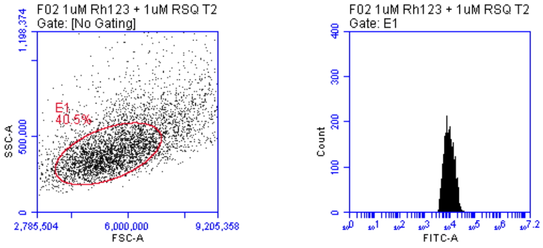

**1 µM Rhodamine 123 + 1 µM Resiquimod**

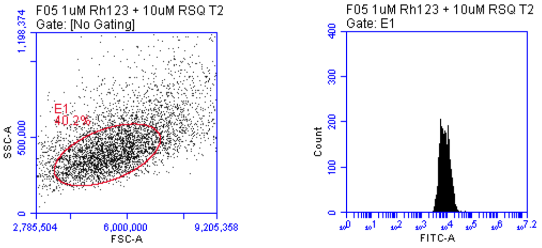

**1 µM Rhodamine 123 + 10 µM Resiquimod**

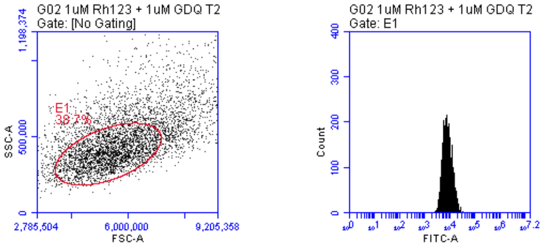

**1 µM Rhodamine 123 + 1 µM Gardiquimod**

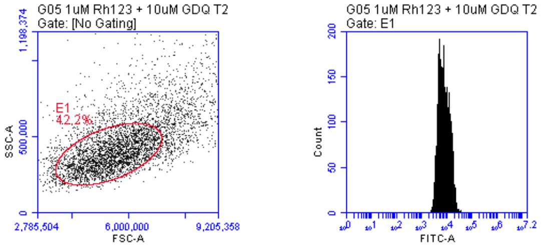

**1 µM Rhodamine 123 + 10 µM Gardiquimod**

### Synthetic Procedure of Gardiquimod

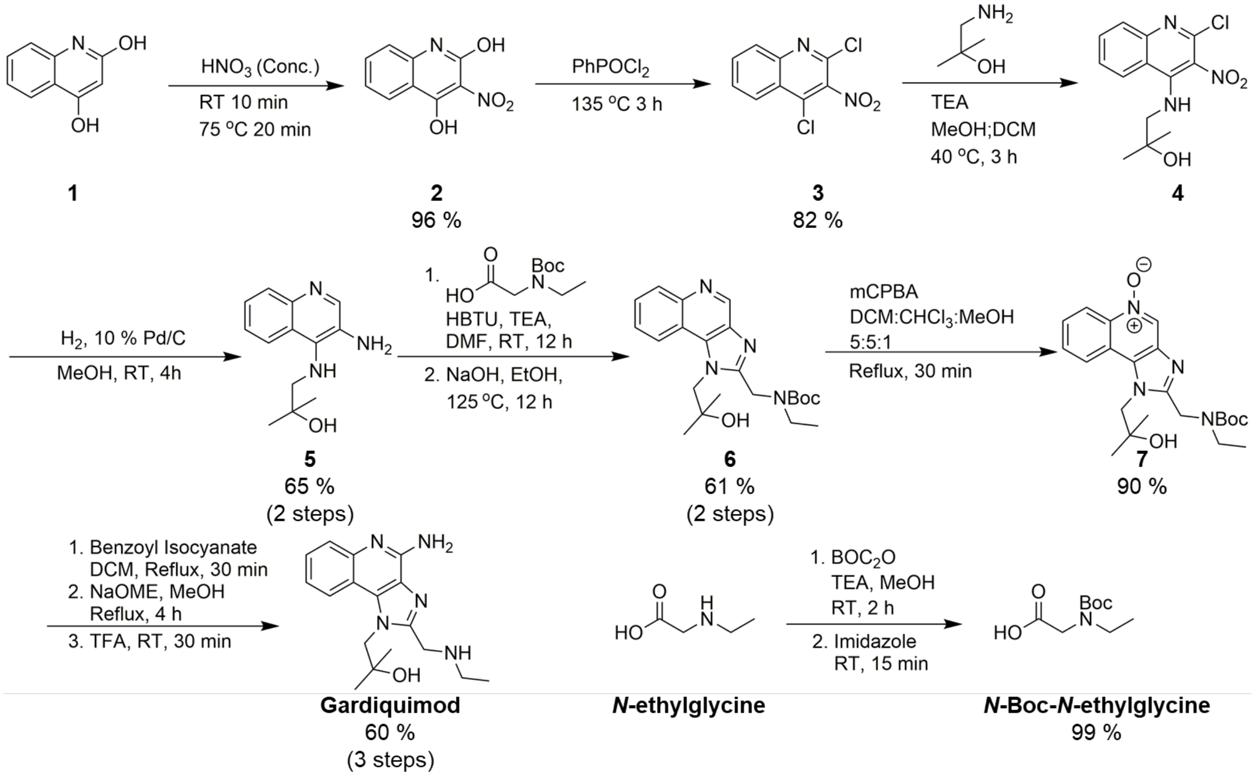

**Scheme 1:** Synthetic route used to prepare Gardiquimod in 17 % linear yield from 2,4-dihydroxyquinoline.

Although commercially available, Gardiquimod was synthesized to enable comparison of performance with future derivatives planned on the imidazole portion of the molecule. Here, 2,4-dihydroxy quinoline (**1**) was chosen as an inexpensive and readily available starting material with the synthetic route used to synthesize Gardiquimod adapted from the methods reported by Taguchi(2) for compounds **4** and **5;** Gerster(3) for compound (**3**) and David(4,5) for compounds **2, 7**, and Gardiquimod.

Briefly, 2,4-dihydroxyquinoline (**1**) is nitrated in neat concentrated nitric acid to yield 2,4-dihydroxy-3-nitroquinoline (**2**) which can be precipitated in ice cold $H_2O$. Compound (**2**) is then heated in neat phenylphosphonic dichloride at 135 ˚C to yield 2,4-dichloro-3-nitroquinoline (**3**) which can be recovered following precipitation in ice cold $H_2O$, trituration in benzene and lyophilization of the organic filtrate. Compound (**3**) undergoes nucleophilic aromatic substitution under basic conditions with 1-amino-2-methylpropan-2-ol to yield the secondary amine (**4**). The reaction preferentially occurs at the 4 position, however, and diamine resulting from displacement of both chlorides can be separated by recrystallization in benzene.

At this point, the nitro group is reduced by hydrogenation. The 2-chloro substituent is reduced with palladium catalyst and diamine (**5**) is obtained in good purity following removal of catalyst and evaporation of solvent. *N*-Boc-*N*-ethylglycine is activated with

HBTU and reacted with diamine (**5**), after 12 h solvent is removed, and the resultant amide refluxed under basic conditions to aromatize the system to accomplish the Traube reaction yielding imidazoquinoline (**6**) which is purified by column chromatography with isocratic Ethyl acetate with 1% TEA.

To install the aniline on the 4 position, imidazoquinoline (**6**) is oxidized with mCPBA to yield $N^5$-oxide (**7**). When the *N*-oxide is treated with benzoyl isocyanate in DCM at reflux the benzoyl protected aniline is provided. The benzoyl group is removed via methanolysis to liberate the anilino group and the Boc protecting group is removed with standard trifluoroacetic acid deprotection, and the resultant crude residue freebased with triethylamine and purified via a reverse phase C18 flash column using a gradient of 10 to 90 % acetonitrile in water to yield Gardiquimod in 17% linear yield over 10 steps from 2,4-dihydroxy quinoline.

#### Characterization of Gardiquimod

Gardiquimod

$^1$H NMR (600 MHz, DMSO-*d6*) δ 8.21 (d, *J* = 8.2 Hz, 1H; Ar-H), 7.59 (d, *J* = 8.3 Hz, 1H; Ar-H), 7.39 (t, *J* = 7.6 Hz, 1H, Ar-H), 7.21 (t, *J* = 7.5 Hz, 1H, Ar-H), 6.52 (s, 2H; $NH_2$), 4.70 (s, 2H; $CH_2$), 4.10 (s, 2H; $CH_2$), 2.60 (q, *J* = 7.1, 6.5 Hz, 2H; $CH_2$), 1.16 (s, 6H; 2 $CH_3$), 1.03 (t, *J* = 7.2 Hz, 3H; $CH_3$) ppm; $^{13}$C NMR (151 MHz, DMSO-*d6*) δ = 152.8, 151.9, 145.1, 133.9, 126.3, 126.3, 126.0, 121.0, 120.5, 115.4, 70.2, 55.2, 45.6, 43.1, 27.8, 14.7 ppm; UV/VIS(Acetonitrile): $\lambda_{Max}$(ε)= 228 nm (35400), $\lambda_2$(ε)= 247 nm (34700); IR(ATR): 3301 (br, m), 2984 (m), 1673 (s), 1200 (s), 1131 (s), 721 (w); HRMS(MALDI): m/z calcd for $C_{17}H_{24}N_5O$: 314.1981 $[M+H]^+$, observed 314.19733. (Δ=2.45 ppm).

$^1H$ NMR Spectra of Gardiquimod

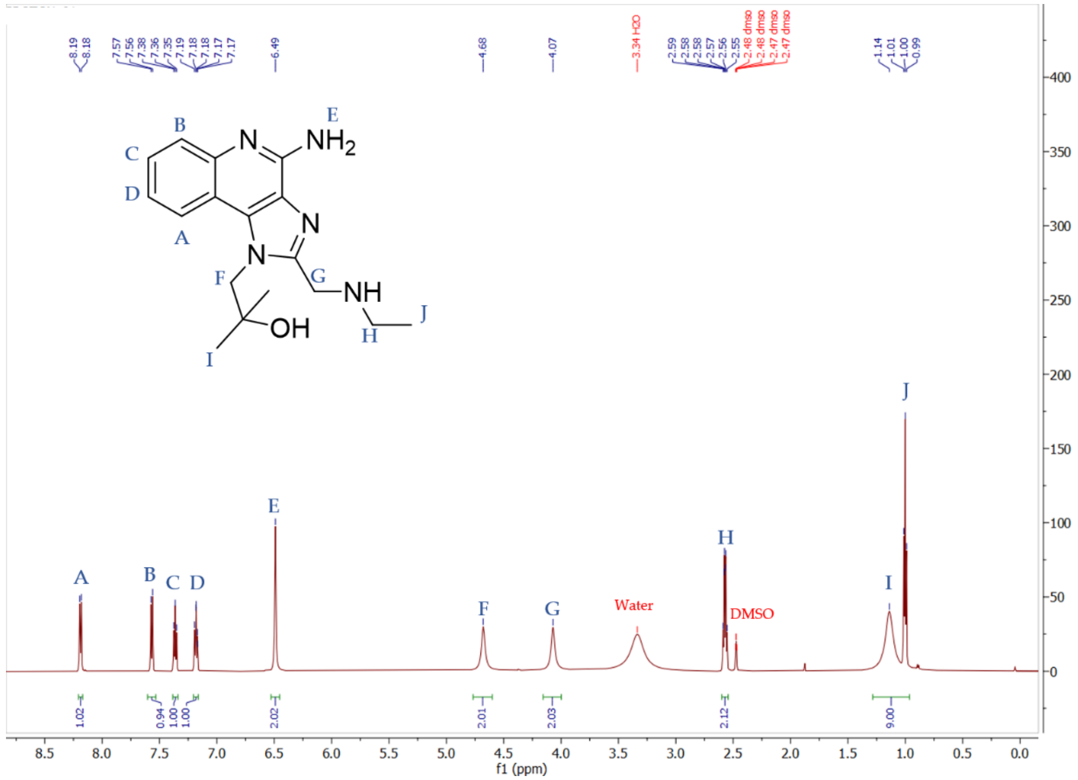

$^{13}C$ Spectra of Gardiquimod

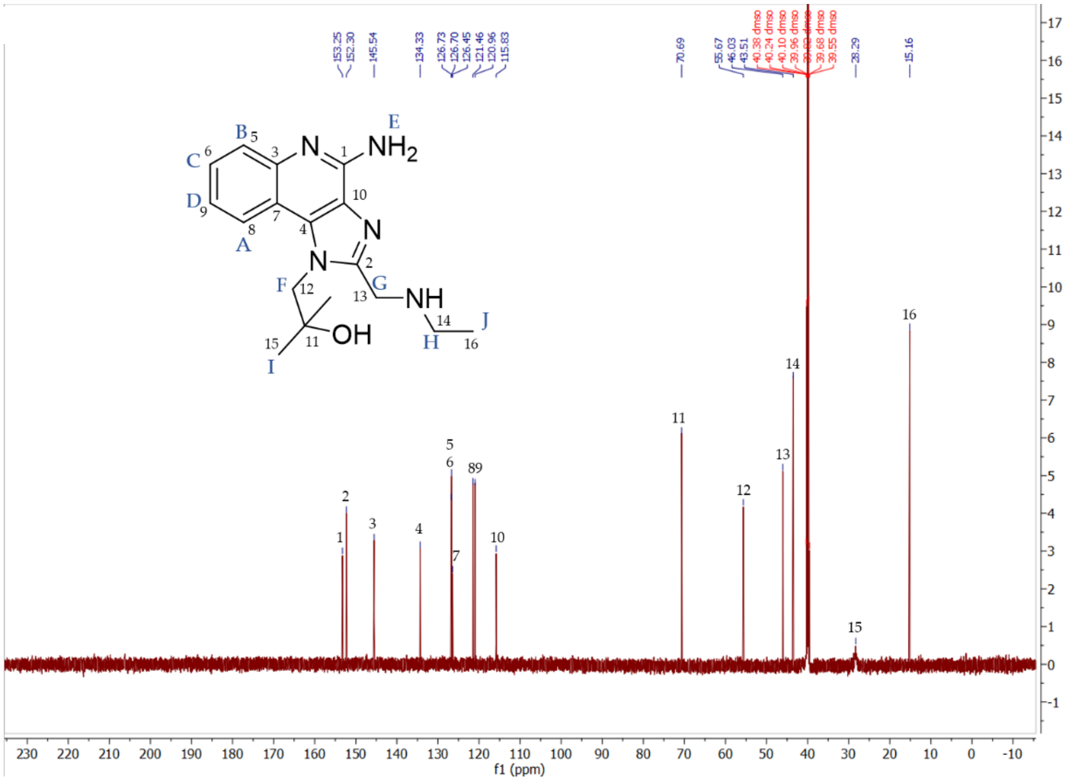

COSY Spectra of Gardiquimod

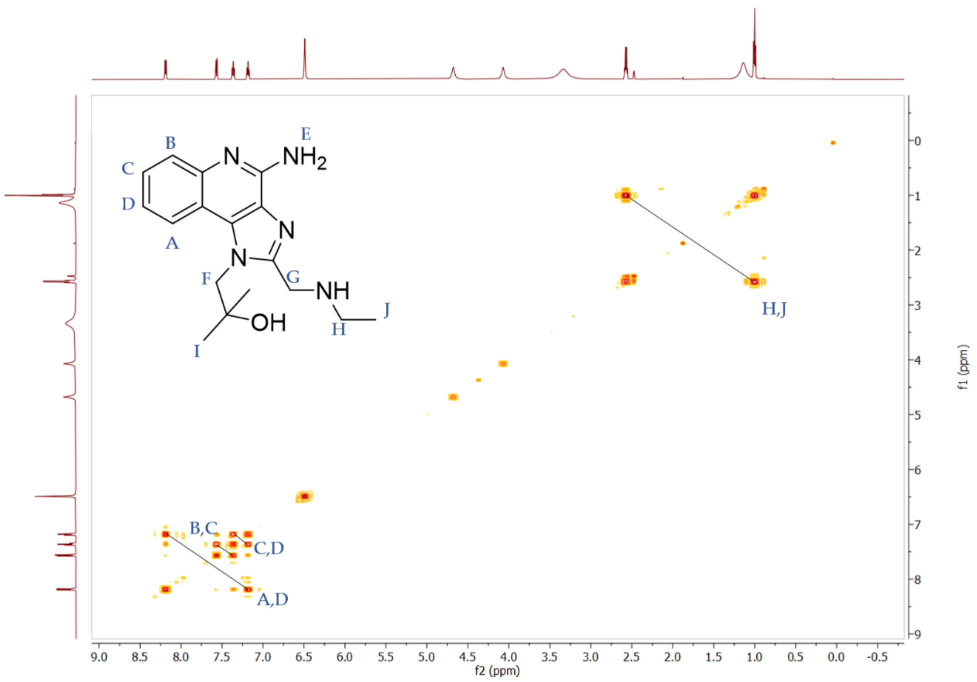

HSQC Spectra of Gardiquimod

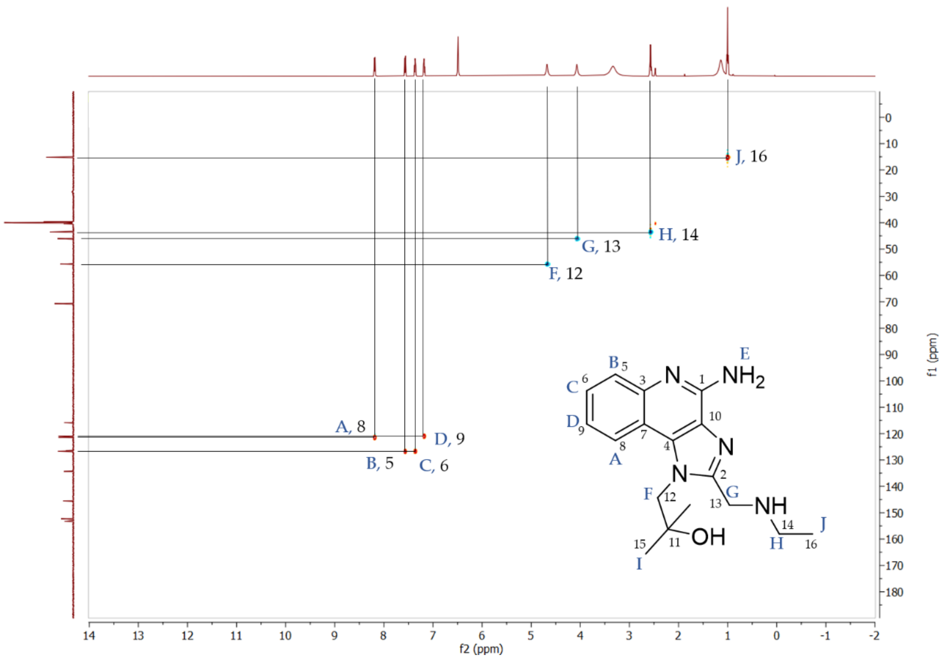

HSQC with enhanced intensity to show signal for I

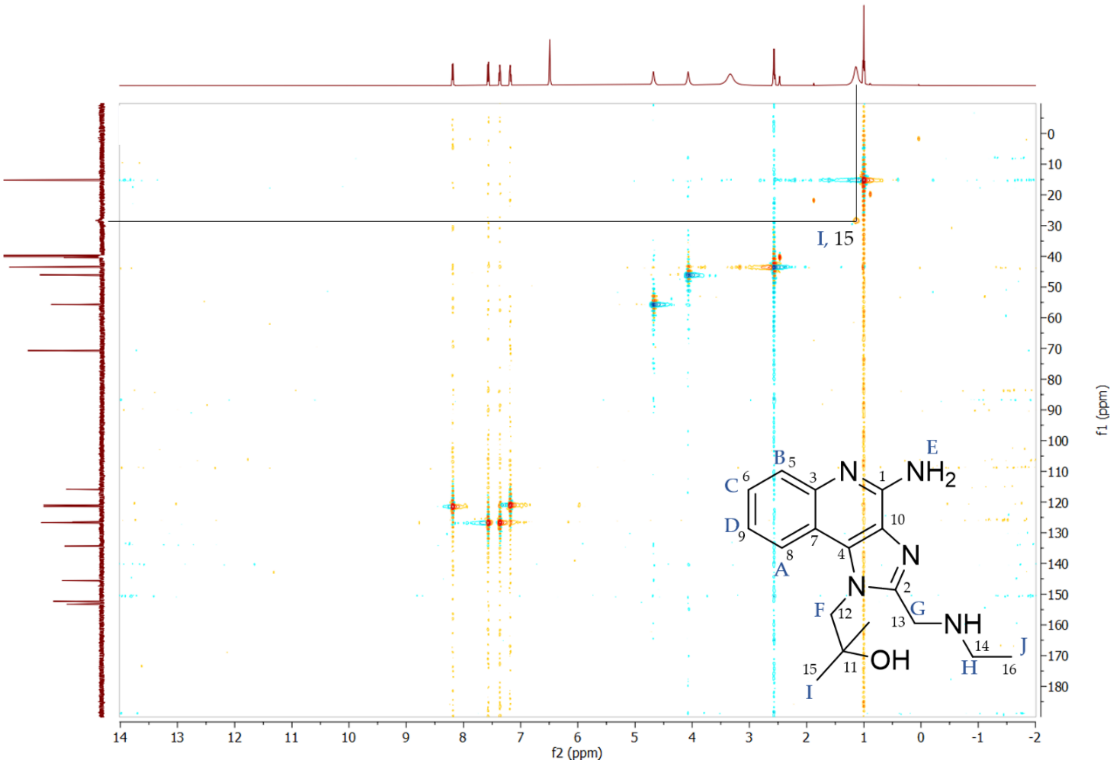

$^{1}$H NMR Spectra for Numbered Intermediates **2**-**7**

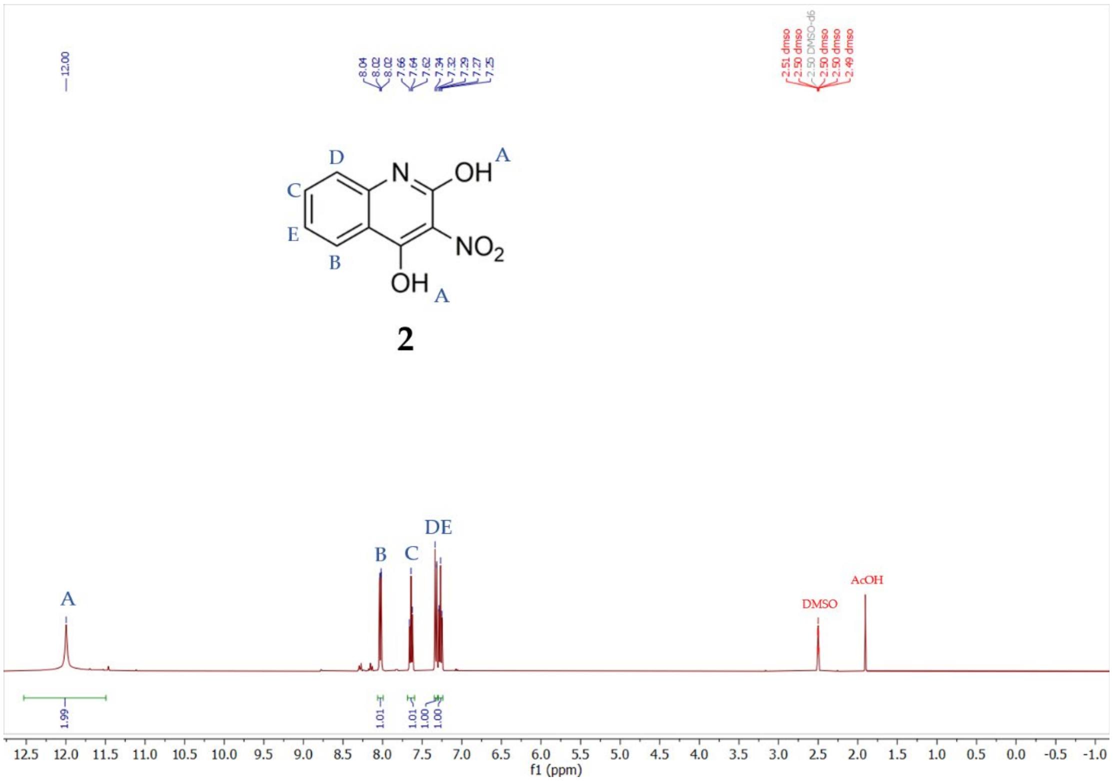

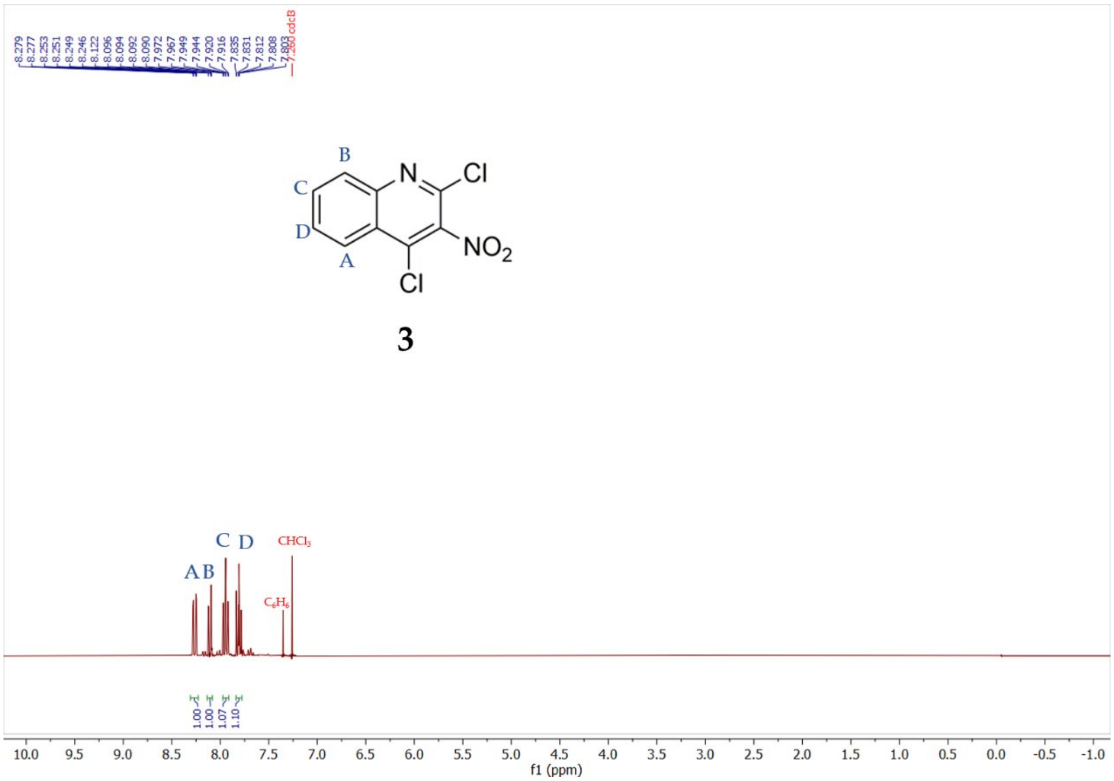

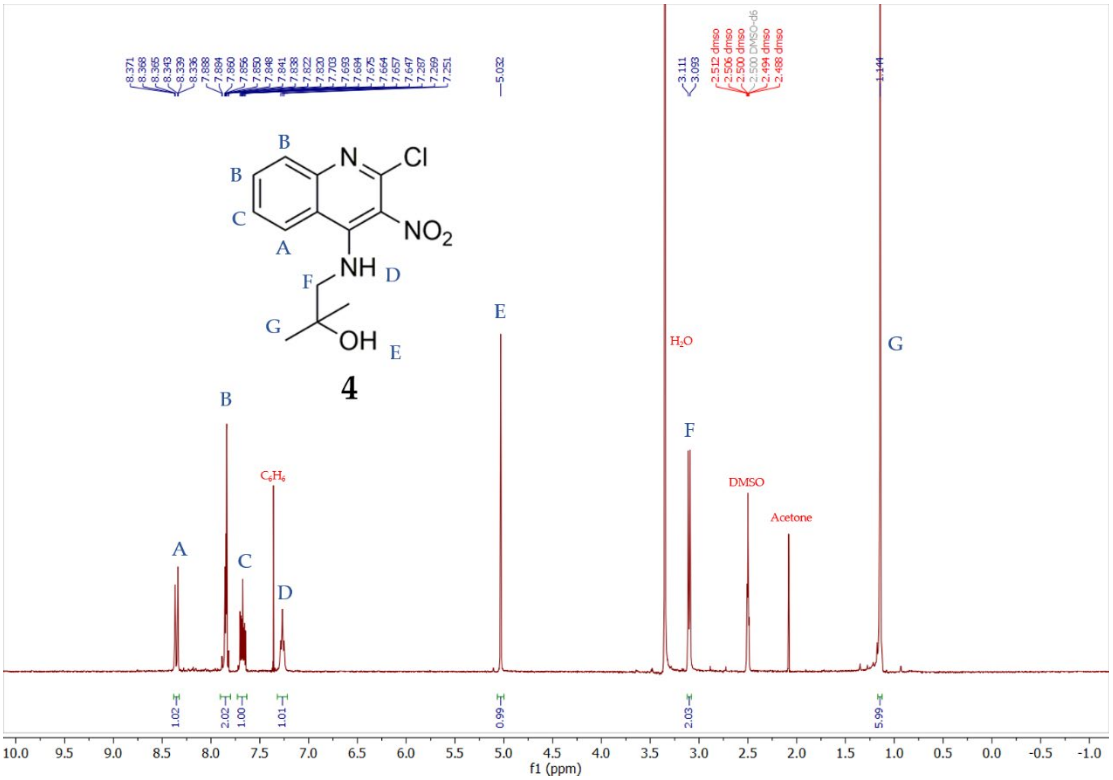
8.371
8.368
8.365
8.343
8.339
8.336
7.888
7.884
7.860
7.856
7.850
7.848
7.841
7.838
7.822
7.820
7.703
7.693
7.684
7.675
7.664
7.657
7.647
7.287
7.269
7.251
5.032
3.111
3.093
2.512 dmso
2.506 dmso
2.500 dmso
2.500 DMSO-d6
2.494 dmso
2.488 dmso
1.144
B
B
C
A
N
Cl
NO2
NH
D
F
G
OH
E
4
H2O
C6H6
DMSO
Acetone
1.02
2.02
1.00
1.01
0.99
2.03
5.99
f1 (ppm)

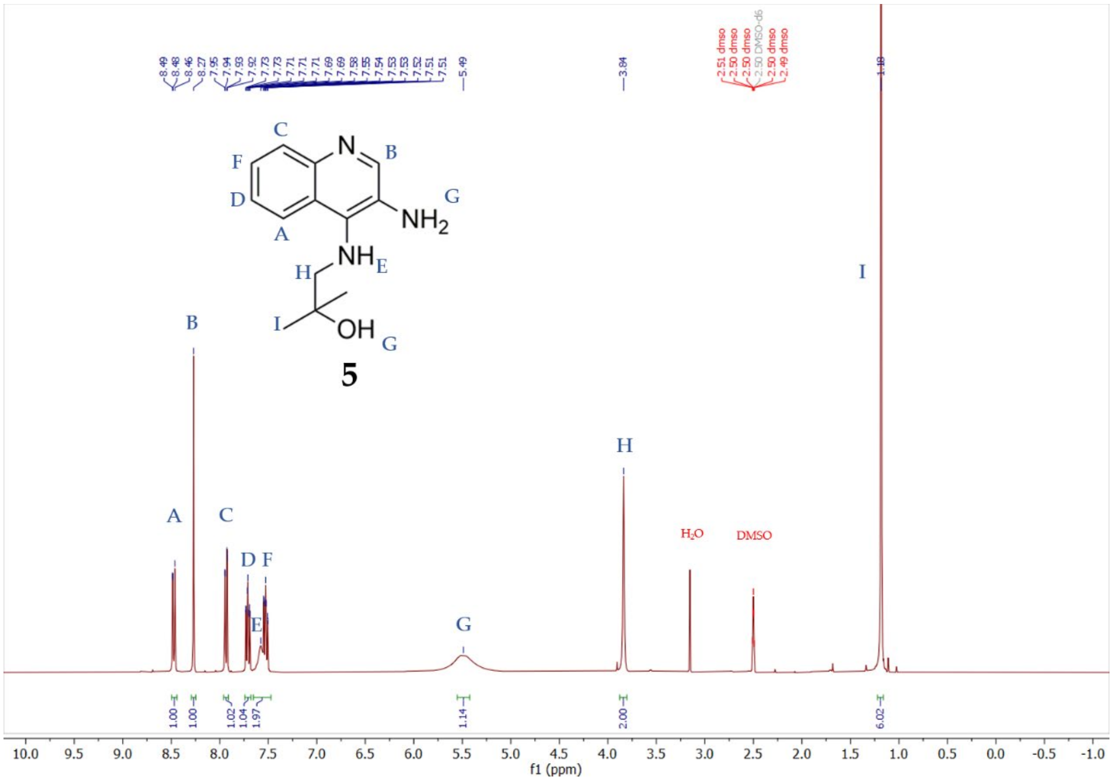
8.49
8.48
8.46
8.27
7.95
7.94
7.93
7.92
7.73
7.73
7.71
7.71
7.69
7.69
7.58
7.55
7.54
7.53
7.53
7.52
7.51
5.49
3.84
2.51 dmso
2.50 dmso
2.50 DMSO-d6
2.50 dmso
2.49 dmso
1.18
C
N
B
F
D
G
NH2
A
NH
E
H
I
OH
G
5
H2O
DMSO
1.00
1.00
1.02
1.04
1.97
1.14
2.00
6.02
f1 (ppm)

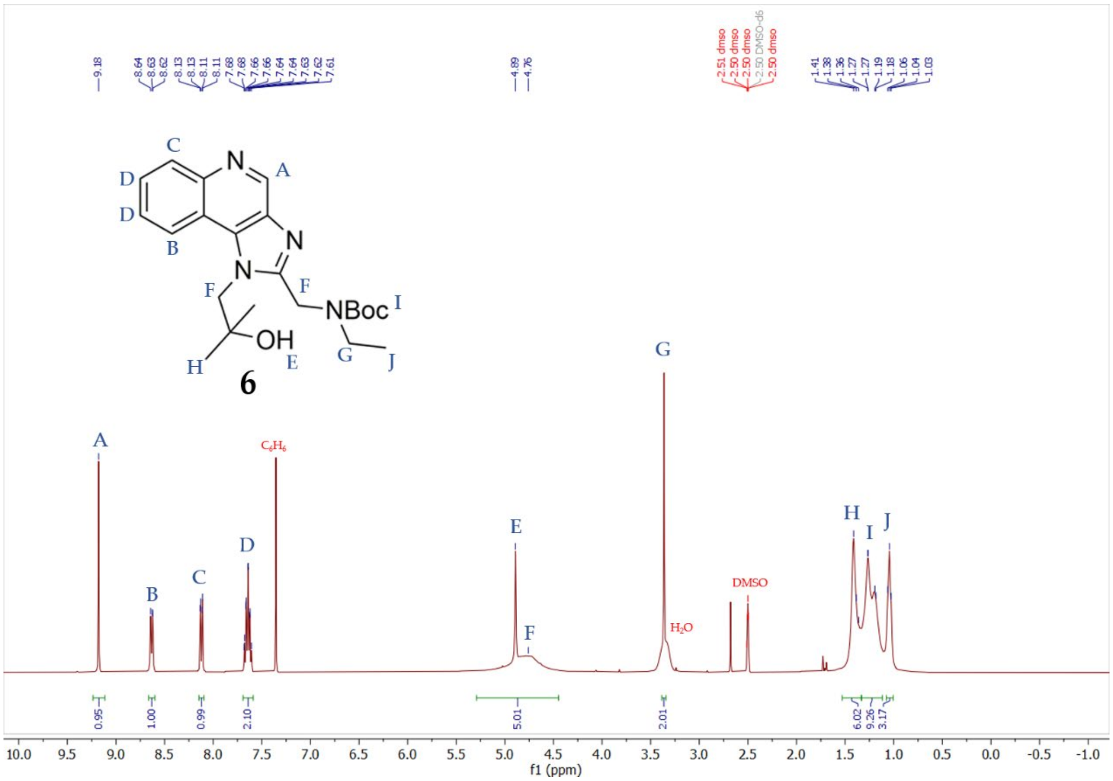

9.18
8.64 8.63 8.62 8.13 8.13 8.11 8.11 7.68 7.66 7.66 7.64 7.64 7.63 7.62 7.61
4.89 4.76
2.51 dmso 2.50 dmso 2.50 dmso 2.50 DMSO-d6 2.50 dmso
1.41 1.38 1.36 1.27 1.27 1.19 1.18 1.06 1.04 1.03
6
A B C D E F G H I J
C6H6
H2O
DMSO
0.95 1.00 0.99 2.10 5.01 2.01 6.02 9.26 3.17
f1 (ppm)

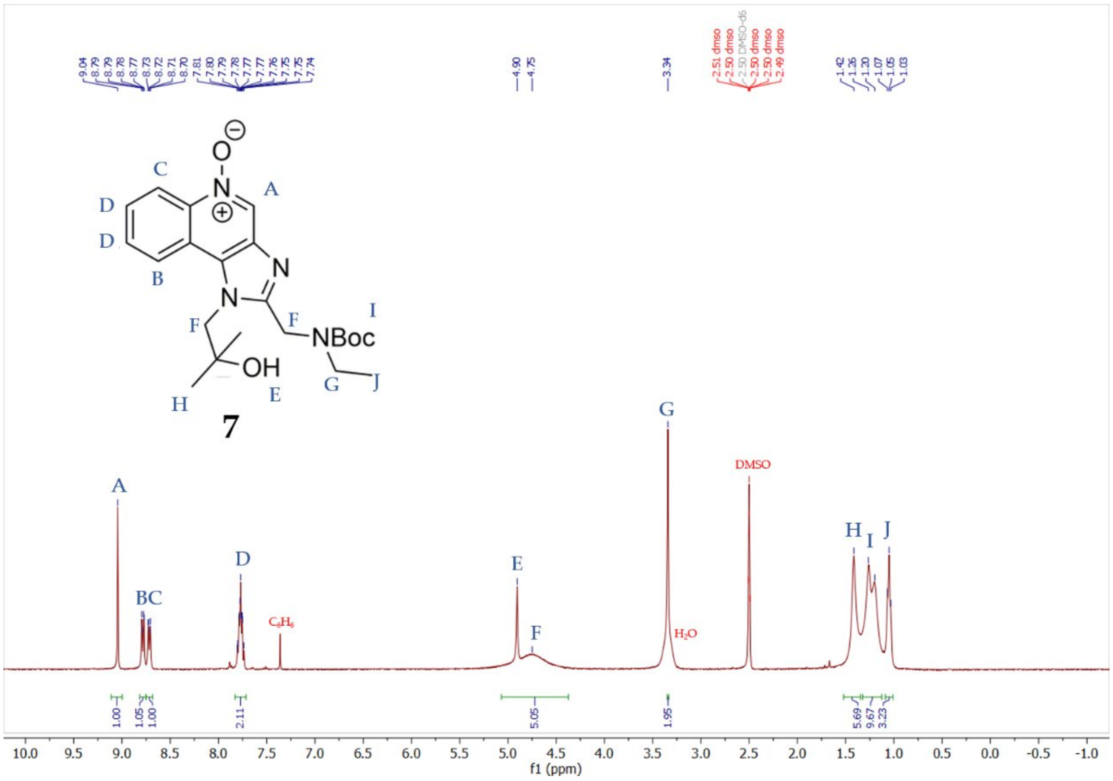

9.04 8.79 8.79 8.78 8.77 8.73 8.72 8.71 8.70 7.81 7.80 7.79 7.78 7.77 7.77 7.76 7.75 7.75 7.74
4.90 4.75
3.34
2.51 dmso 2.50 dmso 2.50 DMSO-d6 2.50 dmso 2.50 dmso 2.49 dmso
1.42 1.36 1.20 1.07 1.05 1.03
7
A BC D E F G H I J
C6H6
H2O
DMSO
1.00 1.05 1.00 2.11 5.05 1.95 5.69 9.67 3.23
f1 (ppm)

#### References

1. Replogle-Schwab TS, Schwab ED, Pienta KJ. Development of doxorubicin resistant rat prostate cancer cell lines. Anticancer Res. 1997;17:4535–8.

2. Fujita Y, Hirai K, Nishida K, Taguchi H. 6-(4-Amino-2-butyl-imidazoquinolyl)-norleucine: Toll-like receptor 7 and 8 agonist amino acid for self-adjuvanting peptide vaccine. Amino Acids. 2016;48:1319–29.

3. Gerster JF, Lindstrom KJ, Miller RL, Tomai MA, Birmachu W, Bomersine SN, et al. Synthesis and Structure−Activity-Relationships of 1H-Imidazo[4,5-c]quinolines That Induce Interferon Production. J Med Chem. 2005;48:3481–91.

4. Shukla NM, Malladi SS, Mutz CA, Balakrishna R, David SA. Structure−Activity Relationships in Human Toll-Like Receptor 7-Active Imidazoquinoline Analogues. J Med Chem. 2010;53:4450–65.

5. Shukla NM, Kimbrell MR, Malladi SS, David SA. Regioisomerism-dependent TLR7 agonism and antagonism in an imidazoquinoline. Bioorganic & Medicinal Chemistry Letters. 2009;19:2211–4.
